# Devonian stromatoporoid historical collections in the Natural History Museum, London (UK): redescription, taxonomic revision and implications for stromatoporoid global paleobiogeography

**DOI:** 10.1101/2024.06.30.601411

**Authors:** Jiayuan Huang, Consuelo Sendino, Stephen Kershaw

**Author notes:** **Corresponding author:** ORCID ID: 0000-0003-1099-9076.

## Abstract

Devonian stromatoporoid collections in the Natural History Museum, London (UK) have been deposited for over 100 years. The characteristics and systematic position of these specimens, however, have received little attention. In this study, a total of 307 Devonian stromatoporoid specimens comprising material documented by Nicholson (1886–1892) from the UK, Germany, United States, and Canada, plus specimens described by Ripper (1933, 1937a, b, c) from Australia, were re-examined. Overall, 50 species belonging to 29 genera were systematically redescribed based on recent progress, mainly including *Actinostroma*, *Petridiostroma*, *Stictostroma*, *Pseudotrupetostroma*, *Parallelopora*. Three-dimensional reconstructions of the type genus are illustrated following the earlier works of Stearn (1966) and Cockbain (1984). The reconstructed skeletons reveal stromatoporoid architectural patterns, crucial for enhancing understanding and revision of stromatoporoid identification. This study underscores the significance of three-dimensional reconstruction in taxonomic research on stromatoporoid. The NHMUK material is combined with data from publications and the Paleobiology Database (PBDB) to perform a network analysis of the global occurrence of Devonian stromatoporoids at the generic level; this reveals a close relationship of global stromatoporoid fauna during the Early Devonian, indicating a widespread distribution, despite this interval being regarded as a time of global stromatoporoid contraction. The Middle Devonian assemblage shows a much higher cosmopolitan occurrence in the context of the subsequent Eifelian-Givetian global stromatoporoid proliferation, consistent with the known pattern from other studies of Middle Devonian stromatoporoids. Overall, the NHMUK collections are a valuable resource to help understand the global occurrence of Devonian stromatoporoids.

## 1. Introduction

The Devonian Period witnessed exceptional proliferation of stromatoporoids during their ca. 100 million-year evolutionary history, reaching their highest biodiversity in the Eifelian Stage (Stearn, 2015a), and during the Givetian-Frasnian Stages showing the highest abundance over global carbonate platforms manifested by reef occurrences (Copper, 2002). The extensive presence of Devonian stromatoporoids has attracted attention in taxonomy over the past two centuries, particularly in regions as western Europe (Goldfuss, 1825; Bargatzky, 1881; Nicholson, 1886–1892; Lecompte, 1951–1952), Russia (Yavorsky, 1931, 1955–1967), Australia (Ripper, 1933, 1937a, b, c; Cockbain, 1984; Cook, 1999), North America (Parks, 1936; Galloway and St. Jean, 1957; Stearn, 1963; Stock, 2022), the Middle East (Mistiaen, 1985; Mistiaen and Gholamalian, 2000), and South China (Yang and Dong, 1979; Dong, 2001; Huang et al., 2022).

As research progressed and techniques improved, the illustration of fossil skeletons changed accordingly. In the 19th century, stromatoporoid workers used mainly hand-drawn sketches to depict two- and three-dimensional fossil specimens (Goldfuss, 1826; Bargatzky, 1881; Nicholson, 1886). Then, photos taken by microscope were applied to visualize two-dimensional skeletons (Lecompte, 1951, 1952; Kaźmierczak, 1971; Salerno, 2008). The three-dimensional reconstruction of detailed skeletal structures, pioneered by Stearn (1966) and Cockbain (1984), offers a more intuitive method for revealing growth patterns and skeleton arrangements compared to traditional two-dimensional fossil plates and depictions. This approach has provided valuable insights into the classification and revisions of stromatoporoids (Huang et al., 2024).

Since Goldfuss (1825) established the first genus *Stromatopora* in the Middle Devonian of Germany, over 100 Devonian stromatoporoid genera and numerous species were reported by global workers (Steran, 2015b). Although great progress has been achieved regarding Devonian stromatoporoid taxonomy, problems emerged and remain unsolved. Many genera erected based on local specimens are confused and outdated. Therefore, revisions of Devonian stromatoporoids have been ongoing during the last three decades (Stearn, 1993; May, 1999; Stearn et al., 1999; Stearn, 2015b; Huang et al., 2024). In this paper, we focus on revisions of a major collection of Devonian stromatoporoids deposited in the Natural History Museum, London (UK). Most of these specimens were documented by Nicholson (1886–1892), with several classical genera erected (e.g., *Actinostroma*, *Hermatostroma*, *Stromatoporella*).

The aims of this study are to: 1) redescribe and re-illustrate the NHMUK (The Natural History Museum, London) collections in the modern context of stromatoporoid taxonomy; 2) thus, revise problematic taxa based on recent advances; 3) elaborate the skeletal structure of different types of stromatoporoids by three-dimensional reconstructions; and 4) discuss the global Devonian stromatoporoid paleobiogeography based on the NHMUK stromatoporoid legacy material in relation to the growing global database of Devonian stromatoporoids.

## 2. Locations, materials and methods

Thin sections of a total of 307 Devonian stromatoporoid specimens are stored at NHMUK, including specimens from Devon, UK (144), Germany (109), Australia (47), United States (5), and Canada (2), see Fig. 1. Detailed fossil occurrences (i.e., formations and localities) were not recorded at that time, but general locations are indicated on thin section labels, so these are shown on Fig. 1. Most of Nicholson’s UK material is from the Teign valley locations in Devon County, southwest England (Teignmouth, Shaldon, Bishopsteignton, Kingsteignton) in pebbles of Devonian limestones present in Permo-Triassic conglomerates that were transported from paleo-uplands to the west into the Permo-Triassic Teign valley wadi. That ancient wadi is re-exposed as a modern valley in the ria coastline of Devon and there are no *in-situ* Devonian limestone outcrops in the Teign valley. Material from Germany is from numerous locations in western Germany; a smaller number of samples are from Australia and North America.

**Figure 1.**
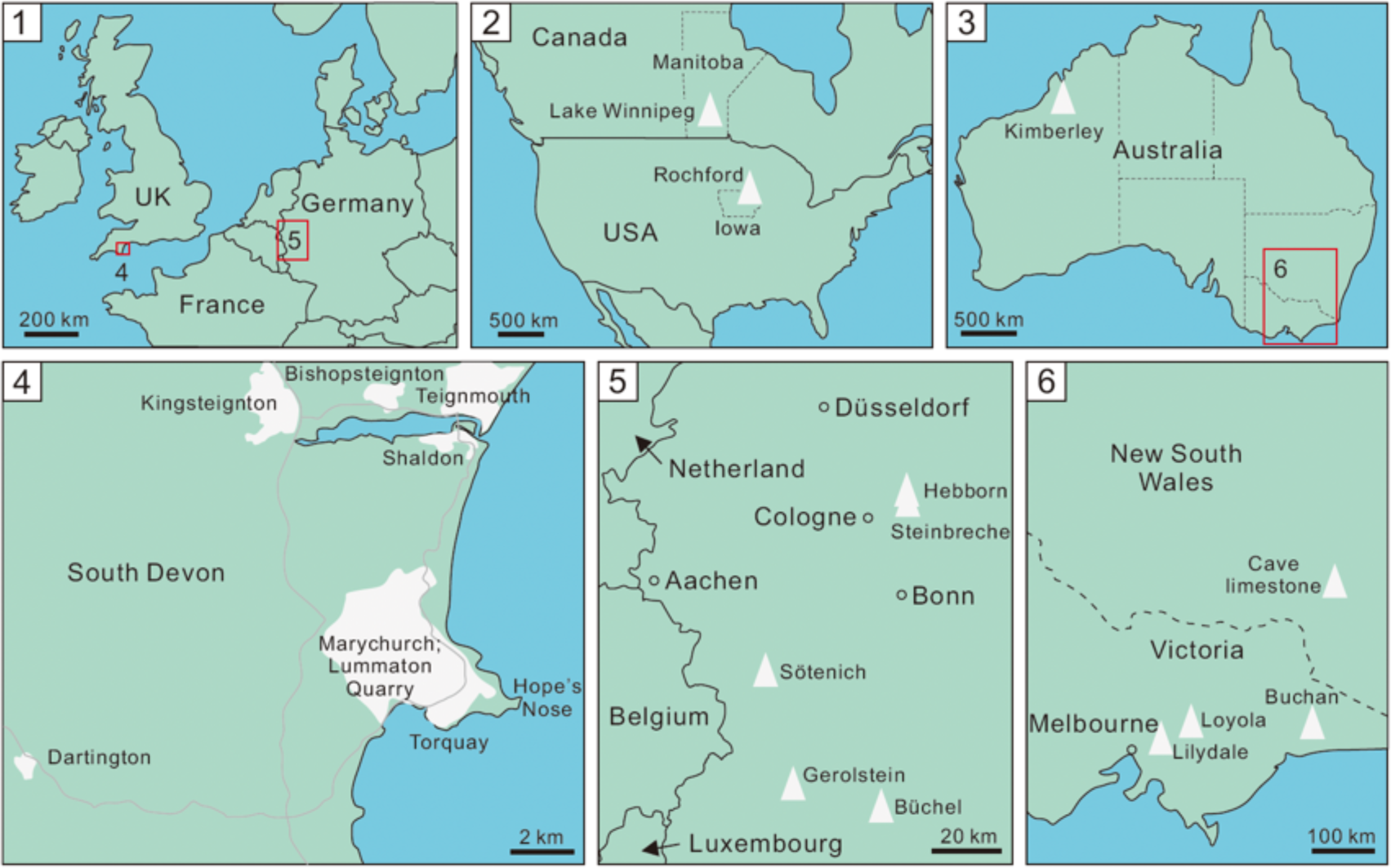
Location map of samples described in this study, based on thin section labels of NHMUK samples.

The stromatoporoid specimens exist as longitudinal and tangential thin sections. For this project, photographs were taken in normal light with a standard stereo-binocular microscope and the images were enhanced on CorelDRAW 2019. Some of the specimens are poorly preserved but taxa were largely confirmed by comparison with better preserved samples in the collection. Redescriptions are conducted mainly based on internal structure and microstructure; thin sections are of standard size (25 x 75 mm) so there is limited information available on external morphology, upper and lower surfaces, and latilaminae, although this is included where possible. All terms applied herein generally follow the usage of the revised *Treatise on Invertebrate*, *part E* (Webby and Kershaw, 2015, pp. 417–486; Stearn, 2015c, pp. 487–520; Stearn, 2015d, pp. 521–533). Three-dimensional reconstructions of representative genera are after Stearn (1966) and Cockbain (1984), see Fig. 2. Figures 3-21 illustrate the taxa studied here, described later in this paper.

**Figure 2.**
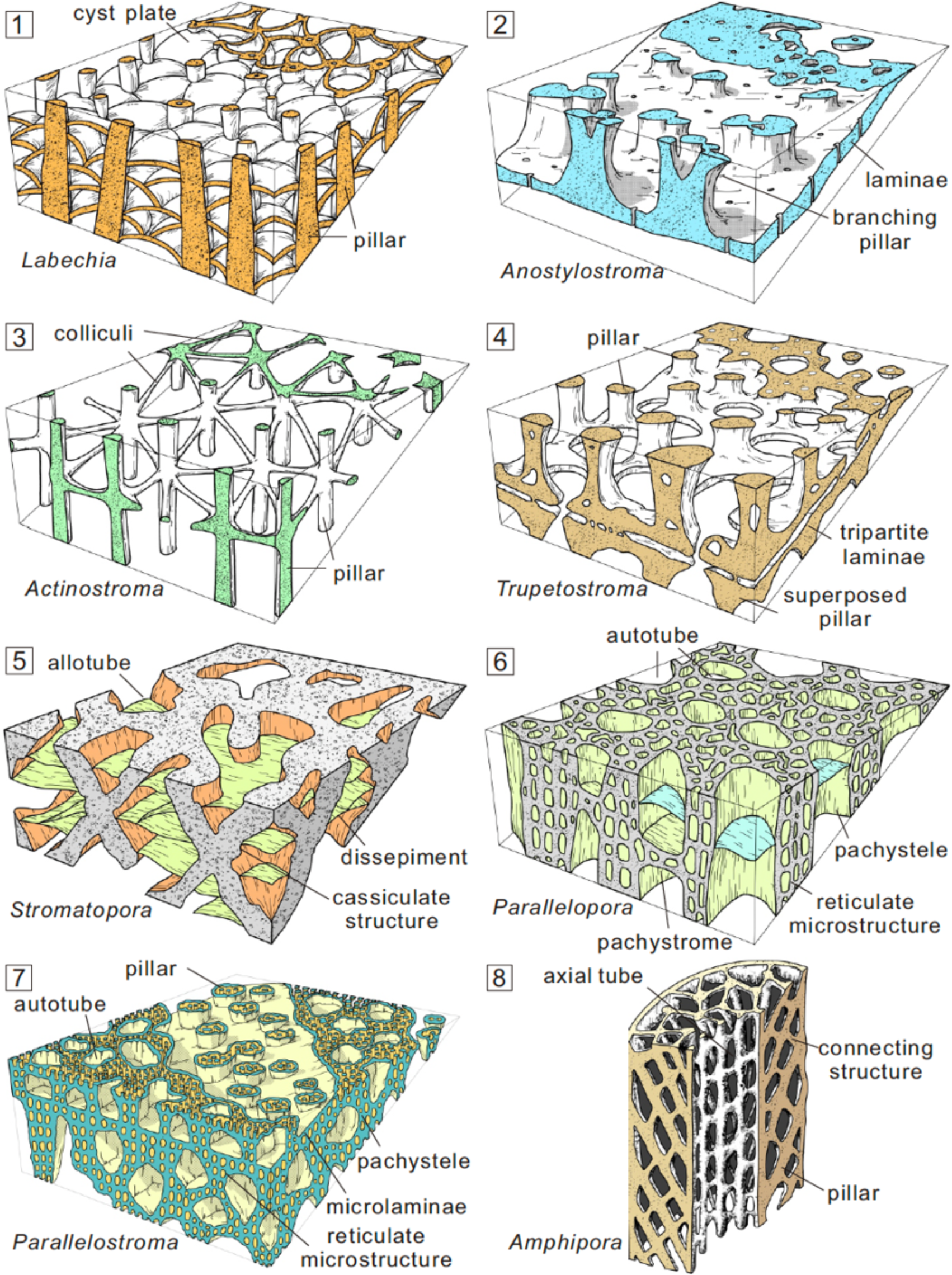
Three-dimensional reconstructions of *Labechia* (1), *Anostylostroma* (2), *Actinostroma* (3), *Trupetostroma* (4), *Stromatopora* (5), *Parallelopora* (6), *Parallelostroma* (7), and *Amphipora* (8) (modified after Stearn, 1966; Cockbain, 1984; Qie et al., 2020; Huang et al., 2024). This diagram demonstrates the variability of stromatoporoid architecture, as reflected in the taxa examined in this study.

**Figure 3.**
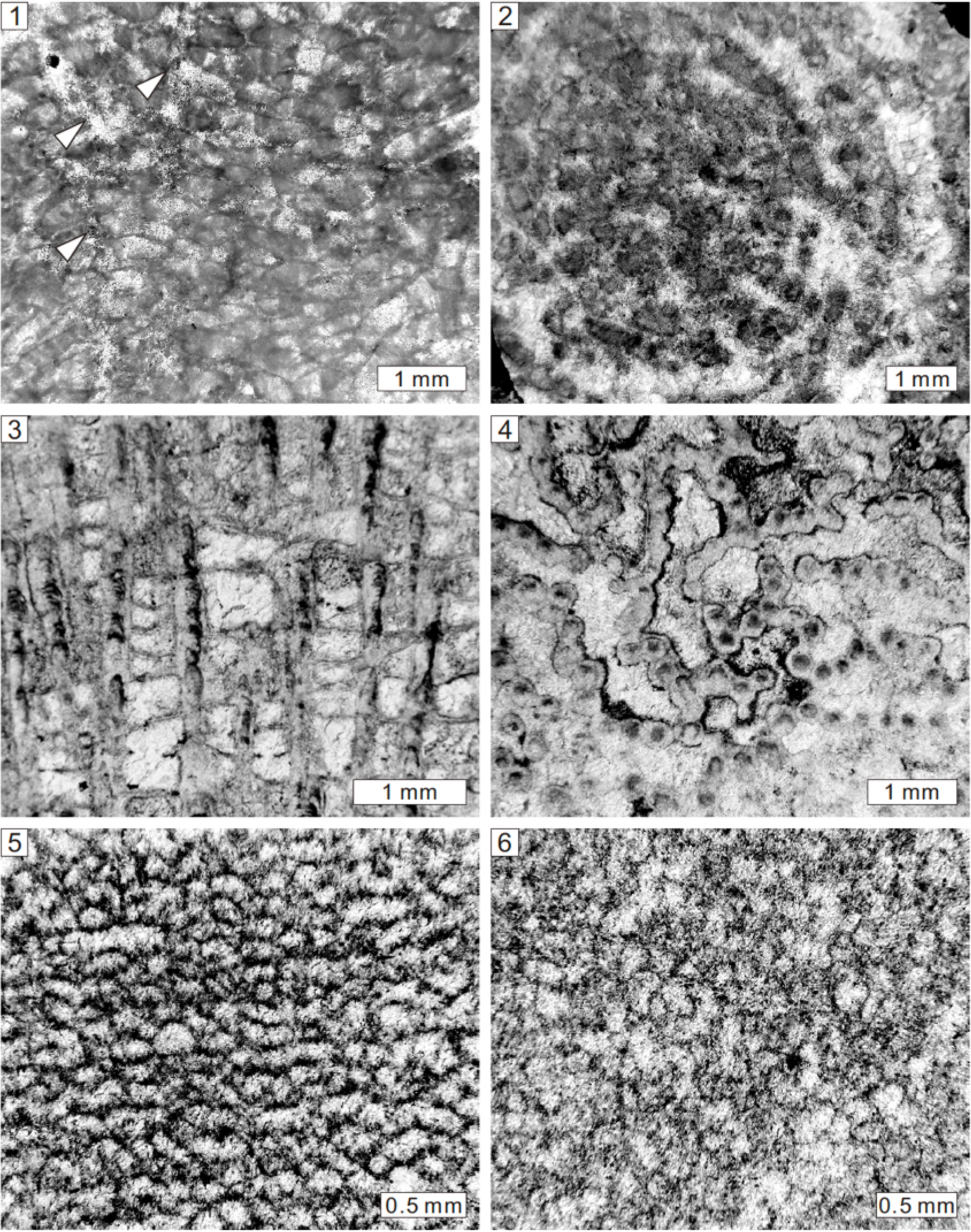
Longitudinal and tangential views of *Labechia stylophora* Nicholson, 1891 from the Middle Devonian of Shaldon, South Devon, UK, NHMUK PI P5987 (1, 2), *Labechiella serotina* (Nicholson, 1886a) from the Middle Devonian of Teignmouth, Devon, UK, NHMUK PI P5988 (3, 4), *Clathrodictyon confertum* Nicholson, 1889 from the Middle Devonian of Hope’s Nose Quarry, Torquay, South Devon, UK, NHMUK PI H3891 (5, 6). Note the unclear pillars (white arrows).

Network analysis of the Devonian stromatoporoids was carried out (Figs. 22, 23) based on the software Gephi version 0.10 (Bastian et al., 2009) wherein the occurrences of Devonian stromatoporoids are compiled as a binary dataset (i.e., continents and genera). Lines in the network diagram link a source node (continent or area) to a target node (genus). An endemic genus is connected by a single line, whereas a cosmopolitan genus is shown by several lines. The size of a source node and target node represents stromatoporoid diversity and degree of cosmopolitanism, respectively.

Modularity is applied to depict the paleobiogeographical similarity between different continents. The Force Atlas 2 option was used here for the layout of diagram. The following parameters within Gephi were used in this study: scaling, 10.0; gravity, 1.0; edge weight influence, 1.0; number of threads, 19; tolerance, 1.0; approximation, 1.2.

The Devonian stromatoporoid dataset for the network analysis was compiled from this study and references, with supplements from the PBDB. Areas and references of the dataset include north-eastern Russia (Omulevka Mountains, Omolon, Ulachan-Sis) (Khromykh, 1971, 1974, 1976), Siberia (Kuznetsk, Sayan-Altai, Salair) (Yavorsky, 1931; Rzhonsnitskaya, 1968; Khalfina, 1961, 1968), Amuria (Inner Mongolia, Mongolia) (Sharkova, 1981; Bolshakova and Ulitina, 1985; Dong, 1984, 1985), Kazakhstania (Tian Shan, Zeravshan, Uzbekistan, Tajikistan, Kyrgyzstan) (Leleshus, 1971; Lesovaya, 1970, 1971, 1972, 1982, 1986; Sapelnikov et al., 1999; Schroeder and Leleshus, 2002; Bardashev et al., 2005), Laurussia (Arctic Canada, western Canada, eastern America, England to Poland, Urals, Russian Platform) (Lecompte, 1951, 1952; Galloway and St. Jean, 1957; Galloway, 1960; Galloway and Ehlers, 1960; Yavorsky, 1963; Birkhead, 1967; Stearn and Mehrotra, 1970; Kazmierczak, 1971; Bogoyavlenskaya, 1977a, b; Mistiaen, 1980; Stearn, 1983, 1990, 1996, 1998; Qi and Stearn, 1993; Kerbedunkel, 1995; Prosh and Stearn, 1996; Stock and Holmes, 1986; Stock, 1988, 1991, 1997; Salerno, 2008), northern Gondwana (Spain, Bohemia, Moravian Karst, Carnic Alps, North Africa, Afghanistan, Tibet) (Zukalova, 1971; Mistiaen, 1985; Méndez-Bedia et al., 1994; May, 1999, 2005, 2006, 2007; Fernández-Martínez et al., 2010; May and Rodriguez, 2012; Liang et al., 2023), South China (Dong, 2001; Huang et al., 2022), and Australia (Webby and Zhen, 1993, 1997, 2008; Webby et al., 1993; Cook, 1999). Figures 22 and 23 show the results of network analysis and are discussed later.

### Repository and institutional abbreviation

All specimens described in this study are housed in the Natural History Museum, London (UK), henceforth referred to as NHMUK. When there are several specimens cited consecutively, only the first one contains the prefix NHMUK, the following ones have it omitted.

## 3. Constructing elements of stromatoporoids

Devonian stromatoporoids consist of seven traditional stromatoporoid orders, including Labechiida, Clathrodictyida, Actinostromatida, Stromatoporellida, Stromatoporida, Syringostromatida, Amphiporida; an eighth group is considered as uncertain. Eight representative three-dimensional diagrams of type genera are illustrated for a better understanding of the stromatoporoid growth pattern (Fig. 2), outlined in the following paragraphs.

*Labechia* is the type genus of Labechiida. The skeleton of this genus is composed of long pillars and cyst plates (Fig. 2.1). In longitudinal section, pillars are thick, vertically continuous but oblique, therefore with different lengths. Cyst plates are thin, imbricating, convex-up, and distributed in the inter-pillar spaces. In tangential section, pillars are round, isolated dots, commonly interconnected with curved cyst plates.

*Anostylostroma* can be regarded as a type genus of Clathrodictyida. Laterally long laminae and short pillars (observed in longitudinal sections) are the internal features of this genus (Fig. 2.2). In longitudinal section, pillars are upward branching, commonly confined to the inter-laminar spaces. Laminae are continuous and are the dominant feature of the skeleton. In tangential section, pillars are isolated dots and laminae are curved.

*Actinostroma* is the representative genus of Actinostromatida. It is dominated by pillars, which regularly show radiating colliculi forming hexactinellid networks (Fig. 2.3). In longitudinal section, pillars are thick and commonly long, but locally cut as short ones due to the oblique outward distribution. Colliculi are discontinuous and appear as short rods or isolated dots due to their laterally dispersed radiation, which is different from laminae. In tangential section, pillars are circular and isolated between the interlaminar spaces.

*Trupetostroma* is a type genus of Stromatoporellida. The skeleton of this genus consists of thick pillars and long laminae (Fig. 2.4). In longitudinal section, pillars are short, confined to the inter-laminar spaces but systematically superposed. Laminae are laterally extensive, tripartite, commonly with central light zones and surrounding dark zones. In tangential section, pillars are isolated dots and laminae are curved. Some other genera of this order are typically ordinicellular (Stearn, 2015b) or show central dark zone and surrounding light zones possibly due to diagenesis.

*Stromatopora* is the conventional genus of Stromatoporida. It mainly comprises cassiculate structure (Fig. 2.5). In longitudinal section, cassiculate structure shows obliquely arranged skeletal elements, that in some cases may give an X-shaped appearance of layering within longitudinal sections. In tangential section, skeletons appear as labyrinthine networks, with irregular allotubes. Microstructure is cellular. Skeletons of Stromatoporida are characterized by amalgamated networks and cellular microstructure.

*Parallelopora* is a representative genus of Syringostromatida. The skeleton of this genus is featured by long pachysteles and suppressed pachystromes (Fig. 2.6). In longitudinal section, pachysteles are continuous and consist of micropillars and microlaminae. Pachystromes are rare or absent. In tangential section, pachysteles are interconnected to form closed networks, enclosing autotubes.

*Parallelostroma* is another typical genus of Syringostromatida. It is composed of short pachysteles and thick pachystromes (Fig. 2.7). In longitudinal section, pachysteles are confined to the inter-pachystrome spaces. Pachystromes are laterally continuous. Both skeletons consist of multiple microlaminae and micropillars. In tangential section, pachysteles are locally isolated. Closed networks also existed to enclose autotubes.

*Amphipora* is the type genus of Amphiporida. The growth form of this genus is dendroid. The skeleton is composed of the axial canal and peripheral portions (Fig. 2.8). In longitudinal section, pillars radiate upward and outward from the axis, connected with short elements to form irregular networks. Peripheral sheaths are sparsely developed. In tangential section, irregular networks enclose the axial canal.

Stearn (2015c) simplified the skeletons of the former six orders into four types: 1) domes and posts represented by cyst plates and pillars (i.e., Labechiida); 2) posts and beams formed by pillars and colliculi (i.e., Actinostromatida); 3) floors and posts constructed by laminae and pillars (i.e., Clathrodictyida and Stromatoporellida); 4) floors and walls formed by pachysteles and pachystromes (i.e., Stromatoporida and Syringostromatida). The four types indicate the general growth pattern of most stromatoporoids, which is significant and valuable in stromatoporoid identification.

## 4. Systematic paleontology

Phylum Porifera Grant, 1836

Class Stromatoporoidea Nicholson and Murie, 1878

Order Labechiida Kühn, 1927

Family Labechiidae Nicholson, 1879

Genus *Labechia* Edwards and Haime, 1851

*Type species*.—*Labechia conferta* (Lonsdale, 1839).

*Labechia stylophora* Nicholson, 1891 Figure 3.1, 3.2

1891 *Labechia stylophora* Nicholson, p. 161, pl. 20, figs. 7, 8.

*Holotype.—*Specimen (NHMUK PI P5987) from the Middle Devonian of Shaldon, in South Devon, UK (Nicholson, 1891, pl. 20, figs. 7, 8).

### Description

*Labechia stylophora* in this study is fragmented, but with an overall laminar form. Its skeletal structure consists of pillars with cyst plates between the pillars. In longitudinal section, pillars are thick, radiate slightly from the bottom and emerge to the surface (Fig. 3.1), spacing 2 to 4 per 2 mm, ranging from 0.1 to 0.3 mm in width. Cyst plates are abundant, imbricating, with widths from 0.5 to 0.7 mm. The galleries are generally crescent-shaped.

In tangential section, pillars are round, isolated (Fig. 3.2), but locally rodlike in oblique section. Cyst plates irregularly or circularly connect pillars. Mamelons are conspicuous, 8 to 10 mm in diameter, manifested by the circularly arranged pillars and cyst plates, with adjacent two mamelons at a distance of 15 mm. Astrorhizal canals are not clear. The microstructure is compact.

### Material

One specimen (NHMUK PI P5987) from the Middle Devonian of Shaldon, in South Devon (UK).

### Remarks

Compared with diverse Ordovician, Silurian and Late Devonian *Labechia*, the Middle Devonian records are rather rare. Nicholson (1891) established this species according to the unique growth form of cylinder-shaped mamelons from the other known species.

Genus *Labechiella* Yabe and Sugiyama, 1930

*Type species*.—*Labechia serotina* (Nicholson, 1886a).

*Labechiella serotina* (Nicholson, 1886a) Figure 3.3, 3.4

1886a *Labechia serotina* Nicholson, p. 15, pl. 2, figs. 3, 4.

**Figure 4.**
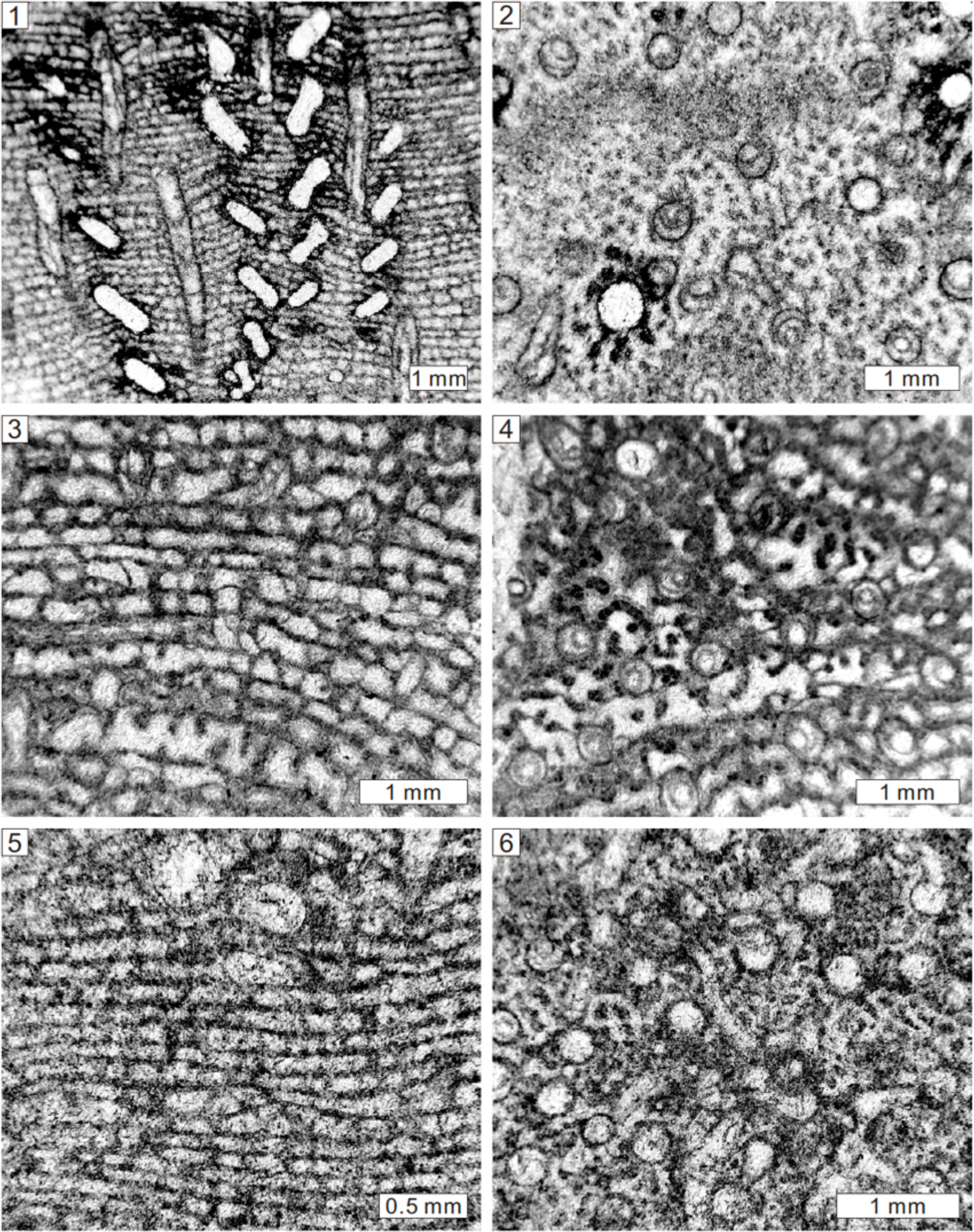
Longitudinal and tangential sections of *Clathrodictyon heathsense* Webby et al., 1993 from the Lower Devonian Cave limestone of New South Wales, Australia, NHMUK PI H3892 (**1, 2**), *Petridiostroma clarum* (Počta,1894) from the Lower Devonian of Commonwealth Quarries, Buchan, Victoria, Australia, NHMUK PI H3912 (**3, 4**), *Petridiostroma delicatulum* (Ripper, 1937a) from the Lower Devonian of Rocky Camp Quarry, Buchan, Victoria, Australia, NHMUK PI H3914 (**5, 6**).

2015 *Labechiella serotina*; Webby, p. 722.

### Holotype

Specimen (NHMUK PI P5988) from the Middle Devonian of Teignmouth, South Devon, UK (Nicholson, 1886a, pl. 2, figs. 3, 4).

### Description

Growth forms are unclear due to the limitation of the thin sections. Latilaminae exist and presumably represent either growth interruptions or growth phases in the specimens; each latilamina is 10 to 20 mm thick. The skeleton is composed of thick pillars and thin laterally distributed cyst plates. In longitudinal section, pillars are thick, continuous, post-like, closely or loosely distributed (Fig. 3.3), spacing 4 to 5 per 2 mm, ranging from 0.15 to 0.3 mm in width. In some areas, the axial zone of pillars shows possibly overlying cone-in-cone banding. Cyst plates are thin, flat, oblique or concave down, generally complete, with rare cases overlapping the adjacent one, spacing 6 to 8 per 2 mm, 0.02 mm in thickness. Galleries are rectangular or rarely irregular. In tangential section, pillars are circular, but closely or incompletely connected, approximately forming a vermiform skeleton, with rare, isolated ones (Fig. 3.4). The axial zone of certain pillars illustrates lighter or darker areas. Cyst plates are arched, can rarely be observed. Microstructure is compact.

### Material

Two specimens (NHMUK PI P5988, OR971), from the Middle Devonian of Teignmouth, in South Devon (UK).

### Remarks

The species was originally assigned as *Labechia* by Nicholson (1886a), but *Labechia* is markedly different from *Labechiella* in terms of imbricating cyst plates (Yabe and Sugiyama, 1930). Thus, this species was revised from *Labechia* into *Labechiella*. Detailed discussion between *Labechiella* and its related genera is provided by Webby (2015, pp. 722–723).

Order Clathrodictyida Bogoyavlenskaya, 1969

Family Clathrodictyidae Kühn, 1939

Genus *Clathrodictyon* Nicholson and Murie, 1878

*Type species*.—*Clathrodictyon vesiculosum* Nicholson and Murie, 1878.

*Clathrodictyon confertum* Nicholson, 1889 Figure 3.5, 3.6

1889 *Clathrodictyon confertum* Nicholson, p. 154, pl. 18, figs. 13, 14.

1957 *Clathrodictyon confertum*; Galloway and St. Jean, p. 94, pl. 1, fig. 2a, b.

1985 *Clathrodictyon confertum*; Bjerstedt and Feldmann, p. 1055, figs. 11.7, 11.8.

### Holotype

Specimen (NHMUK PI P5981) from the Middle Devonian of Pit Park Quarry, Dartington, South Devon, UK (Nicholson, 1889, pl. 18, figs. 13, 14).

### Description

External forms are unconfirmed because of the fragmented skeleton. Skeleton is formed by densely distributed laminae and short pillars. In longitudinal section, pillars are short, confined to the interlaminar spaces (Fig. 3.5), irregularly distributed, 0.02 mm in width. Laminae are thin, undulating, crumpled, locally continuous, but generally intermittent, spacing 14 to 16 per 2 mm, 0.02 mm in thickness. Galleries are irregular, circular or sub-rectangular. In tangential section, laminae are not apparent. Pillars have locally been observed as isolated ones, but are commonly connected with other elements (Fig. 3.6). Astrorhizal canals are unclear. The microstructure is compact.

### Material

Three specimens from the Middle Devonian of Pit Park Quarry, Dartington (NHMUK PI P5981) and Hope’s Nose Quarry, Torquay (NHMUK PI H3891, P6110), in South Devon (UK).

### Remarks

This species is featured by densely packed thin wrinkled laminae and short pillars. It closely resembles *Clathrodictyon vesiculosum* in terms of wrinkled laminae and confined pillars, but this species presents a smaller and closely distributed skeleton (Nicholson, 1889; Galloway and St. Jean, 1957).

*Clathrodictyon heathsense* Webby et al., 1993 Figure 4.1, 4.2

1937a *Clathrodictyon regulare*; Ripper, p. 16, pl. 3, figs. 1, 2.

1937a *Clathrodictyon convictum*; Ripper, p. 19, pl. 3, figs. 4–8.

1993 *Clathrodictyon*? *heathsense*; Webby et al., p. 134, figs. 11C–F, 12A.

### Holotype

Longitudinal and tangential sections (NMV P I41798 and NMV P I41799) from the Lower Devonian of Heath’s Quarry, Buchan Caves Limestone, Buchan, Victoria, Australia (Webby et al., 1993, fig. 11C, F).

### Description

Specimens were preserved as fragments, growth forms are therefore unclear. The skeleton comprises short pillars and laterally extensive laminae. In longitudinal section, pillars are short, post-like or spool-shaped, generally confined to one interlaminar space (Fig. 4.1), spacing about 7 to 9 per 2 mm and 0.05 to 0.2 mm in width. Laminae are thin, inflected upward and downward, horizontally continuous, spacing about 9 to 11 per 2 mm and 0.05 to 0.15 mm in thickness. Galleries are rectangular or locally circular.

Syringoporid symbionts, 0.3 to 0.5 mm in diameter, spacing 2 to 3 per 2 mm, are associated, as well as spiral upward tubeworms (Fig. 4.1). In tangential section, pillars are circular or irregular, commonly isolated (Fig. 4.2), in diameter of 0.1 to 0.2 mm. Laminae are irregularly undulating. Syringoporids and tubeworms are circular. Microstructure is compact.

### Material

Six specimens from the Lower Devonian of Heath’s Quarry (NHMUK PI H3917, H3918, H3922, H3924), in Buchan, Cave limestone (NHMUK PI H3892) of New South Wales, and Mitchel’s Quarry, Cave Hill, Lilydale (NHMUK PI H3900), all in Victoria (Australia).

### Remarks

This species was identified as *Clathrodictyon regulare* and *Clathrodictyon convictum* Ripper (1937a). However, the type specimens of these two species were revised to *Petridiostroma* according to regular skeletons formed by planar laminae and short pillars (Stearn, 1992). The Australian materials, which closely resemble *Clathrodictyon*, show slightly bent laminae. Webby et al. (1993) thus reassigned these specimens as

*Clathrodictyon heathsense*. This study follows the identification of Webby et al. (1993).

Family Gerronostromatidae Bogoyavlenskaya, 1969 Genus *Petridiostroma* Stearn, 1992

*Type species*.—*Simplexodictyon simplex* Nestor, 1966.

*Petridiostroma clarum* (Počta, 1894) Figure 4.3, 4.4

1894 *Clathrodictyon clarum* Počta, p. 152, pl. 18, figs. 7, 8.

1937a *Clathrodictyon clarum*; Ripper, p. 21, pl. 4, figs. 3, 4.

1993 *Petridiostroma clarum*; Webby et al., p. 132, figs. 10A–F, 31A.

### Holotype

Specimen from the Middle Devonian of the Prague Basin, Czech Republic (Počta, 1894, pl. 18, figs. 7, 8).

### Description

External form is unclear based on fragmented skeletons. The skeleton consists of laterally extensive laminae and short pillars. In longitudinal section, pillars are short, post-like, generally confined to one interlaminar space, rarely superposed (Fig. 4.3), spacing about 6 to 8 per 2 mm and 0.02 to 0.1 mm in width. Laminae are thin, planar, laterally continuous, spacing about 8 to 10 per 2 mm and 0.02 to 0.1 mm in thickness. Galleries are laterally elongate or circular. In tangential section, pillars and laminae are isolated dots and undulating (Fig. 4.4), respectively. Syringoporid tabulate corals are in width of 0.3 to 0.4 mm, spacing 3 to 4 per 2 mm.

### Material

Two specimens (NHMUK PI H3912, H3926) from Lower Devonian of Commonwealth Quarries, Buchan, Victoria (Australia).

### Remarks

This species was originally grouped with *Clathrodictyon*, but the common planar laminae show a close relationship with *Petridiostroma*.

*Petridiostroma delicatulum* (Ripper, 1937a) Figure 4.5, 4.6

1937a *Clathrodictyon convictum* var. *delicatula* Ripper, p. 20, pl. 4, figs. 1, 2. 1993 *Petridiostroma delicatulum*; Webby et al., p. 134, fig. 9E, F.

### Holotype

Specimen (NHMUK PI H3914) from the Lower Devonian of Rocky Camp (Commonwealth) Quarry, Buchan, Victoria, Australia (Ripper, 1937a, pl. 4, figs. 1, 2).

### Description

Growth forms appear to be laminar, but were cut as fragments. The skeleton is composed of short pillars and horizontally continuous laminae. In longitudinal section, pillars are short, post-like or spool-shaped, commonly confined to one interlaminar space, locally superposed across laminae, spacing about 12 to 14 per 2 mm and 0.05 to 0.2 mm in width (Fig. 4.5).

Laminae are thin, slightly undulating, laterally extensive, spacing about 15 to 17 per 2 mm and 0.05 to 0.1 mm in thickness. Galleries are rectangular or circular. Syringoporid tabulate corals are common, with a width of 0.3 to 0.4 mm, spacing 2 to 4 per 2 mm. In tangential section, pillars are circular, isolated, in diameter of 0.05 to 0.15 mm (Fig. 4.6). Laminae are irregularly undulating. Syringoporids are circular in tangential sections. Microstructure is compact.

### Material

One specimen (NHMUK PI H3914) from the Lower Devonian of Rocky Camp (Commonwealth) Quarry, Buchan, Victoria (Australia).

### Remarks

This species is characterized mainly by planar laminae and short pillars, which are close to *Petridiostroma* rather than *Clathrodictyon*. It is different from *P*. *clarum* concerning more densely distributed laminae and pillars.

*Petridiostroma* sp. Figure 5.1, 5.2

**Figure 5.**
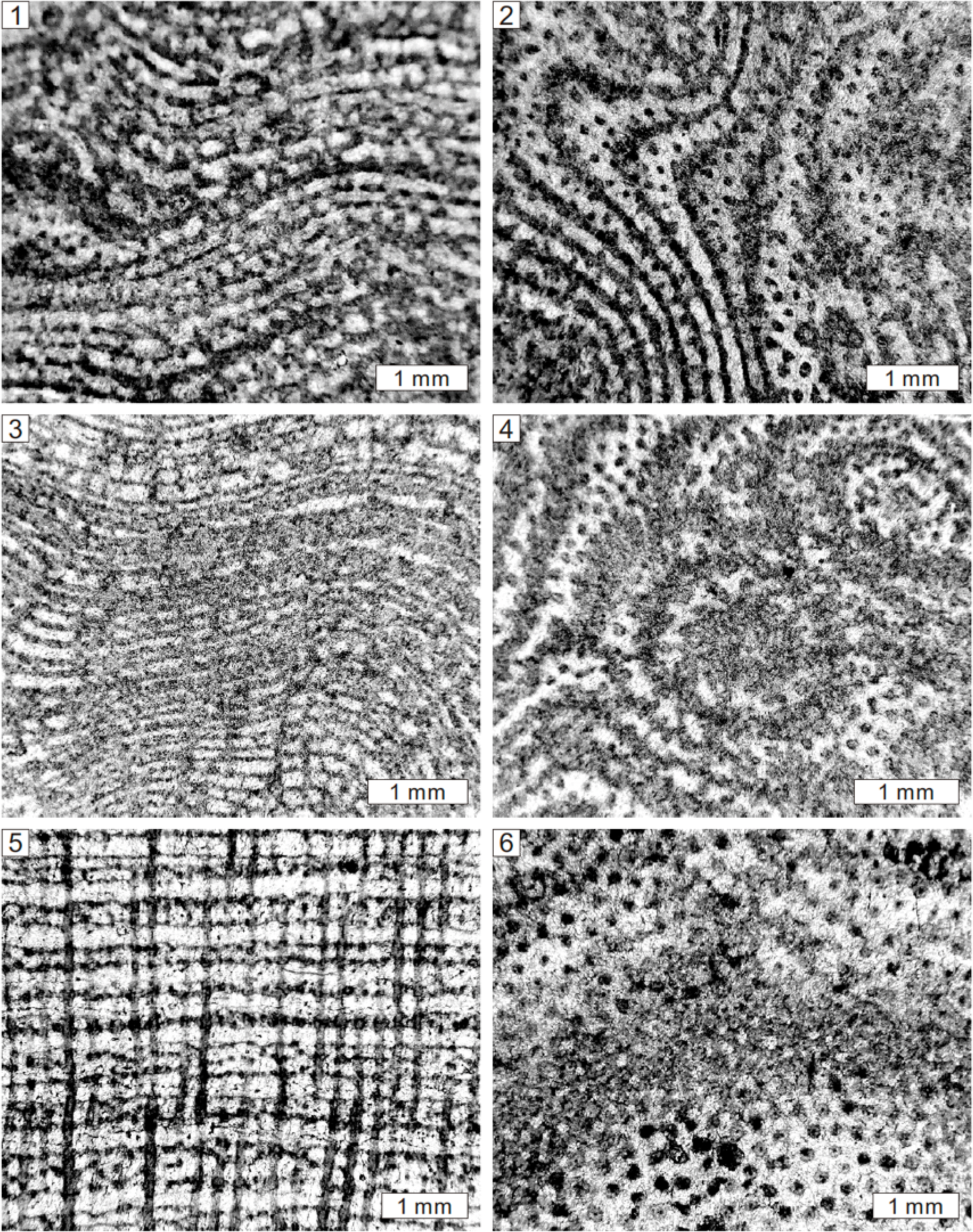
Longitudinal and tangential views of *Petridiostroma* sp. from the Lower Devonian of Griffith’s Quarry, Loyola, Victoria, Australia, NHMUK PI H3930 (**1, 2**), *Gerronostromaria buchanense* (Flügel, 1959) from the Lower Devonian of Commonwealth Quarries, Buchan, Victoria, Australia, NHMUK PI H3913 (**3, 4**), *Actinostroma clathratum* Nicholson, 1886a from the Middle Devonian of Gerolstein, Germany, NHMUK PI P5774 (**5, 6**).

1933 *Clathrodictyon regulare cylindriferum*; Ripper, p. 157, pl. 3, fig. 6A, B.

**Figure 6.**
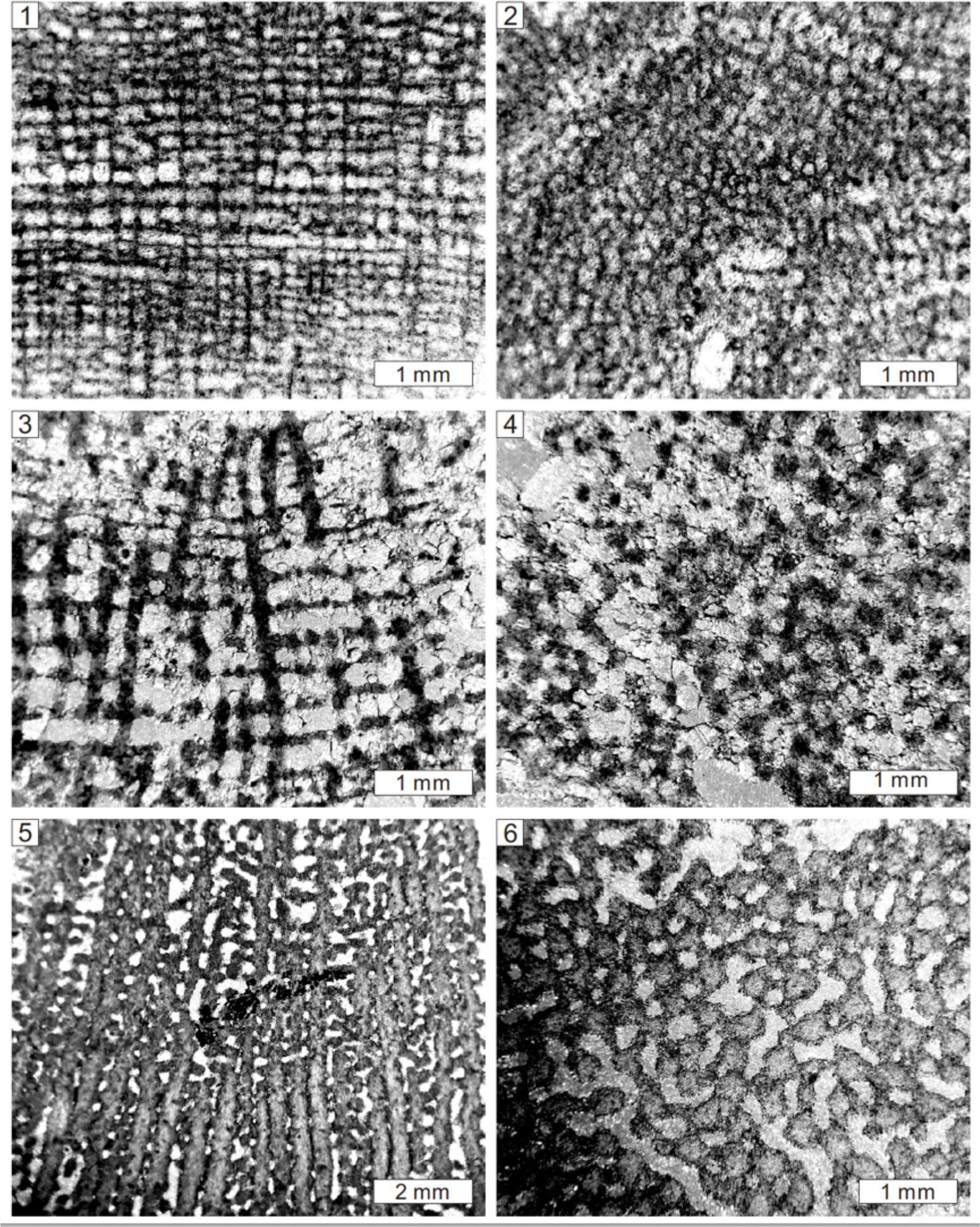
Longitudinal and tangential sections of *Actinostroma compactum* Ripper, 1933 from the Lower Devonian of Rocky Camp Quarry, Buchan, Victoria, Australia, NHMUK PI H3910 (**1, 2**), *Actinostroma expansum* (Hall and Whitfield, 1873) from the Devonian of Rochford, Iowa (USA), NHMUK PI P5803 (**3, 4**), *Actinostroma fenestratum* Nicholson, 1889 from the Devonian of Lake Winnipeg, Manitoba, Canada, NHMUK PI P5601 (**5, 6**).

1937b *Clathrodictyon regulare*; Ripper, p. 2, pl. 1, figs. 1, 2.

1937b *Clathrodictyon* aff. *chapmani*; Ripper, p. 3, pl. 1, figs. 3, 4.

1993 *Petridiostroma* sp.; Webby et al., p. 134, fig. 11A, B.

### Description

Growth forms are possibly laminar, but poorly preserved. The skeleton comprises short pillars and laterally extensive laminae. In longitudinal section, pillars are short, spool-shaped, generally confined to one interlaminar space, locally superposed, spacing about 7 to 10 per 2 mm and 0.05 to 0.2 mm in width (Fig. 5.1). Laminae are thin, gently undulating, horizontally continuous, spacing about 9 to 11 per 2 mm and 0.05 to 0.15 mm in thickness. Galleries are round or rarely rectangular. In tangential section, pillars are circular or irregular, commonly isolated, in diameter of 0.1 to 0.2 mm (Fig. 5.2). Laminae are irregularly undulating. Microstructure is compact. *Material*.—One specimen from the Lower Devonian of Griffith’s Quarry (NHMUK PI H3930), Loyola, Victoria (Australia).

### Remarks

It was assigned by Ripper (1937b) to *Clathrodictyon regulare* and *Clathrodictyon* aff. *chapmani*, but *C*. *regulare* show closely spaced laminae of 18 per 2 mm (Nestor, 1962, 1964) and the type specimen of the latter species (Ripper, 1933) was revised into *Atelodictyon* (Webby et al., 1993). In addition, the planar lamiae resemble *Petridiostroma*. The current species is left in open nomenclature due to the poorly preserved skeleton. It features round galleries and finer texture, which is different from *P*. *delicatulum*.

Genus *Gerronostromaria* Nestor, 2011

*Type species*.—*Gerronostroma elegans* Yavorsky, 1931

*Gerronostromaria buchanense* (Flügel, 1959) Figure 5.3, 5.4

1937b *Actinostroma contortum*; Ripper, p. 14, pl. 2, figs. 3–6.

1959 *Actinostroma buchanense*; Flügel, p. 183, pl. 7, fig. 4.

1993 *Gerronostromaria buchanense*; Webby et al., p. 130, fig. 9A–D.

### Holotype

Specimen (NMV P141758) from the Lower Devonian Buchan Caves Limestone at Heath’s Quarry, Buchan, Victoria, Australia (Webby et al., 1993, fig. 9A).

### Description

Growth forms were described as domical or bulbous by Ripper (1937b). In longitudinal section, pillars are long, passing through several laminae, spacing about 10 to 12 per 2 mm and 0.05 to 0.2 mm in width (Fig. 5.3). Laminae are thin, apparently undulating, laterally continuous, spacing about 14 to 17 per 2 mm and 0.05 to 0.15 mm in thickness. Galleries are rectangular. In tangential section, pillars are circular and isolated, in diameter of 0.1 to 0.2 mm. Very tiny rod-like structures radiate from pillars, less than 0.01 mm in width. Laminae are concentric circular shapes (Fig. 5.4). Mamelons are distinct, in diameter of 4 to 5 mm. Microstructure is compact. *Material*.—Three specimens (NHMUK PI H3909, H3911, H3913) from the Lower Devonian of Commonwealth Quarry, Buchan, Victoria (Australia). *Remarks*.—This species was assigned into *Actinostroma* according to tiny rod-like structures by Ripper (1937b), but colliculi and hexactinellid networks in typical *Actinostroma* (Fig. 1.3) have not been noticed. Regular networks formed by extensive pillars and laminae justify its assignment to *Gerronostromaria*.

Order Actinostromatida Bogoyavlenskaya, 1969

Family Actinostromatidae Nicholson, 1886

Genus *Actinostroma* Nicholson, 1886a

*Type species*.—*Actinostroma clathratum* Nicholson, 1886a.

*Actinostroma clathratum* Nicholson, 1886a Figure 5.5, 5.6

1886a *Actinostroma clathratum* Nicholson, p. 226, pl. 6, figs. 1–3.

1930 *Actinostroma clathratum*; Yavorsky, p. 475, pl. 1, figs. 1–10.

1940 *Actinostroma clathratum*; Chi, p. 09, pl. 2, fig. 2.

1951 *Actinostroma clathratum*; Lecompte, p. 99, pl. 1, figs. 1–12.

1957 *Actinostroma clathratum*; Galloway and St. Jean, p. 149, pl. 10, fig. 4a, b.

1971 *Actinostroma clathratum*; Zukalová, p. 31, pl. 3, figs. 3–5.

1979 *Actinostroma clathratum*; Yang and Dong, p. 31, pl. 9, figs. 1, 2.

1982 *Actinostroma clathratum*; Stock, p. 669, pl. 3, figs. 5, 6.

1989 *Actinostroma clathratum*; Dong et al., p. 264, pl. 5, fig. 2a, b.

1993 *Actinostroma clathratum*; Webby and Zhen, p. 329, fig. 3A–D.

1995 *Actinostroma clathratum*; Krebedünkel, p. 23, pl. 1, figs. 1, 2.

1997 *Actinostroma clathratum*; Webby and Zhen, p. 13, fig. 6A, 6B, 6D.

1999 *Actinostroma clathratum*; Stearn et al., p. 329, fig. 5E–F.

2008 *Actinostroma clathratum*; Salerno, p. 52, pl. 6, fig. 3.

### Holotype

Lectotype (NHMUK PI P5774, Stock, 2015) from the Middle Devonian of Gerolstein, Germany (Nicholson, 1889, pl. 1, figs. 11, 12).

### Description

Growth forms are unclear based on thin sections, but Nicholson (1886a) noted that they are commonly bulbous or domical. The skeleton comprises vertically dominant pillars and laterally subordinate colliculi. In longitudinal section, pillars are thick, long, continuous (Fig. 5.5), possibly throughout the whole skeleton or at least each latilaminae, spacing about 8 to 11 per 2 mm and 0.1 to 0.2 mm in width. In some oblique sections, pillars appear to be short, passing through 1 to 3 laminae. Colliculi are thin, gently undulating, locally cut as isolated circular dots, spacing about 7 to 9 per 2 mm and 0.05 to 0.15 mm in thickness. Galleries are rectangular or round. In tangential section, pillars are circular and isolated between the interlaminar spaces, in diameter of 0.1 to 0.2 mm. Colliculi are interconnected in the previously called laminae, forming a hexactinellid network (Fig. 5.6). *Microstructure is compact*.

### Material

26 specimens from the Middle Devonian of Dartington (NHMUK PI P5708, H3362), Teignmouth (NHMUK PI P5716, P5717, P5719, P5720, P5721), Kingsteignton (NHMUK PI H4335, H4336), Torquay (NHMUK PI H5286) and six unknown locations (NHMUK PI P5718, P2202, P2227, R1217, 25, 15243), in South Devon (UK); Gerolstein (NHMUK PI P5774, P5779, P5787), Sötenich (NHMUK PI P5777), Hebborn (NHMUK PI P5778, P5783, P5786), and Büchel (NHMUK PI P6090), in Germany; and Rough Range (NHMUK PI P4463, P4967) near Mt. Krauss, Kimberley (western Australia). *Remarks*.—According to Nicholson (1886a), this species exhibits variability in external form, but the internal structure is constant. It is characterized by regularly developed long pillars and interrupted colliculi. It resembles *A*. *expansum*, but differs from the latter in thinner skeletons.

*Actinostroma compactum* Ripper, 1933 Figure 6.1, 6.2

1933 *Actinostroma compactum* Ripper, p. 153, fig. 5A, B.

1937a *Actinostroma compactum*; Ripper, p. 15, pl. 2, figs. 7–8.

**Figure 7.**
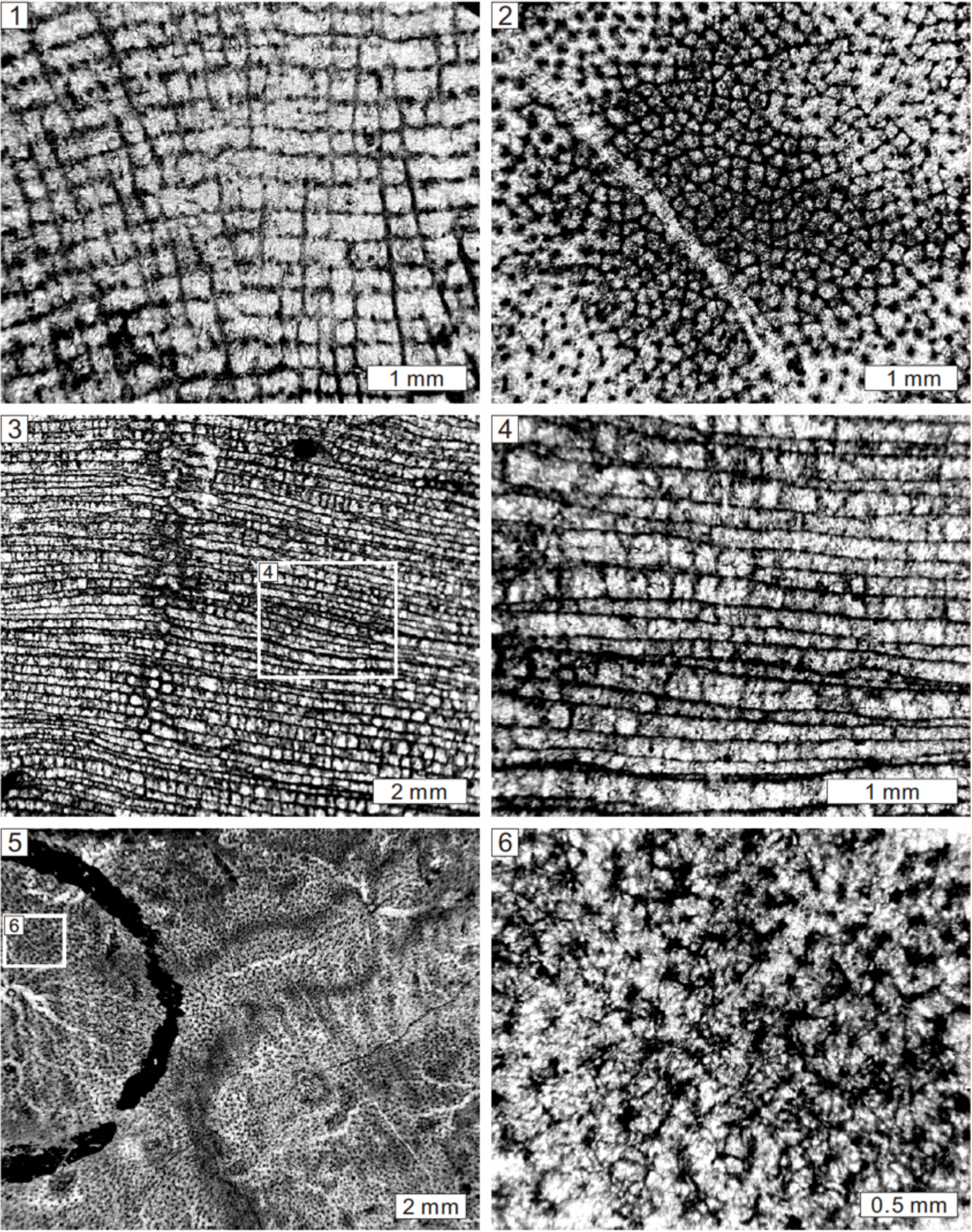
Longitudinal and tangential views of *Actinostroma hebbornense* Nicholson, 1889 from the Middle Devonian of Hebborn, Germany, NHMUK PI P5581 (1, 2), *Actinostroma stellulatum* Nicholson, 1886a from the Middle Devonian of Gerolstein, Germany, NHMUK PI P5570 (3–6).

1993 *Actinostroma compactum*; Webby et al., p. 123, figs. 5A–F, 7A–C.

1997 *Actinostroma compactum*; Webby and Zhen, p. 17, fig. 7A, B.

1999 *Actinostroma compactum*; May, p. 99, pl. 1, fig. 1.

2007 *Actinostroma compactum*; May, p. 143.

### Holotype

Specimen (NMV P141959-60, Webby et al., 1993) from the Lilydale Limestone at Mitchell’s (Cave Hill) Quarry, Lilydale, Victoria, Australia (Ripper, 1933, fig. 5A, B).

### Description

External forms are uncertain due to poor preservation. The skeleton comprises vertically long pillars and laterally colliculi. In longitudinal section, pillars are long, continuous, passing through dozens of colliculi (Fig. 6.1), spacing about 10 to 12 per 2 mm and 0.1 to 0.15 mm in width. Colliculi are thin, locally oblique, laterally discontinuous, showing isolated circular dots, spacing about 12 to 14 per 2 mm and 0.05 to 0.1 mm in thickness. Galleries are rectangular, irregular or round. In tangential section, pillars are circular and isolated between the interlaminar spaces, in diameter of 0.1 to 0.15 mm. Hexactinellid networks occur from 4 to 5 colliculi interconnecting each pillar (Fig. 6.2). Tubeworms are locally observed, in a diameter of 1 mm.

Microstructure is compact.

### Material

Two specimens (NHMUK PI H3910, H3923) from the Lower Devonian of Rocky Camp (Commonwealth) Quarry, Buchan, Victoria (Australia).

### Remarks

Compared with *A*. *clathratum*, this species shows more densely packed and thinner pillars and colliculi.

*Actinostroma expansum* (Hall and Whitfield, 1873) Figure 6.3, 6.4

1873 *Stromatopora expansa* Hall and Whitfield, p. 226, pl. 9, fig. 1.

1891 *Actinostroma expansum*; Nicholson, p. 316, pl. 10, fig. 1, 2.

1971 *Actinostroma expansum*; Kaźmierczak, p. 136, pl. 40, fig. 2a, b; pl. 41, fig. 5.

1984 *Actinostroma expansum*; Stock, p. 774, fig. 2A–E.

1996 *Actinostroma expansum*; Stearn, p. 200, figs. 2.4, 2.6, 3.1, 3.2.

### Holotype

Specimen (NYSM 331) from the Upper Devonian Nora Member of the Shell Rock Formation, Rockford, Floyd County, Iowa, USA. (Stock, 1984, fig. 2A–D).

### Description

External forms are unclear due to poor preservation. Skeleton is composed of vertically thick pillars and laterally colliculi. In longitudinal section, pillars are long, continuous, passing through several colliculi (up to 19), spacing about 6 to 8 per 2 mm and 0.1 to 0.25 mm in width (Fig. 6.3). Colliculi are thin, undulating, laterally discontinuous, locally showing isolated circular dots, spacing about 7 to 10 per 2 mm and 0.05 to 0.1 mm in thickness.

Galleries are rectangular or round. In tangential section, pillars are circular and isolated, in diameter of 0.1 to 0.2 mm. Hexactinellid networks are formed by 4 to 5 colliculi interconnecting each pillar (Fig. 6.4). Microstructure is compact.

### Material

Three specimens from the Upper Devonian: two of Rochford (NHMUK PI P5803, P5807), Iowa (USA) and another one from Rough Range near Mt. Krauss (NHMUK PI P4463), Kimberley (western Australia).

### Remarks

This species differs from other species in widely spaced and thicker colliculi and pillars.

*Actinostroma fenestratum* Nicholson, 1889 Figure 6.5, 6.6

1889 *Actinostroma fenestratum* Nicholson, p. 146, pl. 17, figs. 8, 9.

**Figure 8.**
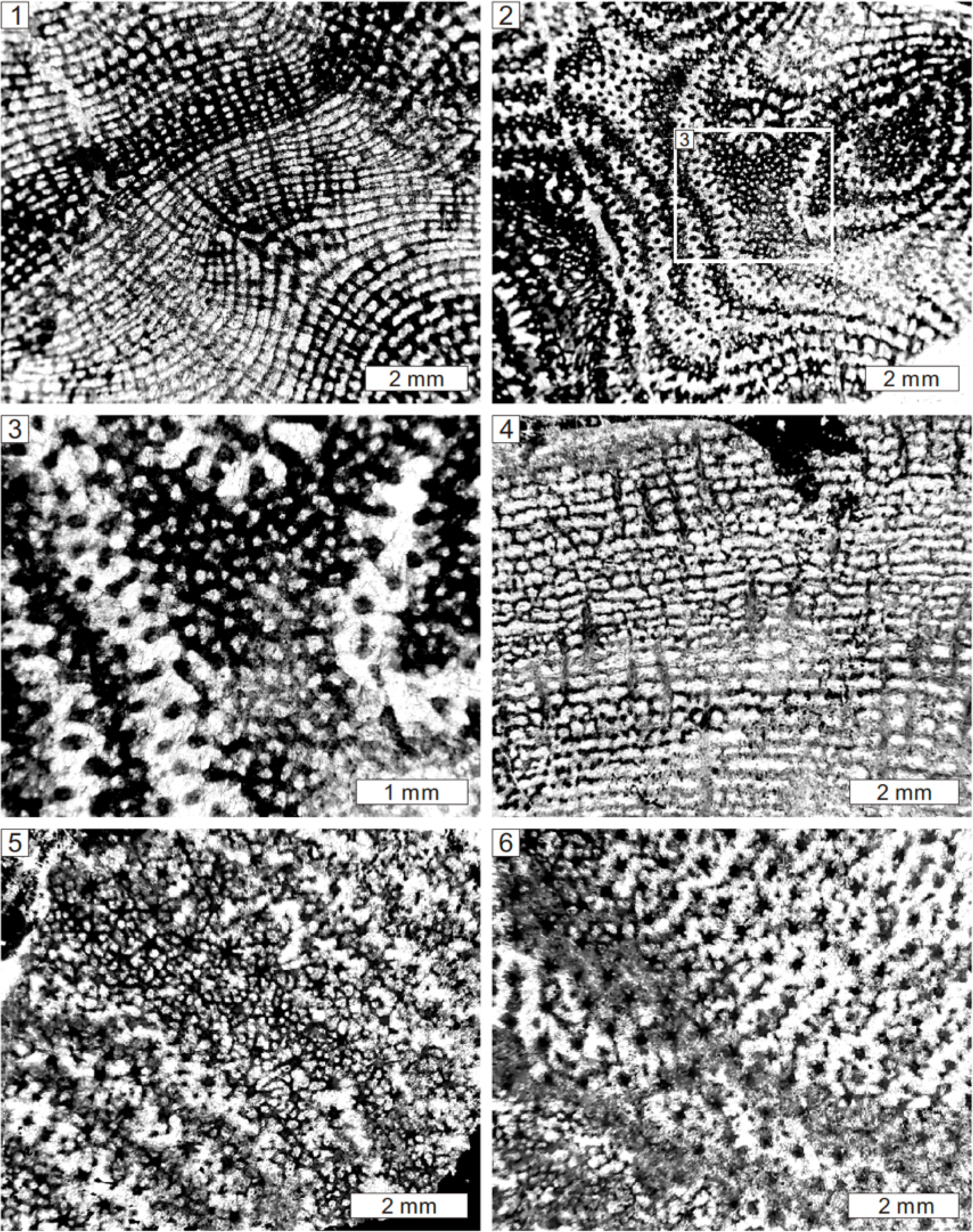
Longitudinal and tangential sections of *Actinostroma verrucosum* (Goldfuss, 1826) from the Middle Devonian of Büchel, Germany, NHMUK PI P5537 (1–3), *Bifariostroma bifarium* (Nicholson, 1889) from the Middle Devonian of Büchel, Germany, NHMUK PI P5639 (4–6).

**Figure 9.**
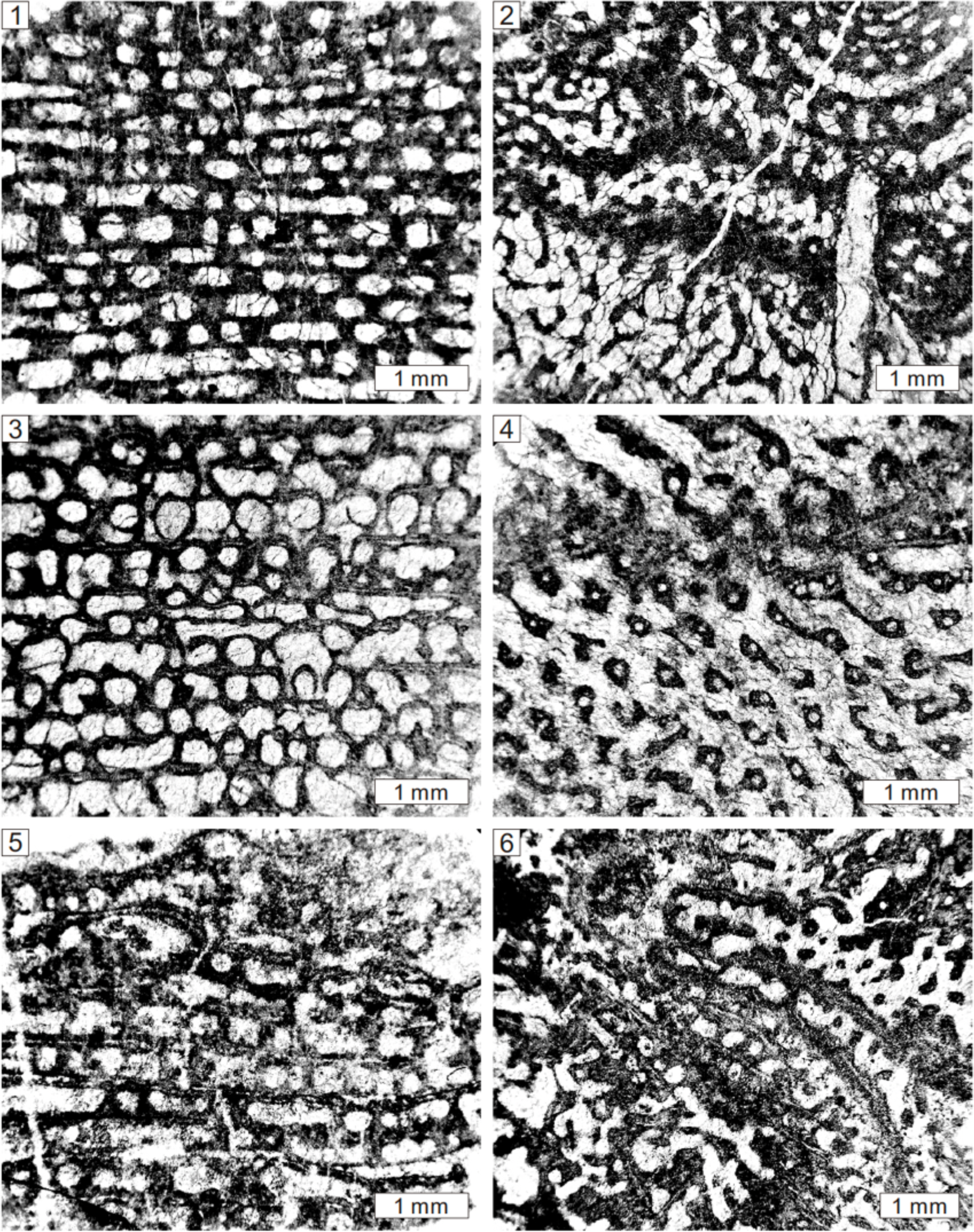
Longitudinal and tangential views of *Stromatoporella arachnoidea* Nicholson, 1886b from the Middle Devonian of Büchel, Germany, NHMUK PI P6063 (1, 2), *Stromatoporella solitaria* Nicholson, 1892 from the Middle Devonian of Gerolstein, Germany, NHMUK PI P6037 (3, 4), *Clathrocoilona curiosa* (Bargatzky, 1881) from the Middle Devonian of Büchel, Germany, NHMUK PI P6051 (5, 6).

1891 *Actinostroma fenestratum*; Nicholson, p. 322, pl. 10, figs. 3, 4.

### Holotype

Specimen (NHMUK PI P6116) from the Middle Devonian of Teignmouth, South Devon, UK (Nicholson, 1889, pl. 17, figs. 8, 9).

### Description

Growth forms were described as domical/bulbous by Nicholson (1889, 1891). The skeleton consists mainly of long pillars and subordinate colliculi. In longitudinal section, pillars are rodlike, thick, long, continuous, spacing about 5 to 6 per 2 mm and 0.3 to 0.6 mm in width (Fig. 6.5). Colliculi are generally cut as isolated circular dots, spacing about 4 to 5 per 2 mm and 0.1 to 0.3 mm in thickness. Galleries are irregular. In tangential section, pillars are round and isolated, in diameter of 0.2 to 0.5 mm. Colliculi are not well developed, interconnecting with pillars, forming irregular or vermicular networks (Fig. 6.6). Microstructure is compact.

### Material

Four specimens from the Devonian: one from Lake Winnipeg (NHMUK PI P5601), in Manitoba (Canada) and three more from Teignmouth (NHMUK PI P5884, P6111, P6116), in South Devon (UK).

### Remarks

This species is different from other species by thick, long pillars and poorly developed colliculi and networks. It also resembles *Neosyringostroma logansportense*, but the latter shows more well-developed lateral skeletons and cellular microstructure.

*Actinostroma hebbornense* Nicholson, 1886b Figure 7.1, 7.2

1886b *Actinostroma hebbornense* Nicholson, p. 228, pl. 7, figs. 7, 8.

1889 *Actinostroma hebbornense*; Nicholson, p. 137, pl. 16, figs. 9–16.

1951 *Actinostroma hebbornense*; Lecompte, p. 92, pl. 3, figs. 4–6.

1971 *Actinostroma hebbornense*; Zukalová, p.30, pl. 3, figs. 1, 2

1985 *Actinostroma hebbornense*; Neidhardt, p. 16, pl. 1, figs. 5, 6.

1995 *Actinostroma hebbornense*; Krebedünkel, p. 30, pl. 1, figs. 7, 8.

### Holotype

Specimen (NHMUK PI P5581) from the Middle Devonian of Hebborn, Germany (Nicholson, 1886b, pl. 7, figs. 7, 8).

### Description

External forms have been noted as domical or bulbous by Nicholson (1889). The skeleton is composed of vertically thick pillars and lateral colliculi. In longitudinal section, pillars are rodlike, long, continuous, but commonly appear to be short in oblique sections (Fig. 7.1), spacing about 9 to 11 per 2 mm and 0.1 to 0.2 mm in width. Colliculi are generally undulating, commonly cut as isolated circular dots, spacing about 10 to 14 per 2 mm and 0.05 to 0.1 mm in thickness. Galleries are rectangular or locally circular. In tangential section, pillars are round and isolated dots between interlaminar spaces, in diameter of 0.05 to 0.2 mm. Colliculi are rodlike, 0.05 to 0.1 mm in width, interconnecting with pillars, forming hexactinellid networks (Fig. 7.2). Syringoporids and tubeworms locally occur. Microstructure is compact. *Material*.—Five specimens from the Middle Devonian: two from Teignmouth (NHMUK PI P5579, P5950), one from Torquay (NHMUK PI P6106), in South Devon (UK) and another two more from Devonian of Hebborn (NHMUK PI P5573, P5581) in Germany.

### Remarks

According to Nicholson (1889), this species is featured by regular hexactinellid networks, slender and densely arranged pillars, and the uncommon occurrence of astrorhizae. It is different from *A*. *clathratum* in thinner and oblique pillars. It is distinguished from *A*. *compactum* by regular and planar colliculi.

*Actinostroma stellulatum* Nicholson, 1886b Figure 7.3–7.6

1886b *Actinostroma stellulatum* Nicholson, p. 231, pl. 6, figs. 8, 9.

1889 *Actinostroma stellulatum*; Nicholson, p. 140, pl. 14, figs. 1–8; pl. 15.

1951 *Actinostroma stellulatum*; Lecompte, p. 111, pl. 11, figs. 1–5.

1969 *Actinostroma stellulatum*; Sleumer, p. 34, pl. 21, fig. 1; pl. 22, fig. 2.

1971 *Actinostroma stellulatum*; Kaźmierczak, p. 133, pl. 37, fig. 3a, b.

1993 *Actinostroma stellulatum*; May, p. 29, pl. 2, fig. 1.

1995 *Actinostroma stellulatum*; Krebedünkel, p. 45, pl. 4, figs. 1, 2.

1999 *Actinostroma stellulatum*; Méndez-Bedia, p. 124, fig. 2A, B.

2008 *Actinostroma stellulatum*; Salerno, p. 60, pl. 9, fig. 1.

### Holotype

Specimen (NHMUK PI P5570) from the Middle Devonian of Gerolstein, Germany (Nicholson, 1886b, pl. 6, figs. 8, 9).

### Description

Growth forms are laminae based on limited specimens, but Nicholson (1889) also noted the occurrence of bulbous or columnar shapes. The skeleton consists of vertically short pillars and laterally long laminae. In longitudinal section, pillars are short (Fig. 7.3), commonly upward spreading (Fig. 7.4), confined to one interlaminar space, locally superposed, spacing about 9 to 14 per 2 mm and 0.1 to 0.15 mm in width. Colliculi are thin, laterally continuous, slightly undulating, spacing about 11 to 13 per 2 mm and 0.05 to 0.1 mm in thickness, with a central dark line. Galleries are arch-shaped, rectangular, or round. Mamelons are inconspicuous, 0.2 mm in height. In tangential section, pillars are round and isolated, in diameter of 0.1 to 0.15 mm. Colliculi are curved (Fig. 7.5). Radial processes are small, thin, rod-like, in diameter of 0.01 to 0.03 mm, radiating from pillars and forming regular hexactinellid networks between the inter-laminae spaces (Fig. 7.6). Astrorhizae are distinct, with central and branching canals, in diameter of 5 to 7 mm. Microstructure is compact.

### Material

16 specimens from the Middle Devonian of Dartington (NHMUK PI P5943, P5948), Teignmouth (NHMUK PI P5942, P5944, P5945, P5946, P5947, P5949), Bishopsteignton (NHMUK PI H4334, H4340) in South Devon (UK), and Gerolstein (NHMUK PI P5570, P5583, P5858, P5860, P5862, P6089), in Germany.

### Remarks

As noted by Stock in Stearn et al. (1999), this species is different from others in atypical features, such as short pillars and thin colliculi.

*Actinostroma verrucosum* (Goldfuss, 1826) Figure 8.1–8.3

1826 *Ceriopora verrucosa* Goldfuss, p.33, pl. 10, fig. 6a–c.

1881 *Stromatopora verrucosa*; Bargatzky, p. 55.

1889 *Actinostroma verrucosum*; Nicholson, p. 134, pl. 16, figs. 1–8.

1951 *Actinostroma verrucosum*; Lecompte, p. 107, pl.. 9, figs. 1–8; pl. 10, figs. 1, 2.

1959 *Actinostroma verrucosum*; Flügel, p. 190, pl. 6, fig. 4.

1971 *Actinostroma verrucosum*; Zukalová, p. 36, pl.. 5, figs. 5, 6.

1980 *Actinostroma verrucosum*; Mistiaen, p. 185, pl.. 2, figs. 7–9; pl. 3, figs. 1–3.

1993 *Actinostroma verrucosum*; May, p. 28, pl. 2, fig. 2.

1995 *Actinostroma verrucosum*; Krebedünkel, p. 36, pl. 5, figs. 5, 6.

1999 *Actinostroma verrucosum*; Méndez-Bedia, p. 124, fig. 2C–E.

2008 *Actinostroma verrucosum*; Salerno, p. 63, pl. 8, fig. 1–3, pl. 12.

### Holotype

Lectotype (Flügel, 1959) from the Middle Devonian of Bensberg, Germany (Goldfuss, 1826, pl. 10, fig. 6).

### Description

External forms are unclear due to poor preservation, but Nicholson (1889) mentioned the existence of bulbous or domical shapes. The skeleton is composed of vertically thick pillars and laterally colliculi. In longitudinal section, pillars are post-like or spool-shaped, confined to one interlaminar space but systematically superposed, and commonly appear to be short in oblique sections, spacing about 8 to 10 per 2 mm and 0.1 to 0.2 mm in width (Fig. 8.1). Colliculi are conspicuously undulating, locally showing isolated circular dots, spacing about 10 to 12 per 2 mm and 0.05 to 0.1 mm in thickness. Galleries are circular. Mamelons are conspicuous, 2 to 3 mm in height. In tangential section, pillars are round and isolated between the interlaminar spaces (Fig. 8.2). 5 to 7 colliculi connected with each pillar, forming hexactinellid networks, 0.05 to 0.15 in width (Fig. 8.3). Mamelons are observed as concentric circles, at a distance of 3 to 5 mm from one to another. Microstructure is compact.

### Material

10 specimens from the Middle Devonian of Teignmouth (NHMUK PI P5538, P5940), in South Devon (UK) and Sötcnich (NHMUK PI P5535, P5540, H3243), Gerolstein (NHMUK PI P5536), Büchel (NHMUK PI P5537, P5539, P5544), Borbach (NHMUK PI P5545), in Germany. *Remarks*.—This species is different from other species in having short but superposed pillars, prominent colliculi and mamelons.

Genus *Bifariostroma* Khalfina, l968

*Type species*.—*Actinostroma bifarium* Nicholson, 1889.

*Bifariostroma bifarium* (Nicholson, 1889) Figure 8.4–8.6

1886b *Actinostroma bifarium* Nicholson, p. 231, pl. 6, figs. 4, 5.

1889 *Actinostroma bifarium*; Nicholson, p. 136, pl. 13, figs. 3–7.

1971 *Actinostroma bifarium*; Kaźmierczak, p. 135, pl. 38, figs. 2, 3; pl. 39, figs. 1, 2.

1971 *Actinostroma* cf. *bifarium*; Zukalová, p. 38, pl. 6, figs. 3–5.

1995 *Actinostroma bifarium*; Krebedünkel, p. 32, pl. 2, figs. 1, 2.

1999 *Bifariostroma bifarium*; Stock in Stearn et al., p. 33.

2015 *Bifariostroma bifarium*; Stock, p. 770, fig. 426.

2016 *Bifariostroma bifarium*; Wolniewicz, p. 240, figs. 10D, 11A, B.

**Figure 10.**
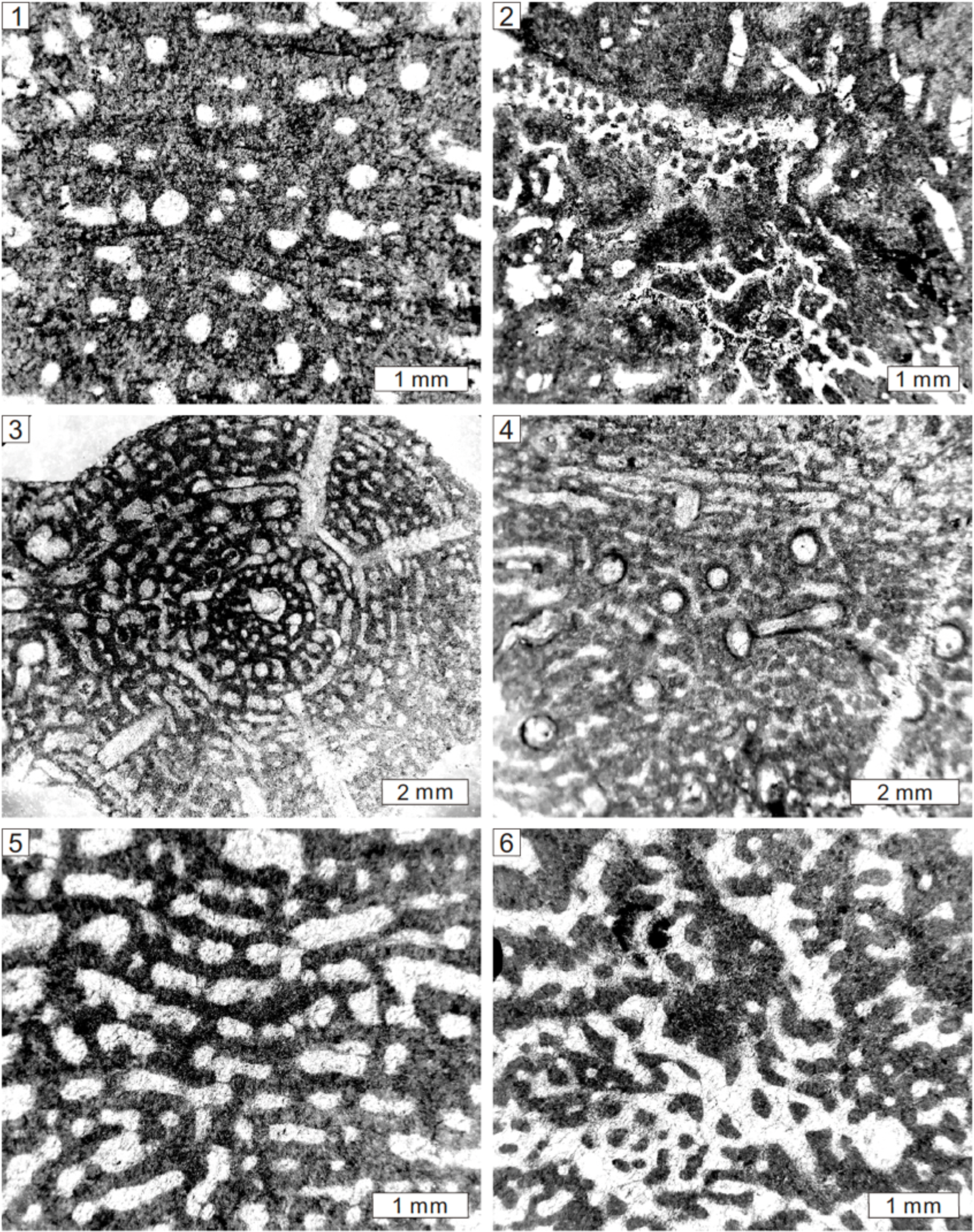
Longitudinal and tangential sections of *Clathrocoilona spissa* (Lecompte, 1951) from the Middle Devonian of Gerolstein, Germany, NHMUK PI P5735 (1, 2), *Dendrostroma oculatum* (Nicholson, 1886a) from the Middle Devonian of Büchel, Germany, NHMUK PI P6073, (3, 4; this sample is the one used by Stearn, 2015b, Fig. 4.36), *Stictostroma damnoniensis* (Nicholson, 1886b) from the Middle Devonian of Teignmouth, South Devon, UK, NHMUK PI P6042 (5, 6).

### Holotype

Specimen (NHMUK PI P5639) from the Middle Devonian of Büchel, Germany (Nicholson, 1886b, pl. 6, figs. 4, 5).

### Description

External forms appear to be laminar, but were prepared as fragments. The skeleton is composed of regular networks. In longitudinal section, pillars dominate, encompassing two types, some vertically extensive, thick, spacing 4 to 5 per 2 mm, 0.2 to 0.5 mm in width (Fig. 8.4). Others are short, post-like or spool-shaped, commonly confined to one interlaminar space, locally superposed across laminae, spacing about 5 to 7 per 2 mm and 0.05 to 0.15 mm in width. Colliculi are thin, spacing about 6 to 8 per 2 mm and 0.05 to 0.15 mm in thickness. Galleries are rectangular. In tangential section, two types of pillars are round in the interlaminar spaces, but show radiating colliculi across spaces, forming complex interconnected networks (Fig. 8.5, 8.6). Microstructure is compact.

### Material

Five specimens from the Middle Devonian of Teignmouth (NHMUK PI P5636, P5641, P5642), in South Devon (UK) and Büchel (NHMUK PI P5639, P5645), in Germany.

### Remarks

This species was previously identified as *Actinostroma bifarium*, but it shows a high resemblance to *Bifariostroma* in possessing two types of radiating pillars, which is different from the one type radiating pillar of *Actinostroma*. Therefore, it was revised to *Bifariostroma* (Stock, 2015; Wolniewicz, 2016).

Order Stromatoporellida Stearn, 1980 Family Stromatoporellidae Lecompte, 1951

Genus *Stromatoporella* Nicholson, 1886a

*Type species*.—*Stromatopora granulata* Nicholson, 1873.

*Stromatoporella arachnoidea* Nicholson, 1886b Figure 9.1, 9.2

1886b *Stromatoporella arachnoidea* Nicholson, p. 237, pl. 8, figs. 1, 2.

### Holotype

Specimen (NHMUK PI P6063) from the Middle Devonian of Büchel, Germany (Nicholson, 1886b, pl. 8, figs. 1, 2).

### Description

External forms are laminar. The skeleton consists of short pillars and laterally long laminae. In longitudinal section, pillars are short, spool-shaped, confined to interlaminar spaces, not superposed, spacing about 6 to 7 per 2 mm and 0.1 to 0.5 mm in width (Fig. 9.1). Laminae are thin, horizontally continuous, slightly undulating, spacing about 8 to 9 per 2 mm and 0.05 to 0.1 mm in thickness. Galleries are round or laterally elongate, with abundant oblique dissepiments. In tangential section, pillars are commonly isolated or irregular. Ring pillars rarely occur (Fig. 9.2). Laminae are undulating or serve as concentric circles in the mamelons. Microstructure is fibrous or cellular.

### Material

Two specimens from the Middle Devonian of Büchel (NHMUK PI P6063) and Gerolstein (NHMUK PI P6064), in Germany.

### Remarks

The rare occurrence of ring pillars in this species derives from upward branching pillars, and it is different from typical ring pillars of *S*. *granulata*, which are derived from upward inflected laminae.

*Stromatoporella solitaria* Nicholson, 1892 Figure 9.3, 9.4

1892 *Stromatoporella solitaria* Nicholson, p. 210, pl. 7, fig. 4; pl. 27, figs 4–7; pl. 2, figs. 9–10.

1952 *Stromatoporella solitaria*; Lecompte, p. 173, pl. 23, figs. 6, 7.

### Holotype

Specimen (NHMUK PI P6035) from the Middle Devonian of Gerolstein, Germany (Nicholson, 1892, pl. 27, figs 4–7).

### Description

Growth forms of illustrated specimens are fragmented, but Nicholson (1892) noted the laminar shapes. The skeleton consists of regular networks formed by short pillars and laterally long laminae. In longitudinal section, pillars are short, spool-shaped, inflecting upward and downward to laminae, with central tubes, confined to interlaminar spaces, not superposed, spacing about 5 to 7 per 2 mm and 0.05 to 0.3 mm in width (Fig. 9.3).

Laminae are thin, locally interrupted, slightly undulating, tripartite with central light zone and surrounding dark zones, spacing about 6 to 7 per 2 mm and 0.1 to 0.2 mm in thickness. Galleries are round or laterally elongate, with sparse oblique dissepiments. Mamelons develop, 4 mm in height. In tangential section, pillars are commonly isolated or irregular. Ring pillars are abundant (Fig. 9.4). Laminae are undulating or concentric around mamelons. Microstructure is ordinicellular, tripartite or cellular.

### Material

10 specimens from the Middle Devonian of Gerolstein (NHMUK PI P6034, P6035, P6036, P6037, P6038, P6039, P6040, P6047, P6048), in Germany and Teignmouth (NHMUK PI P6041), in South Devon (UK).

### Remarks

Abundant ring pillars in the present species are formed by spool-shaped hollow pillars, which is different from ring pillars of *S*. *granulata* by upward inflected laminae.

Genus *Clathrocoilona* Yavorsky, 1931

*Type species*.—*Clathrocoilona abeona* Yavorsky, 1931.

*Clathrocoilona curiosa* (Bargatzky, 1881) Figure 9.5, 9.6

1881 *Stromatopora curiosa* Bargatzky, p. 285.

1886c *Stromatoporella curiosa*; Nicholson, p. 8, pl. 1, figs. 1–3.

1892 *Stromatoporella curiosa*; Nicholson, p. 213, pl. 28, figs. 1–3.

1995 *Stromatoporella curiosa*; Krebedünkel, p. 93, pl. 12, figs. 5, 6.

1997 *Clathrocoilona* (*Clathrocoilona*) *curiosa*; Avlar and May, p. 109, pl. 1, fig. 2.

2008 *Clathrocoilona curiosa*; Salerno, p. 66, pl. 10, fig. 1.

### Holotype

Specimen from the Middle Devonian of Eifel, Germany (Bargatzky, p. 285).

### Description

Growth forms are thin laminar, locally encrusting solitary rugose corals. The skeleton is composed of regular networks formed by short pillars and extensive laminae. In longitudinal section, pillars are post-like, confined to interlaminar spaces, rarely superposed, spacing about 5 to 8 per 2 mm and 0.1 to 0.2 mm in width (Fig. 9.5). Laminae are thin, long, tripartite with a central light line and surrounding dark zones, spacing about 5 to 7 per 2 mm and 0.1 to 0.2 mm in thickness. Galleries are subrectangular, round, or laterally elongated. In tangential section, pillars are round, and isolated (Fig. 9.6). Laminae are undulating or concentric. Mamelon locally occurs. Microstructure is tripartite or ordinicellular.

### Material

Four specimens from the Middle Devonian of Büchel (NHMUK PI P6051, P6052, P6094) and Gerolstein (NHMUK PI P6054), in Germany.

### Remarks

This species was originally assigned in *Stromatoporella* by Nicholson (1886b), but no ring pillar is observed. The thin, encrusting skeletons and tripartite laminae, however, closely resemble *Clathrocoilona*.

*Clathrocoilona spissa* (Lecompte, 1951) Figure 10.1, 10.2

1951 *Stromatoporella spissa* Lecompte, p. 187, pl. 27, figs. 1–4.

1971 *Stromatopora spissa*; Kaźmierczak, p. 92, pl. 21, fig. 2.

1971 *Clathrocoilona spissa*; Zukalová, p. 56, pl. 15, figs. 1, 2.

1974 *Clathrocoilona spissa*; Flügel, p. 165, pl. 24, 26, 27.

1980 *Clathrocoilona spissa*; Mistiaen, p. 196, pl. 7, figs. 3–9.

1984 *Clathrocoilona spissa*; Cockbain, p. 25, pl. 11.

1985 *Clathrocoilona spissa*; Mistiaen, p. 96–101, pl. 6, figs. 6–8.

1988 *Clathrocoilona spissa*; Mistiaen, p. 174.

1993 *Clathrocoilona* (*Clathrocoilona*) *solidula spissa*; May, p. 38, pl. 7, fig. 2; pl. 8, figs. 1, 2.

1999 *Clathrocoilona* (*Clathrocoilona*) *solidula spissa*; May, p. 127.

2005 *Clathrocoilona* (*Clathrocoilona*) *solidula spissa*; May, p. 180, pl. 13, figs. 2, 3.

2008 *Clathrocoilona spissa*; Salerno, p. 74, pl. 11, fig. 2.

2022 *Clathrocoilona spissa*; Huang *et al*., p. 718, fig. 6.

### Holotype

Specimen (Surice 18, n° 7164) from the Middle Devonian of Dinant Basin, Belgium (Lecompte, 1951, pl. 27, fig. 1).

### Description

Growth forms are uncertain due to fragmented specimens. The skeleton consists of densely packed thick pillars and laminae. In longitudinal section, skeletons are varied. In some areas, laminae and pillars can hardly be differentiated. Pillars and laminae are thick, and irregularly amalgamated. Gallery shapes are irregular, locally with abundant dissepiments. In the other areas, skeletons comprise regular laminae and pillars (Fig. 10.1). Pillars are spool-shaped, confined to interlaminar spaces, spacing 4 to 5 per 2 mm, 0.1 to 1 mm in width. Laminae are thick, locally with dark axial lines and the coated light zones, spacing 4 per 2 mm, 0.2 to 0.5 mm in thickness. In tangential section, skeletons are amalgamated, with local pillars round (Fig. 10.2), subtriangular, or irregular, isolated, ranging from 0.1 to 1 mm in diameter. Astrorhizal canals are common, 0.1 to 0.2 mm in width, with sparse dissepiments. Microstructure is compact or tripartite.

### Material

Four specimens from Middle Devonian of Büchel (NHMUK PI P6053), Gerolstein (NHMUK PI P5735), and Sötenich (NHMUK PI P5736, P6091) in Germany.

### Remarks

This species is different from *C*. *curiosa* in having irregular skeletons formed by densely packed, thick pillars and laminae. It is revised from specimens previously assigned to *Stromatoporella damnoniensis* and *S*. *curiosa* by Nicholson (1886b, 1892).

Genus *Dendrostroma* Lecompte, 1952

*Type species*.—*Idiostroma oculatum* Nicholson, 1886a.

*Dendrostroma oculatum* (Nicholson, 1886a) Figure 10.3, 10.4

1886a *Idiostroma oculatum* Nicholson, p. 101, figs. 14, 15.

1892 *Idiostroma oculatum*; Nicholson, p. 225, pl. 29, figs. 10, 11.

**Figure 11.**
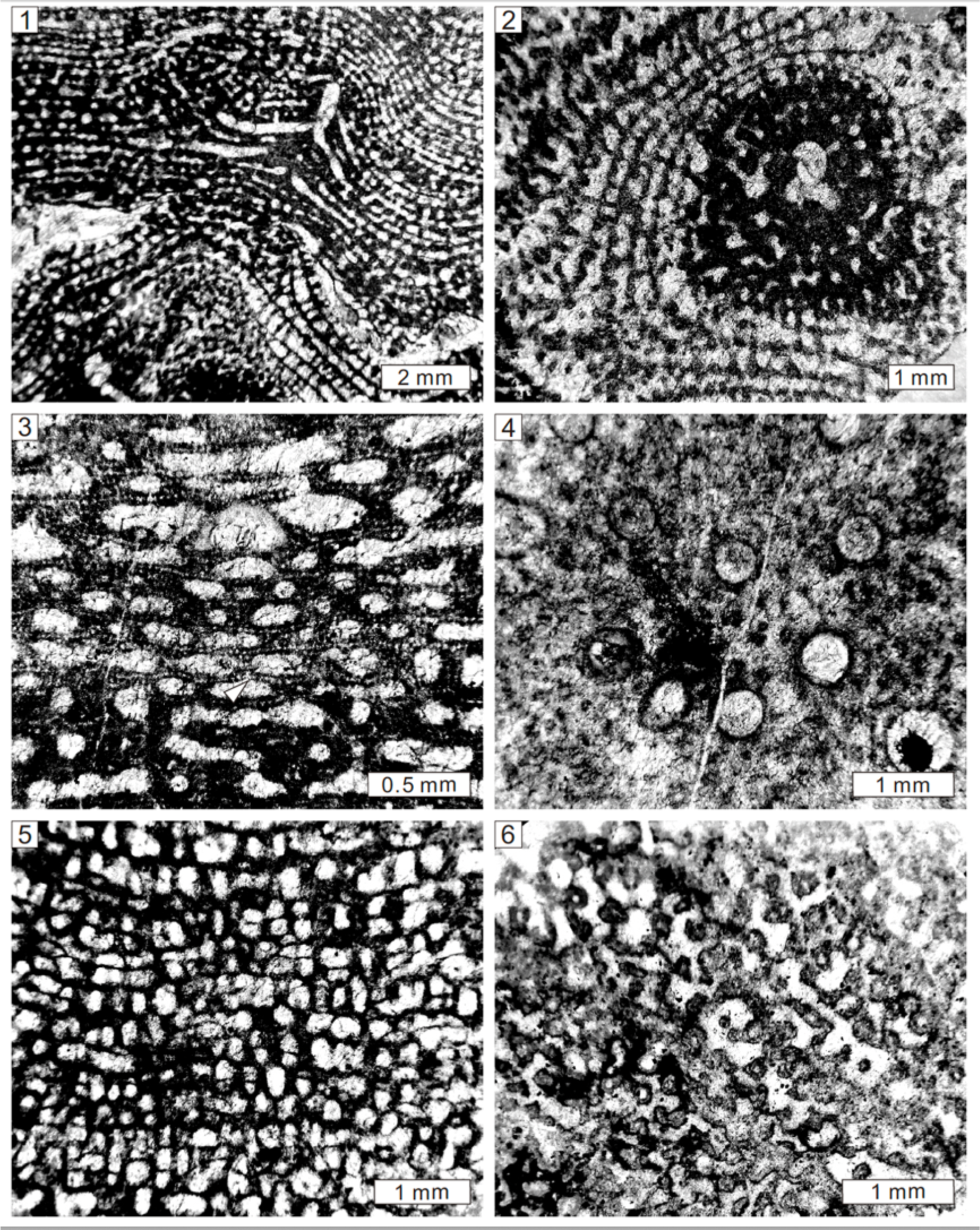
Longitudinal and tangential views of *Stictostroma saginatum* (Lecompte, 1951) from the Middle Devonian of Teignmouth, South Devon, UK, NHMUK PI P5699 (1, 2), *Stictostroma socialis* (Nicholson, 1892) from the Middle Devonian of Gerolstein, Germany, NHMUK PI P6095 (3, 4), *Tubuliporella calamosa* (Ripper, 1933) from the Lower Devonian of Mitchell’s Quarry, Cave Hill, Lilydale, Australia, NHMUK PI H3898 (5, 6). Note the ordinicellular microstructure (white arrow).

1952 *Dendrostroma oculatum*; Lecompte, p. 320, pl. 61. fig. 1.

2015b *Dendrostroma oculatum*; Stearn, p. 784, fig. 436a, b.

### Holotype

Specimen (NHMUK PI P6073) from the Middle Devonian of Büchel, Germany (Nicholson, 1886a, figs. 14, 15).

### Description

Growth forms are dendroid, in diameter of 7 to 10 mm. In longitudinal section, the skeleton consists of short pillars and arched laminae. In the central zone, skeletons are featured by an axial tube, 0.5 to 0.8 mm in diameter. In the peripheral zone, skeletons are well differentiated (Fig. 10.3). Pillars are short, postlike or spool-shaped, confined to interlaminar spaces, not superposed, spacing 4 to 5 per 2 mm and 0.1 to 0.5 mm in width. Laminae are laterally extensive, thick, slightly wavy, locally tripartite with central light line and surrounding dark zones, spacing 8 to 9 per 2 mm and 0.1 to 0.3 mm in thickness. Galleries are elongate. Tabulate coral symbionts are common, in width of 0.5 mm. In tangential section, networks are formed by short pillars and concentric laminae. The central zone presents an axial tube, with coral symbionts spread outward. In oblique section, pillars are isolated (Fig. 10.4).

The microstructure is compact to fibrous.

### Material

Two specimens from the Middle Devonian of Büchel (NHMUK PI P6073, H4777) in Germany and one from the Devonian of Dartington (NHMUK PI P6115) in South Devon (UK).

### Remarks

This species was previously assigned to *Idiostroma* by Nicholson (1886a, 1892), but the occurrence of isolated and confined pillars is distinguished by vermiform and superposed pachysteles of *Idiostroma*.

Genus *Stictostroma* Parks, 1936

*Type species*.—*Stictostroma gorriense* Stearn, 1995.

*Stictostroma damnoniensis* (Nicholson, 1886b) Figure 10.5, 10.6

1886b *Stromatoporella damnoniensis* Nicholson, p. 237, pl. 8, figs. 3, 4.

1892 *Stromatoporella damnoniensis*; Nicholson, p. 207, pl. 27, figs. 8, 9.

### Holotype

Specimen (NHMUK PI P6042) from the Middle Devonian of Teignmouth in South Devon, UK (Nicholson, 1886b, pl. 8, figs. 3, 4).

### Description

External form is unclear but Nicholson (1886b) noted domical or bulbous shapes. The skeleton is composed of networks formed by short pillars and intermittent laminae. In longitudinal section, pillars are post-like or spool-shaped, confined to interlaminar spaces, not superposed, spacing about 4 to 6 per 2 mm and 0.1 to 0.4 mm in width. Laminae are long but interrupted, spacing about 6 to 8 per 2 mm and 0.1 to 0.2 mm in thickness (Fig. 10.5). Galleries are round, irregular, or laterally elongated. In tangential section, pillars are round, isolated (Fig. 10.6). Laminae are undulating or concentric, with circular holes inside. Mamelons locally occur, 8 mm in width. Microstructure is fibrous and coarsely ordinicellular.

### Material

One specimen from the Middle Devonian of Teignmouth (NHMUK PI P6042) in South Devon (UK).

### Remarks

This specimen was assigned to *Stromatoporella* due to the occurrence of possible ring pillars, but these circular skeletons are a part of laminae (Fig. 10.5, 10.6). Lecompte (1951) revised the paratype of this species into *Clathrocoilona crassitexta*. The holotype from UK, however, shows more relationships with *Stictostroma* due to coarsely ordinicellular microstructure.

*Stictostroma saginatum* (Lecompte, 1951) Figure 11.1, 11.2

1951 *Stromatoporella saginata* Lecompte, p. 171, pl. 22, figs. 5–7; pl. 23, figs. 1–3.

1983 *Clathrocoilona* cf. *saginata*; Stearn, p. 549, fig. 5G–H.

1984 *Clathrocoilona saginata*; Cockbain, p. 25, pl. 10, figs. A–D.

1985 *Stictostroma saginatum*; Mistiaen, p. 115, pl. 8, figs. 6–11.

1988 *Stictostroma saginatum*; Mistiaen, p. 176, pl. 21, figs. 5–9.

1999 *Stictostroma saginatum*; Mistiaen, p. 36, pl. 3, figs 9–12; pl. 4, figs. 1, 2.

**Figure 12.**
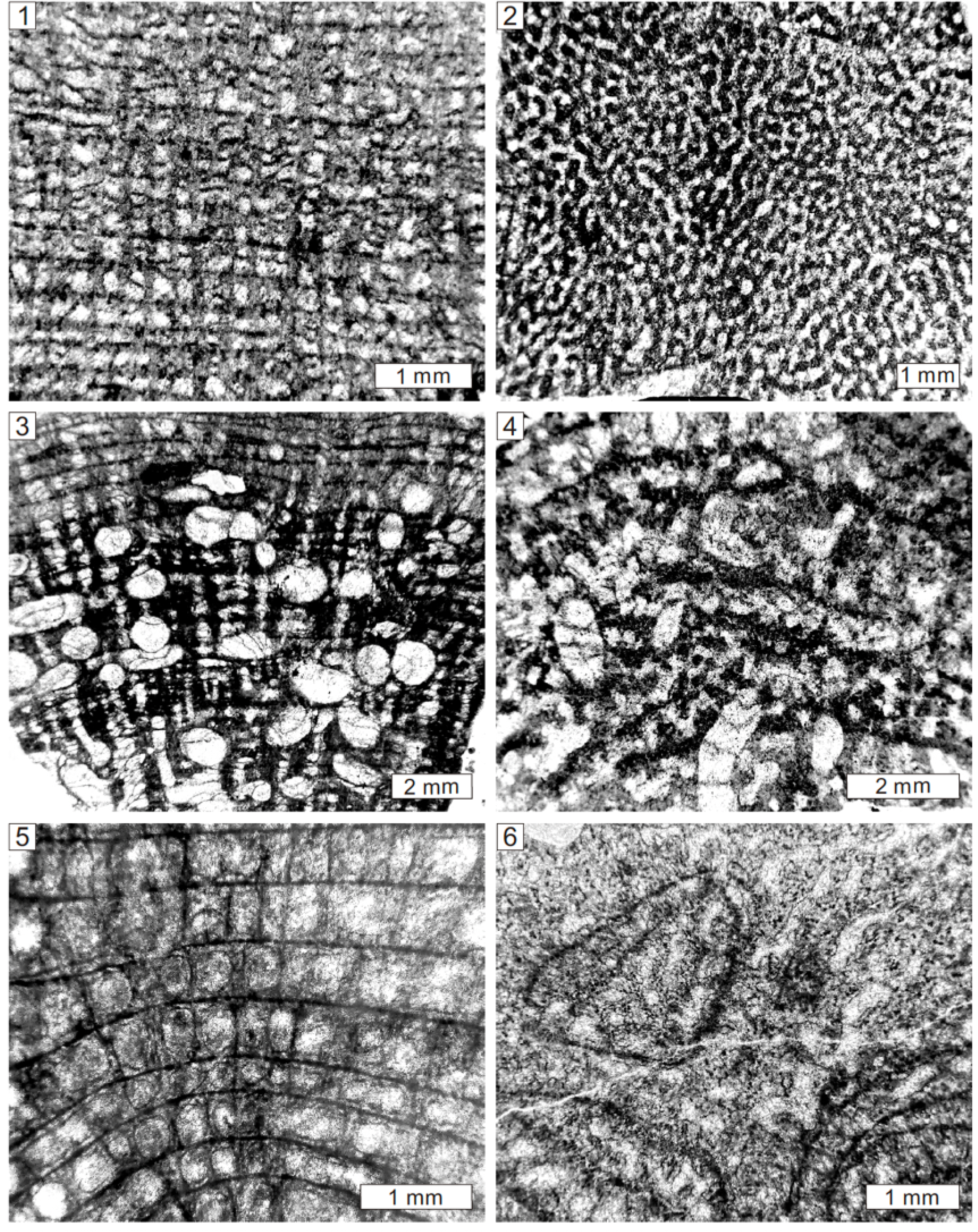
Longitudinal and tangential sections of *Trupetostroma datingtonense* (Carter, 1880) from the Middle Devonian of Dartington, South Devon, UK, NHMUK PI P5746 (1, 2), *Trupetostroma thomasi* Lecompte, 1952 from the Middle Devonian of Hebborn, Germany, NHMUK PI P5934, (3, 4), *Hermatostroma episcopale* Nicholson, 1892 from the Middle Devonian of Bishopsteignton, South Devon, UK, NHMUK PI P6065 (5, 6).

2000 *Stictostroma saginatum*; Mistiaen & Gholamalian, p. 83–84, pl. 6, figs. 6–8.

2008 *Stictostroma saginatum*; Salerno, p. 82–84, pl. 12, fig. 2.

2022 *Stictostroma saginatum*; Huang *et al*., p. 720, fig. 7A, B.

### Holotype

Specimen (Rance 8275, n° 7502) from the Upper Devonian of Dinant Basin, Belgium (Lecompte, 1951, pl. 23, fig. 1).

### Description

Growth forms are unclear due to limited preservation of specimen. The skeleton consists of short pillars and undulating laminae. In longitudinal section, pillars are spool-shaped, confined to interlaminar spaces, locally superposed, spacing about 7 to 8 in 2 mm, 0.1 to 0.2 mm in width (Fig. 11.1). Laminae are laterally extensive, undulating especially around mamelons, spacing 8 to 10 per 2 mm, 0.05 to 0.2 mm in thickness. Galleries are circular, or laterally elongated. Mamelons are conspicuous, generally 4 mm in height, with central and branching astrorhizal canals. In tangential section, pillars are round, isolated. Laminae are undulating or present as concentric circles (Fig. 11.2). Mamelons are apparent, with astrorhizal canals, laterally separated by distances of 2 to 4 mm. The microstructures are compact or ordinicellular.

### Material

Four specimens from the Middle Devonian of Teignmouth (NHMUK PI P5482, P5699, P5874, P5949), in South Devon (UK).

### Remarks

This specimen was assigned to *Stromatoporella socialis*, but ring pillars have not been observed. Lecompte (1951) revised this species into *Stromatoporella saginata*, which is widely accepted as *Stictostroma saginatum*. We therefore reassign it to *Stictostroma*.

*Stictostroma socialis* (Nicholson, 1892) Figure 11.3, 11.4

1892 *Stromatoporella socialis* Nicholson, p. 206, pl. 26, figs. 5–7.

1952 *Stromatoporella socialis*; Lecompte, p. 163, pl. 21. figs. 2, 3.

### Holotype

Specimen (NHMUK PI P6043) from the Middle Devonian of Teignmouth in South Devon, UK (Nicholson, 1892, pl. 26, figs. 5–7).

### Description

External forms appear to be laminar, in thickness of 1 to 2 cm. The skeleton consists of short pillars and long laminae. In longitudinal section, pillars are spool-shaped, confined to interlaminar spaces, not superposed, spacing 5 to 9 in 2 mm, 0.1 to 0.3 mm in width. Laminae are laterally extensive, slightly undulating, tripartite with central hollow cavities (Fig. 11.3), spacing 12 to 14 per 2 mm, 0.05 to 0.2 mm in thickness. Galleries are circular, or laterally elongated. Syringoporid tabulate corals are observed, 0.5 to 1 mm in width. In tangential section, pillars are round, isolated (Fig. 11.4). The microstructures are ordinicellular.

### Material

15 specimens from the Middle Devonian of Teignmouth (NHMUK PI P5696, P5700, P5701, P5703, P6043, P6044, P6045, P6046), Dartington (NHMUK PI P5702), in South Devon (UK); Büchel (NHMUK PI P6094); and Gerolstein (NHMUK PI P6049, P6050, P6092, P6093, P6095), in Germany.

### Remarks

The current species was identified as *Stromatoporella* by Nicholson (1892), but the ordinicellular microstructure and absence of ring pillars justify its assignment to *Stictostroma*.

Genus *Tubuliporella* Khalfina, 1968

*Type species*.—*Tubuliporella lecompti* Khalfina, 1968.

*Tubuliporella calamosa* (Ripper, 1933) Figure 11.5, 11.6

1933 *Clathrodictyon calamosum* Ripper, p. 160, fig. 6E, F.

1968 *Clathrodictyon calamosum*; Fliigel and Flugel-Kahler, p. 56.

1993 *Tubuliporella calamosa*; Webby et al., p. 148, figs. 16F, 17A–D.

### Holotype

Specimen (NHMUK PI H3898) from the Lower Devonian of Lilydale Limestone, Mitchell’s, Cave Hill, Lilydale, Victoria, Australia (Ripper, 1933, fig. 6E, F).

### Description

Growth forms are unclear due to poorly preserved specimens. The skeleton commonly shows networks formed by short pillars and laterally extensive laminae. In longitudinal section, skeleton is dominated by ring pillars (Fig. 11.5). Ring pillars are locally superposed, forming vertical channels filled by thin laminae or dissepiments,0.2 to 0.4 mm in width. Laminae are weakly developed, thin, spacing about 7 to 9 per 2 mm and 0.05 to 0.2 mm in thickness. Galleries are round. The tangential section presents ring pillars (Fig. 11.6), in diameter of 0.1 to 0.2 mm. Laminae are inconspicuous.

Microstructure is compact.

### Material

One specimen from the Lower Devonian of Mitchell’s Quarry (NHMUK PI H3898), Cave Hill, Lilydale, Victoria (Australia).

### Remarks

The current species was assigned to *Clathrodictyon* by Ripper (1933) and Flügel and Flugel-Kahler (1968), but the superposed ring pillars justify its assignment as *Tubuliporella* (Webby et al., 1993). It resembles *Stromatoporella*, but differs from the latter genus in irregular skeletons, superposed ring pillars, and compact microstructure.

Family Trupetostromatidae Germovsek, 1954 Genus *Trupetostroma* Parks, 1936

*Type species*.—*Trupetostroma warreni* Parks, 1936.

*Trupetostroma dartingtonense* (Carter, 1880) Figure 12.1, 12.2

1880 *Stromatopora dartingtonensis* Carter, p. 346, pl. 18, figs. 1–5.

1891 *Parallelopora dartingtonensis*; Nicholson, p. 199, pl. 4, fig. 1; pl. 24, figs. 13–15.

**Figure 13.**
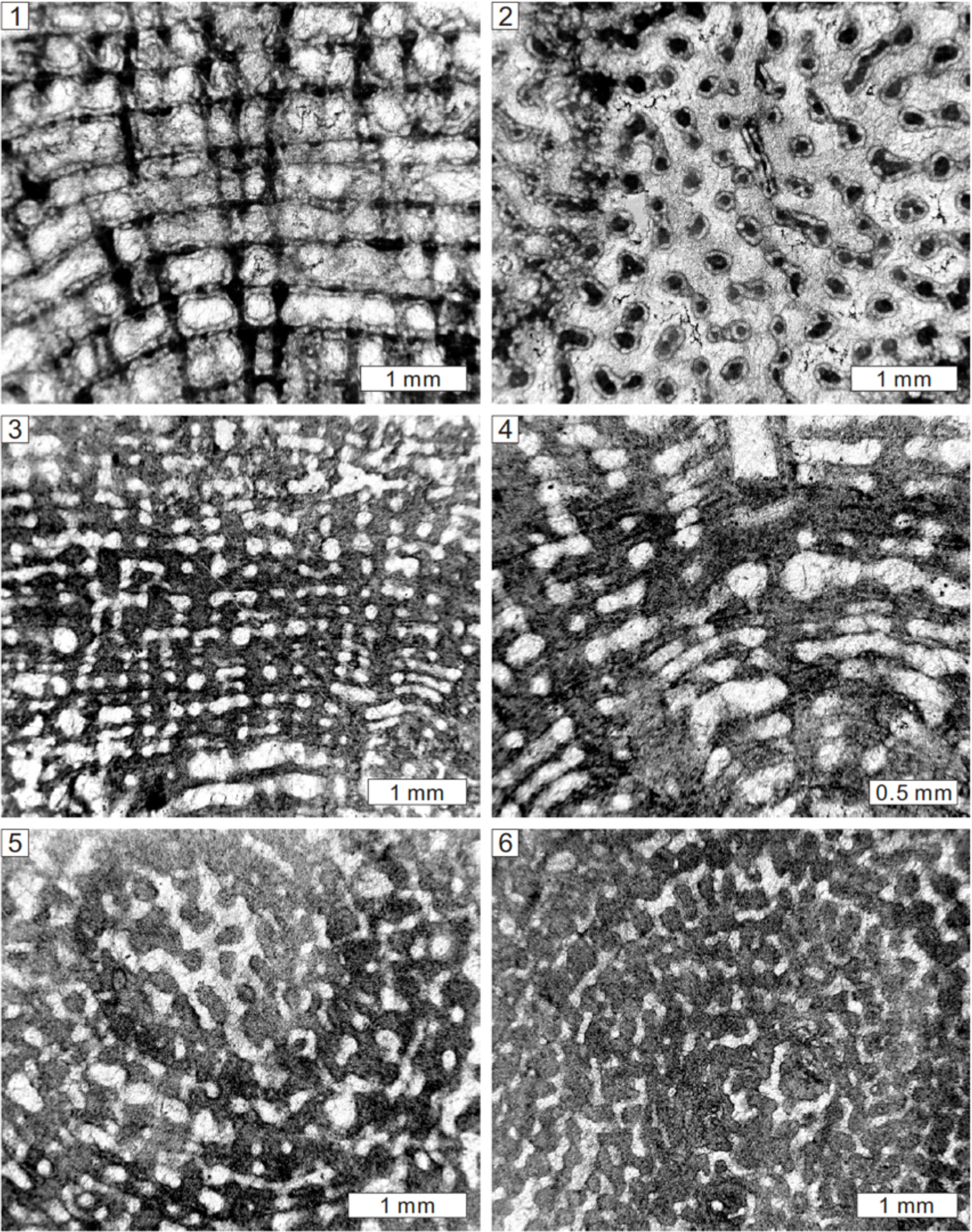
Longitudinal and tangential views of *Hermatostroma schlueteri* Nicholson, 1886a from the Middle Devonian of Hebborn, Germany, NHMUK PI P5527 (**1, 2**), *Hermatostromella holmesae* (Webby et al., 1993) from Lower Devonian of Mitchel’s Quarry, Cave Hill, Lilydale, Victoria, Australia, NHMUK PI H3896 (**3**–**6**).

**Figure 14.**
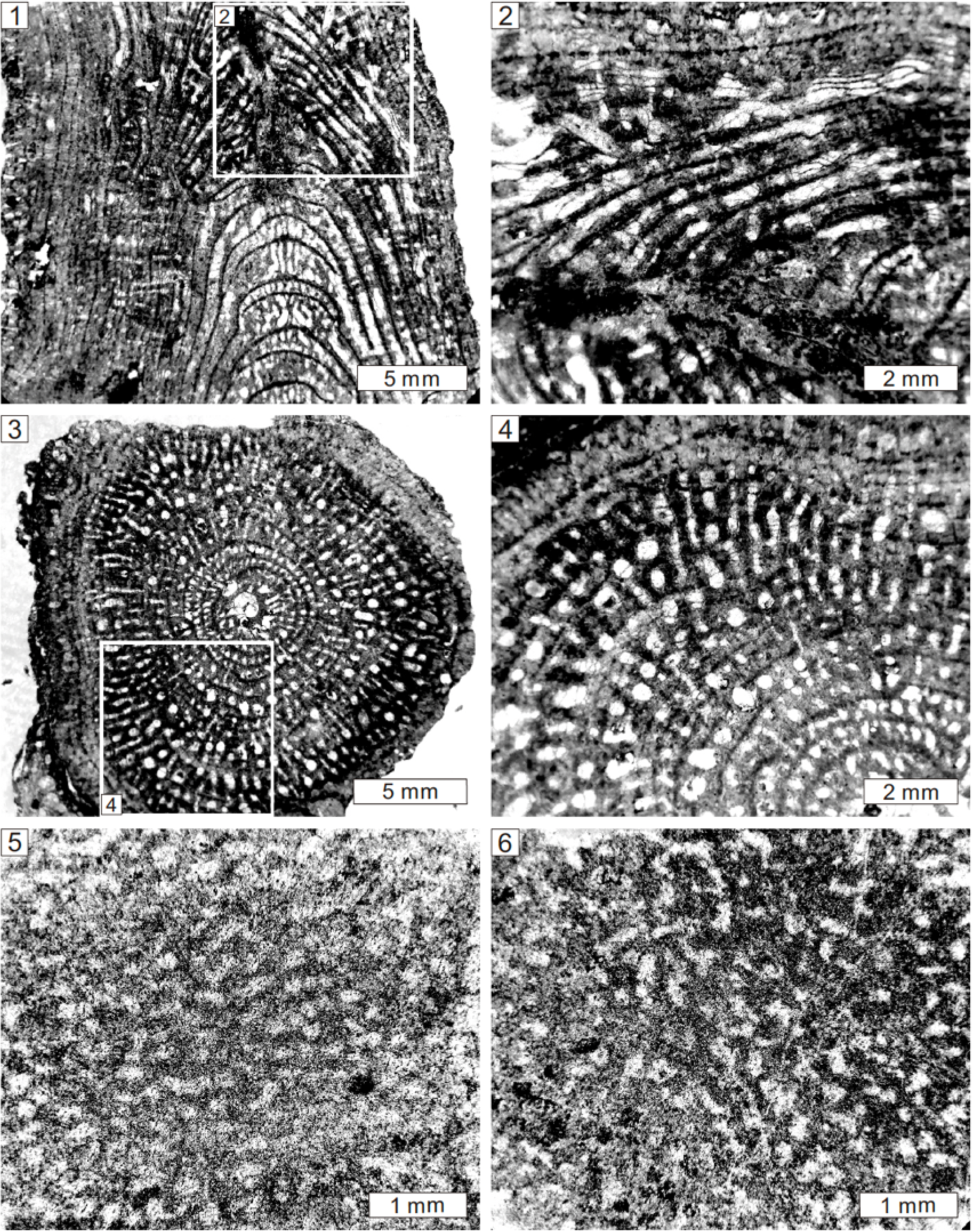
Longitudinal and tangential sections of *Idiostroma roemeri* Nicholson, 1886a from the Middle Devonian of Hebborn, Germany, NHMUK PI P6076 (**1**–**4**) *Stromatopora aff. polaris* (Stearn, 1983) from the Lower Devonian of Buchan, Victoria, Australia, NHMUK PI H3905 (**5, 6**).

**Figure 15.**
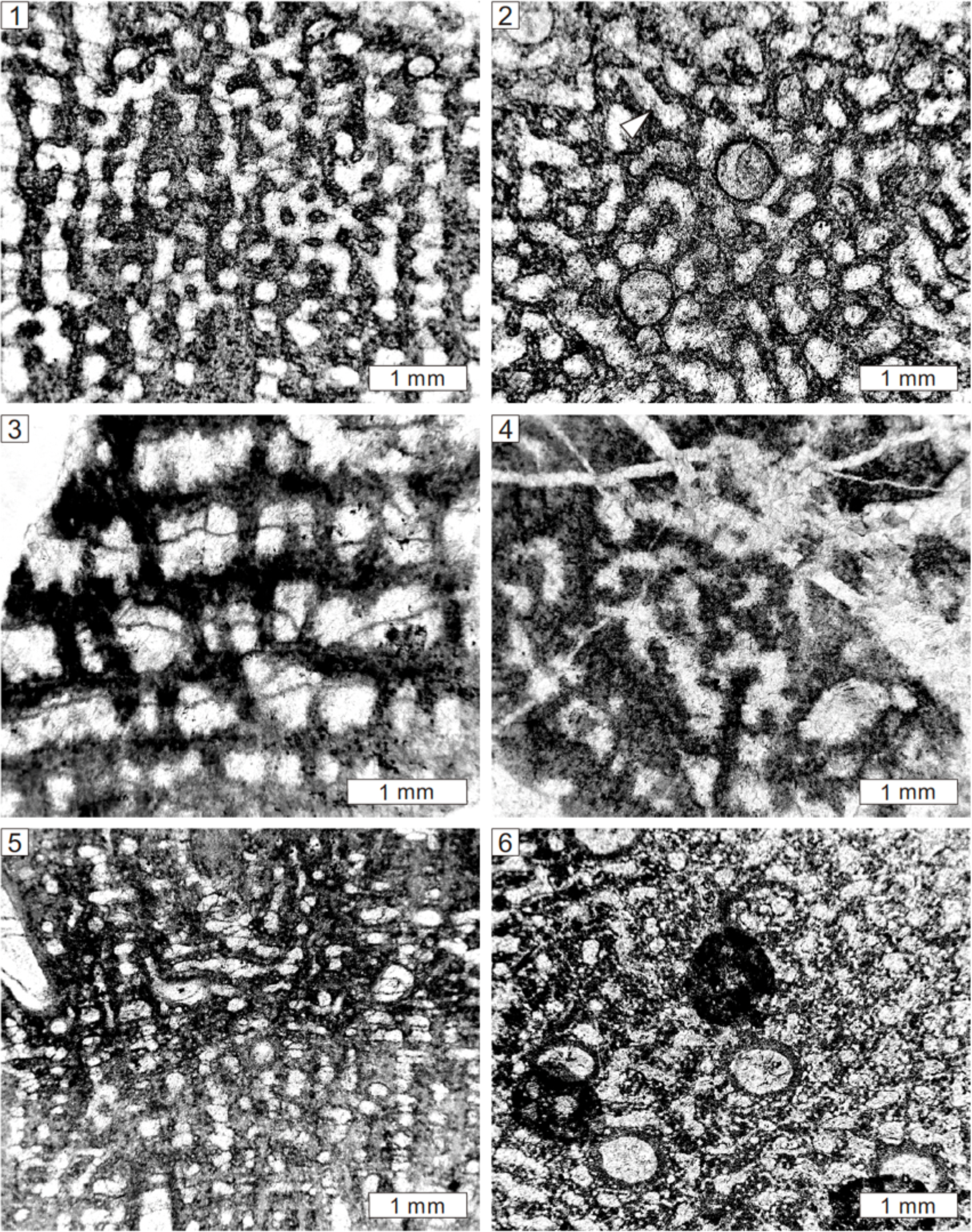
Longitudinal and tangential views of *Stromatopora hüpschii* (Bargatzky, 1881) from Middle Devonian of Büchel, Germany, NHMUK PI P5488 (**1, 2**), *Pseudotrupetostroma buchanense* (Ripper,1937a) from Lower Devonian of Buchan, Victoria, Australia, NHMUK PI H3906 (**3, 4**), *Pseudotrupetostroma porosum* (Yang and Dong, 1979) from the Middle Devonian of Teignmouth, South Devon, UK, NHMUK PI P5933 (**5, 6**). Note the allotubes (white arrows).

1952 *Parallelopora* cf*. dartingtonensis*; Lecompte, p. 295, pl. 49, fig. 3.

### Holotype

Specimen from the Middle Devonian of Pit-Park Quarry, Dartington, South Devon, UK (Carter, 1880, pl. 18, figs. 1–5).

### Description

Nicholson (1891) noted bulbous, domical, or laminar external forms. The skeleton shows regular networks formed by long pillars and laminae. In longitudinal section, pillars are post-like, vertically continuous, passing through dozens of laminae (Fig. 12.1), spacing 8 to 9 per 2 mm and 0.1 to 0.2 mm in width. Laminae are laterally long, tripartite with a central dark line and surrounding light zones, spacing about 7 to 8 per 2 mm and 0.05 to 0.2 mm in thickness. Galleries are circular, with abundant dissepiments. In tangential section, pillars are circular, isolated, locally connected around laminae (Fig. 12.2). Astrorhizal canals are sparse. Microstructure is compact. *Material*.—15 specimens from the Middle Devonian of Dartington (NHMUK PI P5740, P5746, P5753, P5937) and Teignmouth (NHMUK PI P5738, P5739, P5741, P5742, P5743, P5744, P5748, P5750, P5752, P5923, P5937), in South Devon (UK).

### Remarks

This species is characterized by regular networks formed by extensive, isolated pillars and tripartite laminae, which closely resemble *Trupetostroma* but differ from *Parallelopora*. Kaźmierczak (1971) revised this species into *Pseudoactinodictyon* based on Polish materials, but the British specimens (Carter, 1880; Nicholson, 1891) show regular networks formed by pillars and laminae.

*Trupetostroma thomasi* Lecompte, 1952 Figure 12.3, 12.4

1891 *Parallelopora capitata*; Nicholson, p. 197, pl. 25, figs. 10–13.

1952 *Trupetostroma thomasi*; Lecompte, p. 240.

### Holotype

Specimen (NHMUK PI P5934) from the Middle Devonian of Hebborn (Paffrath district), Germany (Nicholson, 1891, fig. 27A, B).

### Description

Nicholson (1891) illustrated external skeleton as bulbous form. The skeleton presents regular networks formed by short pillars and thin laminae. In longitudinal section, pillars are spool-shaped, confined to interlaminar spaces but systematically superposed, spacing 3 to 5 per 2 mm and 0.1 to 0.5 mm in width (Fig. 12.3). Laminae are slightly undulating, laterally extensive, tripartite with a central dark zone and surrounding light zones, spacing about 5 to 7 per 2 mm and 0.1 to 0.2 mm in thickness.

Galleries are circular, or laterally elongated. Syringoporid intergrowth rarely develops. Astrorhizal canals are locally abundant, with a width of 10 to 20 mm. In tangential section, skeletons are densely packed. Pillars are circular, isolated. Laminae are undulating (Fig. 12.4). Microstructure is vacuolate, cellular, or compact.

### Material

15 specimens from the Middle Devonian of Teignmouth (NHMUK PI P5520, P5522, P5523, P5556, P5893, P5908, P5935, P6080) in South Devon (UK); Hebborn (NHM UK PI P5524, P5526, P5546, P5934, P6081, P6100) and Steinbreche (NHM UK PI P5549), in Germany. *Remarks*.—This species was originally assigned into *Parallelopora* by Nicholson (1891), but the regular skeletons differ from the dominant interconnected pachysteles and orthoreticular microstructure in the latter genus. It was revised into *Trupetostroma thomasi* by Lecompte (1952), we follow this decision.

Genus *Hermatostroma* Nicholson, 1886a

*Type species*.—*Hermatostroma schlueteri* Nicholson, 1886a.

*Hermatostroma episcopale* Nicholson, 1892 Figure 12.5, 12.6

1892 *Hermatostroma episcopale* Nicholson, p. 219, pl. 28, figs. 4–11.

1952 *Hermatostroma episcopale*; Lecompte, p. 216, pl. 48. fig. 4; pl. 49, figs. 1, 2.

1960 *Hermatostroma episcopale*; Galloway, p. 635, pl. 77, fig. 4.

1963 *Hermatostroma episcopale*; Yang and Dong, p. 162, pl. 10, figs. 3–6.

1979 *Hermatostroma episcopale*; Yang and Dong, p. 68, pl. 32, figs. 7, 8.

1982 *Hermatostroma episcopale*; Dong and Wang, p. 23, pl. 14, figs. 1, 2.

### Holotype

Specimen (NHMUK PI P6065) from the Middle Devonian of Bishopsteignton, South Devon, UK (Nicholson, 1892, pl. 28, figs. 5, 6). *Description*.—Growth forms are uncertain due to limited preservation, but Nicholson (1892) noted bulbous/domical shapes. The skeleton shows regular networks formed by short pillars and continuous laminae. In longitudinal section, pillars are spool-shaped, confined to one interlaminar space, superposed across several laminae (Fig. 12.5), spacing about 3 to 4 per 2 mm and 0.1 to 0.3 mm in width. Laminae are gently wavy, extensive, tripartite with a central dark zone and surrounding light zones, spacing about 4 to 6 per 2 mm and 0.2 to 0.3 mm in thickness. Galleries are circular, 0.1 to 0.5 mm in width. Syringoporid tabulate corals occur, with a width of 1 mm. Dissepiments are common, oblique. In tangential section, pillars are circular, subcircular, isolated or laterally connected. Laminae are circular or undulating. Both pillars and laminae show central dark zone and peripheral light zones (Fig. 12.6). Astrorhizae are well developed, in width of 3 to 4 mm, 5 mm from one to another. Microstructure is compact.

### Material

Six specimens from the Middle Devonian of Teignmouth (NHMUK PI P5686, P5688, P5689, P6086) and Bishopsteignton (NHMUK PI P5692, P6065), in South Devon, (UK).

### Remarks

This species differs from *H*. *schlueteri* in short pillars, common dissepiments, and well-developed astrorhizae.

*Hermatostroma schlueteri* Nicholson, 1886a Figure 13.1, 13.2

1886a *Hermatostroma schlueteri* Nicholson, p. 105, pl. 3, figs. 1, 2.

1892 *Hermatostroma schlueteri*; Nicholson, p. 215, pl. 28, figs. 12, 13.

1952 *Hermatostroma schlueteri*; Lecompte, p. 250, pl. 45, fig. 1.

1957 *Hermatostroma schlueteri*; Galloway and St. Jean, p. 218, pl. 21, fig. 1.

1979 *Hermatostroma schlueteri*; Yang and Dong, p. 69, pl. 37, figs. 1, 2.

1982 *Hermatostroma schlueteri*; Dong and Wang, p. 23, pl. 13, figs. 7, 8.

1984 *Hermatostroma schlueteri*; Cockbain, p. 27, fig. 16A–D.

**Figure 16.**
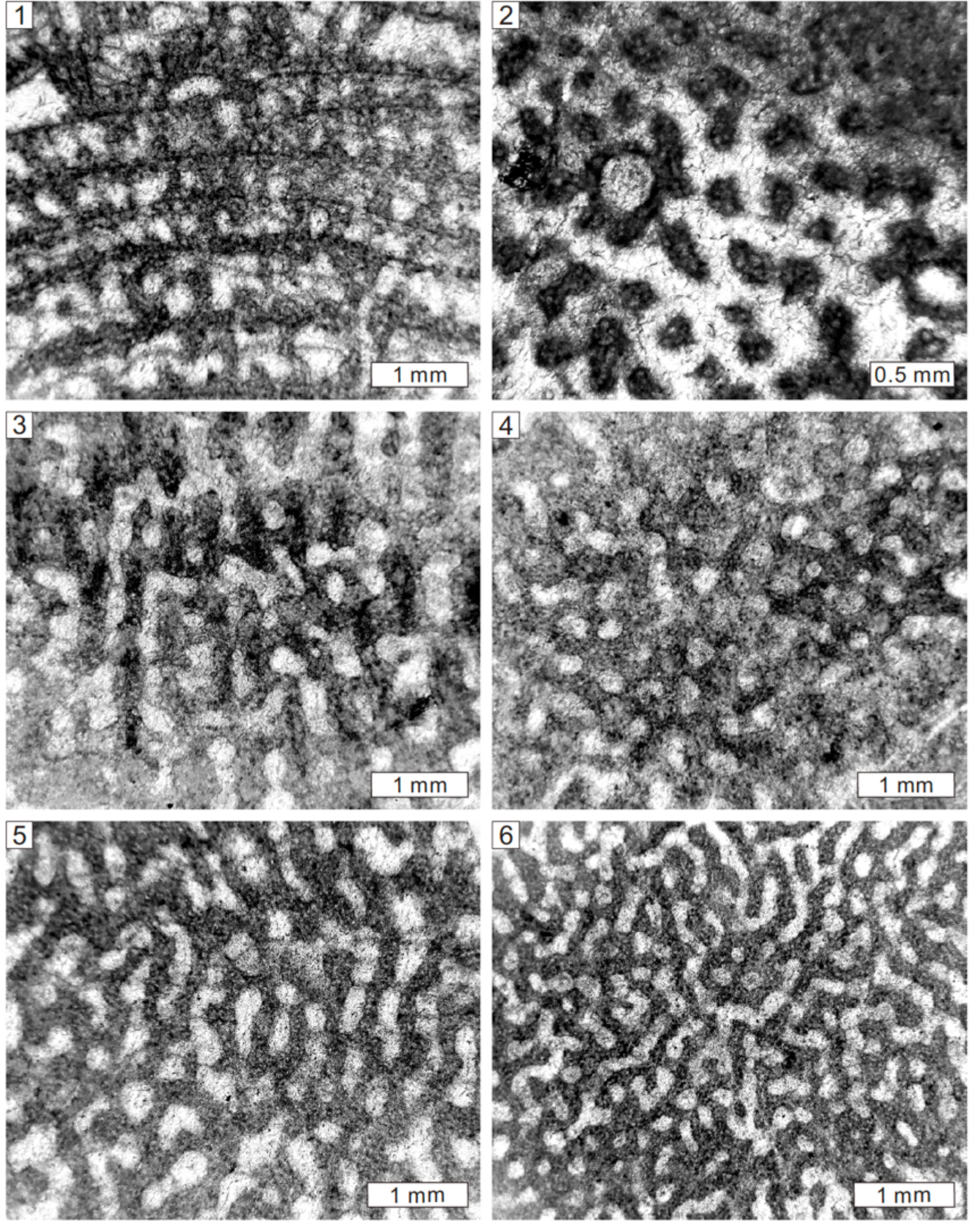
Longitudinal and tangential views of *Pseudotrupetostroma ripperae* Webby and Zhen 1993 from Lower Devonian of Heath’s Quarry, Buchan, Victoria, Australia, NHMUK PI H3916 (**1, 2**), *Neosyringostroma* sp. from the Middle Devonian of Büchel, Germany, NHMUK PI P5530 (**3, 4**), *Syringostromella* cf. *labryrinthea* Stearn, 1990 from Lower Devonian of Heath’s Quarry, Buchan, Victoria, Australia, NHMUK PI H3921 (**5, 6**).

1985 *Hermatostroma schlueteri*; Mistiaen, p.175, pl. 15, figs. 9–11; pl. 16, fig. 1.

1989 *Hermatostroma schlueteri*; Dong et al., p. 267, pl. 8, fig. 3a, b.

1995 *Hermatostroma schlueteri*; Krebedünkel, p. 65, pl. 7, figs. 7, 8.

2008 *Hermatostroma schlueteri*; Salerno, p. 84, pl. 13, figs. 1, 2.

### Holotype

Specimen (NHMUK PI P5527) from the Middle Devonian of Hebborn, Germany (Nicholson, 1886a, pl. 3, figs. 1, 2).

### Description

External forms are uncertain due to poor preservation, but Nicholson (1892) noted bulbous/domical shapes. The skeleton shows regular networks formed by continuous pillars and laminae. In longitudinal section, pillars are long, spool-shaped, confined to one interlaminar space but regularly superposed across dozens of laminae (Fig. 13.1), spacing 6 to 7 per 2 mm and 0.2 to 0.3 mm in width. Laminae are gently wavy, laterally extensive, spacing about 5 to 6 per 2 mm and 0.1 to 0.2 mm in thickness.

Both pillars and laminae are tripartite with a central dark zone and surrounding light zones. Galleries are circular, 0.2 to 1 mm in width. Tubeworms locally occur, with a width of 0.5 to 0.8 mm. In tangential section, pillars are circular or subcircular, isolated, with central dark dots and peripheral light zones or vesicles (Fig. 13.2). Laminae are circular or undulating. Microstructure is compact and vacuolate.

### Material

One specimen from the Middle Devonian of Hebborn (NHMUK PI P5527) in Germany.

### Remarks

The current species is distinguished from *H*. *episcopale* in regular networks formed by long pillars and laminae, absence of astrorhizae, and well-developed tripartite skeletons.

Genus *Hermatostromella* Khalfina, 1961

*Type species*.—*Hermatostromella parasitica* Khalfina, 1961.

*Hermatostromella holmesae* (Webby et al., 1993) Figure 13.3–13.6

1937b *Syringostroma* aff. *ristigouchense*; Ripper, p. 181, pl. 8, figs. 1, 2.

1993 *Amnestostroma holmesae*; Webby et al., 1993, p. 151, figs. 19A–E, 31C–D.

### Holotype

Longitudinal and tangential sections (NMV P141860, P141861, P141862, and P141863) from the Lower Devonian of Lilydale Limestone, Mitchell’s (Cave Hill) Quarry, Victoria, Australia (Webby et al., 1993, fig. 19A– D).

### Description

The skeleton is composed of the regular network formed by pillars and laminae. In longitudinal section, pillars are thick, spool-shaped, confined to interlaminar spaces, but systematically superposed (Fig. 13.3), spacing 7 to 8 per 2 mm and 0.1 to 0.2 mm in width. Laminae are undulating, tripartite with central dark line (Fig. 13.4), spacing 10 to 13 per 2 mm and 0.02 to 0.1 mm in thickness. Galleries are circular. Syringoporid symbionts existed. In tangential section, pillars are circular and isolated (Fig. 13.5, 13.6).

Laminae appear as concentric circles (Fig. 13.5). Microstructure is compact. *Material*.—One specimen (NHMUK PI H3896) from Lower Devonian of Mitchel’s Quarry, Cave Hill, in Lilydale, Victoria (Australia).

### Remarks

This species was originally grouped with *Amnestostroma*, which is regarded as a junior synonym of *Hermatostromella*. This study therefore reassigned the present species in the latter genus.

Family Idiostromatidae Nicholson, 1886a

Genus *Idiostroma* Winchell, 1867

*Type species*.—*Stromatopora caespitosa* Winchell, 1866.

*Idiostroma roemeri* Nicholson, 1886a Figure 14.1–14.4

1886a *Idiostroma roemeri* Nicholson, p. 99, pl. 9, figs. 6–11.

1952 *Idiostroma roemeri*; Lecompte, p. 316, pl. 66, fig. 3.

1960 *Idiostroma roemeri*; Galloway and Ehlers, p. 67, pl. 4.

2015b *Idiostroma roemeri*; Stearn, p. 797, fig. 405a–c.

### Holotype

Specimen (NHMUK PI P6076) from the Middle Devonian of Hebborn, Germany (Nicholson, 1886a, pl. 9, figs. 6–11).

### Description

The preserved specimens show vertically elongated skeletons with diameter of 3 cm. The skeleton consists of the axial zone and peripheral zone. In longitudinal section, the axial zone is subdivided into short pachysteles and arched laminae (Fig. 14.1). Pachysteles are spool-shaped, confined to interlaminar spaces, regularly superposed, spacing 6 to 8 per 2 mm and 0.05 to 0.15 mm in width. Laminae arch upward, spacing about 5 to 6 per 2 mm and 0.1 to 0.2 mm in thickness. The axial tube is observed, 1 mm in width. Galleries are circular, 0.1 to 0.3 mm in width. The peripheral zone is composed of short pachysteles and parallel laminae (Fig. 14.2). In some areas, skeletons are well differentiated. Pachysteles are spool-shaped, confined to interlaminar spaces, rarely superposed, spacing 5 to 6 per 2 mm and 0.1 to 0.3 mm in width. Laminae are continuous, tripartite with central dark line and surrounding light zones, spacing 4 to 5 per 2 mm and 0.3 to 0.5 mm in thickness. Dissepiments are common in galleries. In the other area, skeletons are densely compact and can hardly be differentiated. In tangential section, pachysteles are spool-shaped, confined to interlaminar spaces, regularly superposed. Laminae are concentric (Fig. 14.3). Galleries are circular or vertically elongated (Fig. 14.4). The axial tube is well developed, 1 mm in width. Microstructure is compact or peripherally vacuolate.

### Material

Four specimens from the Middle Devonian of Hebborn (NHMUK PI P6074, P6075, P6076, P6077), in Germany; and one of Teignmouth (NHMUK PI P6078), in South Devon (UK).

### Remarks

This species is different from the type species *I*. *caespitosum* in larger skeletons and the peripherally vacuolate microstructure.

Order Stromatoporida Stearn, 1980

Family Stromatoporidae Winchell, 1867

Genus *Stromatopora* Goldfuss, 1826

*Type species*.—*Stromatopora concentrica* Goldfuss, 1826.

*Stromatopora* aff. *polaris* (Stearn, 1983) Figure 14.5, 14.6

1937a *Stromatopora concentrica*; Ripper, p. 24, pl. 4, figs. 7, 8.

1983 *Ferestromatopora polaris*; Stearn, p. 551, figs. 5A–D.

1990 *Stromatopora polaris*; Stearn, p. 507, fig. 3.8.

1993 *Stromatopora* aff. *polaris*; Webby et al., p. 158, figs. 5F, 23A–F, 24A.

### Holotype

Specimen (GSC 66987) from the Lower Devonian Blue Fiord Formation of the Ellesmere Island, Arctic Canada (Stearn, 1983, fig. 5A, B). *Description*.—Growth forms are unclear due to limited preservation. The skeleton consists mainly of cassiculate structure. In longitudinal section, cassiculate structure is made up of oblique skeletons, interconnected with short pachysteles and pachystromes to form irregular networks (Fig. 14.5), 0.1 to 0.2 mm in width. Pachysteles are scarce, vertically distributed, confined to inter-pachystromes. Pachystromes are short, locally occurred, spacing 7 to 8 per 2 mm. Allotubes are circular or irregular, with rare dissepiments. Syringoporid symbionts rarely existed, 0.3 to 0.5 mm in width. In tangential section, irregular networks are observed (Fig. 14.6). Allotubes are circular, labyrinth-shaped, or irregular. Microstructure is cellular.

### Material

Three specimens from Lower Devonian of Citadel rocks, Murrindal River (NHMUK PI H3904, H3905) and Commonwealth Quarries (NHMUK PI H3925), Buchan in Victoria (Australia).

### Remarks

The cassiculate structure in this species closely resembles the specimens depicted by Webby et al. (1993). It is different from *S*. *hüpschii* in terms of less long vertical pachysteles and a more common cassiculate structure.

*Stromatopora hüpschii* (Bargatzky, 1881) Figure 15.1, 15.2

1881 *Caunopora hüpschii* Bargatzky, p. 290.

1891 *Stromatopora hüpschii*; Nicholson, p. 176, pl. 10, figs 6, 9; pl. 22, figs. 3–7.

1952 *Stromatopora hüpschii*; Lecompte, p. 268, pl. 52, figs. 1–3.

1961 *Stromatopora hüpschii*; Yavorsky, p. 43, pl. 26, figs. 4–6.

1980 *Stromatopora hüpschii*; Mistiaen, p. 209, pl. 13, figs. 3–6.

1999 *Stromatopora huepschii*; Cook, p. 519, fig. 40 A–F.

2005 *Stromatopora huepschii*; May, p. 194, pl. 22, fig. 1; pl. 30, fig. 1.

2008 *Stromatopora huepschii*; Salerno, p. 91, pl. 14, fig. 1.

### Holotype

Specimen from the Middle Devonian of Büchel, Germany (Lecompte, 1952, pl. 52, fig. 2, 2a,b).

### Description

External forms appear to be laminar. The skeleton is composed of short pachysteles and cassiculate structure. In longitudinal section, pachysteles are commonly vertically discontinuous, spacing 5 to 6 per 2 mm and 0.1 to 0.2 mm in width (Fig. 15.1). Lateral skeletons are short, locally showing cassiculate structures or dots. Allotubes are irregular or vertically elongated, with abundant dissepiments. Branching syringoporid symbionts are also abundant, 0.3 to 0.6 mm in width, spacing 1 per 2 mm. In tangential section, pachysteles form irregular networks, enclosing allotubes (Fig. 15.2).

Microstructure is cellular, 0.02 mm in diameter.

### Material

25 specimens from Middle Devonian of Dartington (NHMUK PI P5843, P5877, P5878, P5887, P5888), Teignmouth (NHMUK PI P5486, P5835, P5836, P5838, P5844, P5886, P5892), in South Devon (UK); Büchel (NHMUK PI P5488, P5878, P5880, P5881), Gerolstein (NHMUK PI P5671, P5882, P5889, P5913, 2) and Hebborn (NHMUK PI P5532), in Germany; and Buchan (NHMUK PI H3907, H3921, H3925), in Victoria (Australia).

### Remarks

This species is characterized by short pachysteles and cassiculate structure. It is distinguished from *Salairella buecheliensis* by less extensive pachysteles or more irregular skeletons.

Genus *Pseudotrupetostroma* Khalfina and Yavorsky, 1971

*Type species*.—*Stromatopora pellucida artyschtensis* Yavorsky, 1955.

*Pseudotrupetostroma buchanense* (Ripper,1937a) Figure 15.3, 15.4

1937a *Hermatostroma episcopale* var. *buchanensis* Ripper, p. 32, pl. 5, figs. 9, 10.

1993 *Pseudotrupetostroma buchanense*; Webby et al., p. 154, fig. 20A.

### Holotype

Lectotype (NMV P 141671-72, Webby et al., 1993) from the Lower Devonian Buchan Caves Limestone near Hicks’s, Murrindal, Victoria, Australia (Webby et al., 1993, fig. 20A).

### Description

The skeleton is composed of short pillars and long laminae. In longitudinal section, pillars are short, confined to interlaminar spaces, rarely superposed (Fig. 15.3), spacing 5 to 6 per 2 mm, 0.1 to 0.2 mm in width.

Laminae are extensive, tripartite with a central dark line and surrounding light structure, spacing 2 to 4 per 2 mm, 0.1 to 0.2 mm in thickness. Galleries are circular or rectangular, with abundant dissepiments. In tangential section, pillars are isolated (Fig. 15.4). The microstructure is coarsely cellular.

### Material

Two specimens (NHMUK PI H3906, H3908) from Lower Devonian of Murrindal, Buchan, Victoria (Australia).

### Remarks

This species was revised to *Pseudotrupetostroma buchanense* by Webby et al. (1993) based on possible cellular microstructure. This study tentatively follows this identification.

*Pseudotrupetostroma porosum* (Yang and Dong, 1979) Figure 15.5, 15.6

1979 *Hermatostroma porosum* Yang and Dong, p. 68, pl. 38, figs. 7, 8.

1982 *Trupetostroma cellulosum*; Dong and Wang, p. 13, pl.5, figs. 1, 2.

1982 *Hermatostroma porosum*; Dong and Wang, p. 22, pl. 13, figs. 1, 2.

2008 *Pseudotrupetostroma cellulosum*; Salerno, p. 92–96, pl. 15, figs. 1–3.

2022 *Pseudotrupetostroma porosum*; Huang et al., p. 722, fig. 8A.

### Holotype

Specimen (NIGP 3312 and 33130) from the Middle Devonian Donggangling Formation of Xiangzhou, Guangxi, China (Yang and Dong, 1979, pl. 38, figs. 7, 8).

### Description

Growth form appears to be laminar. The skeleton locally shows regular networks. In longitudinal section, pillars are vertically long, spacing 7 to 8 per 2 mm, 0.1 to 0.2 mm in width. Laminae are short, passing through several pillars, locally continuous with a central dark line and surrounding cellular structure (Fig. 15.5), spacing 7 to 8 per 2 mm, 0.1 to 0.2 mm in thickness. Syringoporids are common, 0.5 mm in width, with abundant dissepiments. In tangential section, skeletons consist of irregular networks, with rare isolated pillars (Fig. 15.6). The microstructure is cellular.

### Material

One specimen from the Middle Devonian of Teignmouth (NHMUK PI P5933), in South Devon (UK).

### Remarks

The present specimens were previously assigned to *Paralellopora goldfussi*, but it differs from the latter in having regular networks and cellular microstructure.

*Pseudotrupetostroma ripperae* Webby and Zhen 1993 Figure 16.1, 16.2

1937a *Hermatostroma episcopale*; Ripper, p. 29, pl. 5, figs. 7, 8.

1993 *Pseudotrupetostroma ripperae*; Webby and Zhen, p. 340, figs. 7A–F, 8C–D, 9A–B.

1993 *Pseudotrupetostroma ripperae*; Webby et al., p. 152, figs. 19F, 20B–C.

### Holotype

Specimen (SUP 97268) from the Lower Devonian Jesse Limestone of the Limekilns area, New South Wales, Australia (Webby and Zhen, 1993, figs. 7E, F, 8C).

### Description

The skeleton consists of short pillars and long laminae. In longitudinal section, pillars are short, confined to interlaminar spaces, locally superposed, spacing 5 to 7 per 2 mm, 0.1 to 0.2 mm in width. Laminae are laterally extensive, tripartite with a central dark line and surrounding celluar structure (Fig. 16.1), spacing 3 to 4 per 2 mm, 0.1 to 0.2 mm in thickness.

Galleries are circular or irregular, locally with dissepiments. In tangential section, pillars are isolated (Fig. 16.2). Laminae are circular. The microstructure is cellular.

### Material

One specimen (NHMUK PI H3916) from Lower Devonian of Heath’s Quarry, Buchan, in Victoria (Australia).

### Remarks

This species was revised to *Pseudotrupetostroma* due to cellular microstructure. It differs from *P*. *porosum* in short pillars and *P*. *buchanense* in cellular microstructure.

Genus *Neosyringostroma* Kaźmierczak, 1971

*Type species*.—*Hermatostroma logansportense* Galloway and St. Jean, 1957.

*Neosyringostroma* sp. Figure 16.3, 16.4

1891 *Stromatopora beuthii*; Nicholson, p. 183, pl. 5, figs. 12, 13; pl. 23, figs. 8–13; pl. 24, fig. 1.

### Description

External forms are noted as bulbous or domical (Nicholson, 1891). The skeleton is composed mainly of long pillars and short pachystromes. In longitudinal section, pillars are vertically long, spacing 4 to 5 per 2 mm and 0.2 to 0.3 mm in width (Fig. 16.3). Pachystromes are commonly short, confined to inter-pachystele spaces, locally continuous and showing cassiculate structure. Syringoporid symbionts locally occur, 0.2 to 0.3 mm in width. Autotubes are vertically elongated or irregular, with rare dissepiments. In tangential section, pillars are interconnected, forming irregular structures (Fig. 16.4), or locally isolated. Microstructure is cellular or melanospheric.

### Material

19 specimens from the Middle Devonian of Hebborn (NHMUK PI P5528, P5533, P5903), Büchel (NHMUK PI P5530, P5874), Gerolstein (NHMUK PI P5529, P5850, P5869), in Germany; Dartington (NHMUK PI P5904), Bishopsteignton (NHMP5840, P5841, P5905), Teignmouth (NHMUK PI P5839, P5883, P5885, P5907), Torquay (NHMUK PI P5842), and one unknown location (NHMUK PI P5906), in South Devon (UK); also Giessen, 60 km north of Frankfurt, Germany (NHMUK PI P6147).

### Remarks

The original specimens of *Stromatopora beuthii* had been revised as *Hermatostroma beuthii* (Lecompte, 1952). However, specimens illustrated by Nicholson (1891) closely resemble *Neosyringostroma* due to dominant pachysteles and cellular microstructure. The present specimens differ from the type species of *Neosyringostroma* by less cassiculate structure, this study therefore leaves it in open nomenclature.

Family Syringostromellidae Stearn, 1980 Genus *Syringostromella* Nestor, 1966

*Type species*.—*Stromatopora borealis* Nicholson, 1891

*Syringostromella* cf. *labryrinthea* Stearn, 1990 Figure 16.5, 16.6

1937a *Stromatopora concentrica*; Ripper, p. 24, pl. 5, figs. 1, 2.

1937a *Stromatopora hupschii*; Ripper, p. 28, pl. 5, figs. 5, 6.

1990 *Syringostromella labyrinthea*; Stearn, p. 507, figs. 5.1, 5.2, 7.5, 7.6, 8.5.

1993 *Syringostromella* cf. *labryrinthea*; Webby et al., p. 163, figs. 25D–F, 32C.

### Holotype

Specimen (GSC 95779) from the Lower Devonian Stuart Bay Formation, Bathurst Island, Arctic Canada (Stearn, 1990, fig. 5.1, 5.2).

### Description

The skeleton is composed of long pachysteles and short pachystromes. In longitudinal section, pachysteles are vertically long, upward joining or dividing (Fig. 16.5), spacing 4 to 6 per 2 mm and 0.1 to 0.2 mm in width. Pachystromes are short, confined to inter-pachystele spaces. Allotubes are vertically elongated, with rare dissepiments. In tangential section, skeleton is labyrinthine (Fig. 16.6). Microstructure is cellular.

### Material

Two specimens from Lower Devonian of Murrindal (NHMUK PI H3907) and Heath’s Quarry (NHMUK PI H3921), Buchan, in Victoria (Australia).

### Remarks

Specimens with dominant vertically long pachysteles were revised to *Syringostromella* by Stearn (1990). We follow this decision.

*Syringostromella zintchenkovi* (Khalfina, 1961) Figure 17.1, 17.2

**Figure 17.**
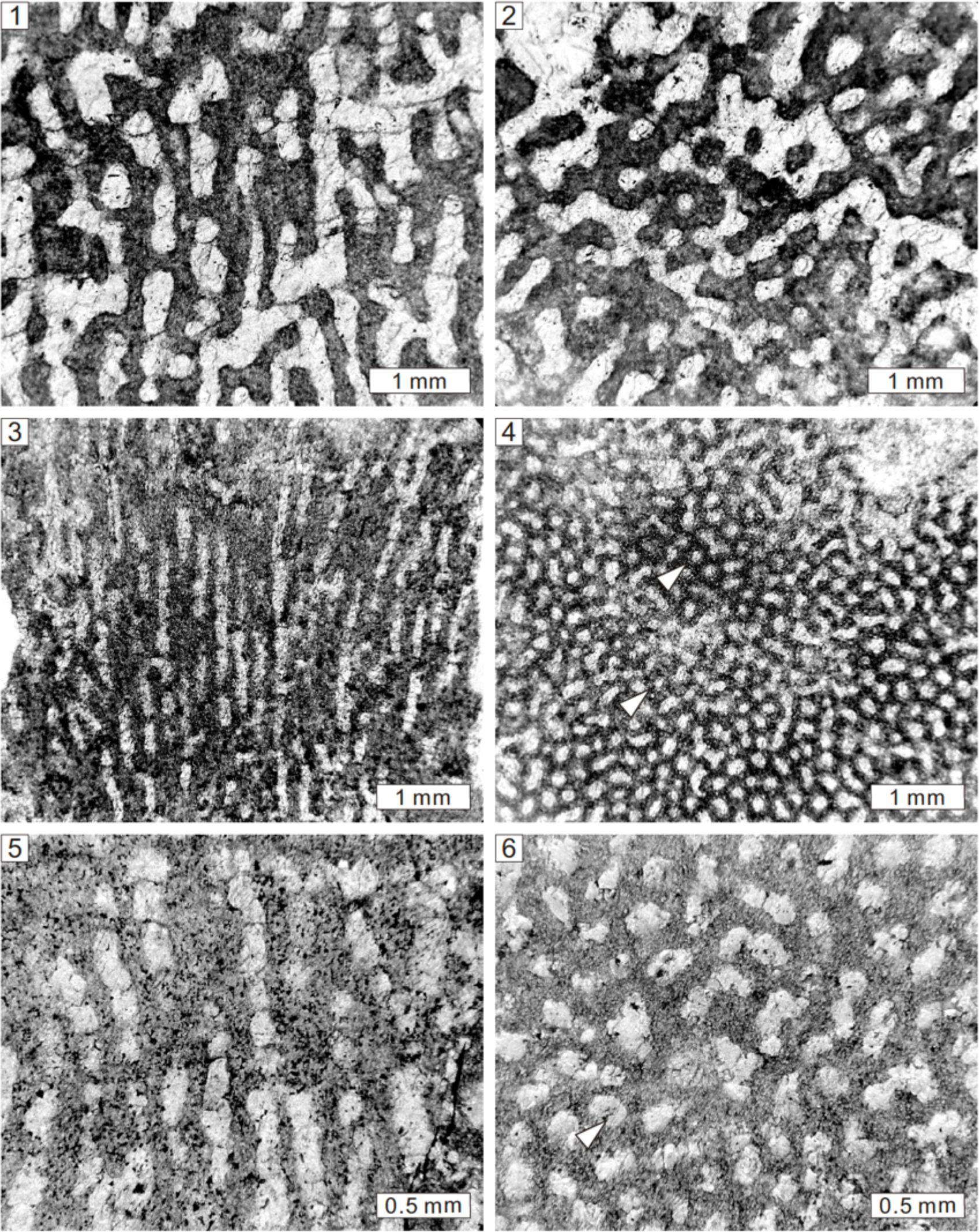
Longitudinal and tangential sections of *Syringostromella zintchenkovi* (Khalfina, 1961) from the Lower Devonian of Cave Hill, Lilydale, Victoria, Australia, NHMUK PI H3894 (**1, 2**), *Salairella buecheliensis* (Bargatzky, 1881) from the Middle Devonian of Büchel, Germany, NHMUK PI P5909 (**3, 4**), *Salairella lilydalensis* (Ripper, 1937c) from Lower Devonian of Cave Hill Quarry, Lilydale, Victoria, Australia, NHMUK PI H3902 (**5, 6**). Note the autotubes (white arrows).

1937c *Stromatopora* aff. *hüpschii*; Ripper, p. 186, pl. 8, figs. 7, 8.

1961 *Stromatopora zintchenkovi*; Khalfina, p. 327, pl. D3, fig. la, b.

1993 *Syringostromella zintchenkovi*; Webby et al., p. 163, figs. 24E–F, 25A–C.

1996 *Syringostromella zintchenkovi*; Prosh and Stearn, p. 34, pl. 15, figs. 1–3.

2012 *Syringostromella zintchenkovi*; May and Rodríguez, p. 228, pl. 1, fig. K, L.

### Holotype

Specimen from the Lower Devonian of Salair, southwestern Siberia, Russia (Khalfina, 1961, pl. D3, fig. la, b).

### Description

The skeleton is composed of long pachysteles and short pachystromes. In longitudinal section, pachysteles are thick, long, discontinuous, vertically joining or dividing (Fig. 17.1), spacing 4 to 6 per 2 mm and 0.1 to 0.3 mm in width. Pachystromes are short, connected to pachysteles. Allotubes are vertically elongated or irregular, with sparse dissepiments. In tangential section, skeleton is labyrinthine (Fig. 17.2). Microstructure is coarsely cellular.

### Material

One specimen from Lower Devonian of Mitchel’s Quarry (NHMUK PI H3894), Cave Hill, Lilydale, Victoria (Australia).

### Remarks

This species is characterized by long pachysteles, and therefore revised to *Syringostromella* by Webby et al. (1993). It differs from *S*. cf. *labryrinthea* in thicker and longer pachysteles.

Genus *Salairella* Khalfina, 1961

*Type species*.—*Salairella multicea* Khalfina, 1961.

*Salairella buecheliensis* (Bargatzky, 1881) Figure 17.3, 17.4

1881 *Caunopora bücheliensis* Bargatzky, p. 290.

1891 *Stromatopora bücheliensis*; Nicholson, p. 186, pl. 10, figs 5–7; pl. 23, figs. 4–7.

1952 *Parallelopora bücheliensis*; Lecompte, p. 290, pl. 50, figs. 3, 4.

1969 *Stromatopora huepschii*; Fischbuch, p. 174, pl. 6, figs. 1–5.

1980 *Stromatopora* cf. *bücheliensis*; Mistiaen, p. 210, pl. 13, figs. 7–9; pl. 14, figs. 1–3.

1985 *Salairella buecheliensis*; Mistiaen, p. 145, pl. 12, figs 10–12; pl. 13, fig. 1.

1992 *Stromatopora hüpschii*; Dong and Song, p. 30, pl. 3, fig. 1a, b.

1999 *Salairella buecheliensis*; May, p. 129, pl. 1, fig. 6.

1999 *Salairella buecheliensis*; Cook, p. 525, fig. 45A–E.

2005 *Salairella buecheliensis*; May, p. 204, pl. 21, figs. 1, 2; pl. 30, figs. 2, 3; pl. 31, fig. 1; pl. 32, fig. 1.

2008 *Salairella buecheliensis*; Salerno, p. 97, pl. 14, fig. 2.

2022 *Salairella buecheliensis*; Huang et al., p. 724, fig. 8B.

### Holotype

Specimen from the Middle Devonian of Büchel, Germany (Lecompte, 1952, pl. 50, fig. 3, 3a).

### Description

Growth forms are laminar, bulbous or domical. The skeleton is composed mainly of long pachysteles and subordinate short pachystromes. In longitudinal section, pachysteles are vertically long (Fig. 17.3), spacing 7 to 9 per 2 mm and 0.1 to 0.2 mm in width. Pachystromes locally occur, short, confined to inter-pachystele spaces. Autotubes are vertically elongated, with sparse or abundant dissepiments. Syringoporid symbionts are locally abundant, with a width of 0.3 to 0.6 mm, spacing 2 per 2 mm. In tangential section, pachysteles are interconnected, forming closed networks (Fig. 17.4), enclosing autotubes and allotubes. Microstructure is cellular, 0.02 mm in diameter.

### Material

33 specimens from the Middle Devonian of Büchel (NHMUK PI P5909, P5910, P5911, P5912, P5922), Gerolstein (NHMUK PI P5670, P5673, P5870), in Germany; Teignmouth (NHMUK PI P5875, P5891, P5914, P5915, P5916, P5924, P5926, P5927, P5928, P5929), Dartington (NHMUK PI P5917, P5918, P5919, P5920, P5921), Bishopsteignton (NHMUK PI H4332), Lummaton Quarry (NHM P5871, P5872), Marychurch (NHMUK PI P5876), in South Devon (UK); Lake Winnipeg (NHMUK PI P5590), in Manitoba (Canada); and Lower Devonian Cave Hill (NHMUK PI H3897, H3901, H3902, H3903), Lilydale, and Loyola (NHMUK PI H3928), in Victoria (Australia).

### Remarks

This species was assigned to *Stromatopora*, but the latter species features by cassiculate structure. It is also different from *Parallelopora* in cellular microstructure.

*Salairella lilydalensis* (Ripper, 1937c) Figure 17.5, 17.6

1937c *Stromatopora lilydalensis* Ripper, p. 189, pl. 9, figs. 1, 2.

1993 *Salairella lilydalensis*; Webby et al., p. 156, figs. 21A–F, 22A–C, 31E.

### Holotype

Lectotype (NMV P141924-25, P141992-93, Webby et al., 1993) from the Lower Devonian of the Lilydale Limestone at Mitchell’s (Cave Hill) Quarry, Lilydale, Victoria, Australia (Webby et al., 1993, fig. 21B, C).

### Description

The skeleton consists mainly of long pachysteles and short pachystromes. In longitudinal section, pachysteles are vertically extensive, spacing 8 to 10 per 2 mm and 0.1 to 0.3 mm in width (Fig. 17.5).

Pachystromes are rare, short, confined to inter-pachystele spaces. Autotubes are vertically elongated, with sparse dissepiments. In tangential section, closed or labyrinthine network is formed by interconnected pachysteles (Fig. 17.6). Autotubes are round. Microstructure is coarsely cellular.

### Material

Four specimens from Lower Devonian of Mitchel’s Quarry (NHMUK PI H3895, H3901) and Cave Hill Quarry (NHMUK PI H3902, H3903), Lilydale, in Victoria (Australia).

### Remarks

This species differs from *S*. *buecheliensis* in closely distributed skeletons.

Order Syringostromatida Bogoyavlenskaya, 1969

Family Coenostromatidae Waagen and Wentzel, 1887

Genus *Coenostroma* Winchell, 1867

*Type species*.—*Stromatopora monticulifera* Winchell, 1866.

*Coenostroma* sp. Figure 18.1, 18.2

**Figure 18.**
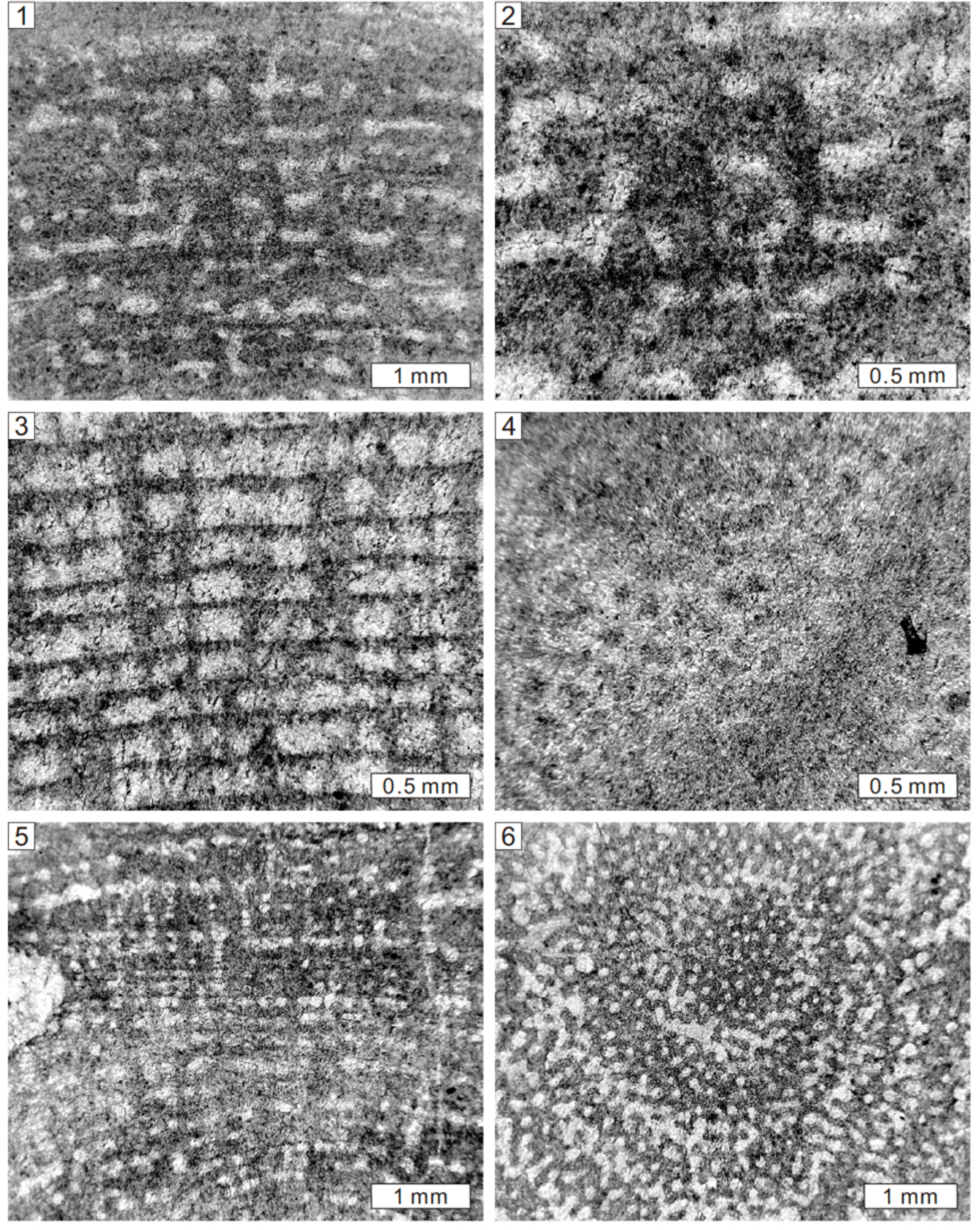
Longitudinal section of *Coenostroma* sp. from Lower Devonian of Buchan, Victoria, Australia, NHMUK PI H3920 (**1, 2**). Longitudinal and tangential sections of *Atopostroma distans* (Ripper, 1937a) from Lower Devonian of Heath’s Quarry, Buchan, Victoria, Australia, NHMUK PI H3915 (**3, 4**), *Columnostroma clathratum* Webby et al., 1993 from Lower Devonian of Mitchel’s Quarry, Cave Hill, Lilydale, Victoria, Australia, NHMUK PI H3893 (**5, 6**).

1937a *Stromatopora concentrica* var. *colliculata*; Ripper, p. 26, pl. 5, figs. 3, 4.

1993 *Coenostroma* sp.; Webby et al., p. 166, figs. 26A–D.

### Description

The skeleton consists of short pachysteles and extensive pachystromes. In longitudinal section, pachysteles are spool-shaped, confined to interlaminar spaces, and locally superposed, spacing 6 to 8 per 2 mm and 0.1 to 0.3 mm in width (Fig. 18.1). Pachystromes are long, thick, spacing 8 to 10 per 2 mm and 0.1 to 0.3 mm in thickness (Fig. 18.2). Galleries are small, circular or laterally elongated, with rare dissepiments. The tangential section is not preserved. Microstructure is unclear.

### Material

Two specimens from Lower Devonian of Buchan (NHMUK PI H3919, H3920), in Victoria (Australia).

### Remarks

Webby et al. (1993) revised this species to *Coenostroma* based on limited skeleton. This study follows this identification.

Genus *Atopostroma* Yang and Dong, 1979

*Type species*.—*Atopostroma tuntouense* Yang and Dong, 1979.

*Atopostroma distans* (Ripper, 1937a) Figure 18.3, 18.4

1937a *Actinostroma stellulatum* var. *distans* Ripper, p. 12, pl. 2, figs. 1, 2.

1993 *Atopostroma distans*; Webby et al., p. 171, figs. 27F, 28A–D.

2008 *Atopostroma distans*; Webby and Zhen, p. 222, fig. 3A–F.

### Holotype

Specimen (NMV P141754-57) from the Lower Devonian Buchan Caves Limestone, Heath’s Quarry, Buchan, Victoria, Australia (Webby et al., 1993, fig. 28A, B).

### Description

The skeleton is composed of thick pillars and thin laminae. In longitudinal section, pillars are rod-like, long, passing through dozens of laminae (Fig. 18.3), spacing 8 to 12 per 2 mm and 0.1 to 0.2 mm in width. Laminae are thin, laterally extensive, locally thickened, spacing 12 to 16 per 2 mm, ranging from 0.02 to 0.1 mm in thickness. Galleries are circular or rectangular. In tangential section, pillars are isolated (Fig. 18.4). Laminae are undulating. Microstructure is unclear.

### Material

One specimen from Lower Devonian of Heath’s Quarry (NHMUK PI H3915), Buchan, Victoria, Australia.

### Remarks

The present species was assigned to *Actinostroma*, but typical colliculi have not been noticed. The regular network formed by long pillars and laminae shows a close relationship with *Atopostroma*. It was therefore grouped with *Atopostroma* by Webby et al. (1993).

Genus *Columnostroma* Bogoyavlenskaya, 1972

*Type species*.—*Coenostroma ristigouchense* Spencer, 1884.

*Columnostroma clathratum* Webby et al., 1993 Figure 18.5, 18.6

1937b *Syringostroma* aff. *niagarense*; Ripper, p. 179, text-fig. 1.

1993 *Columnostroma clathratum*; Webby et al., p. 173, figs. 29A–F, 32F.

### Holotype

Specimen (NMV P141922-23, P 141990-91) from the Lower Devonian Lilydale Limestone at Mitchell’s (Caves Hill) Quarry, Lilydale, Victoria, Australia (Webby et al., 1993, fig. 29A).

### Description

The skeleton consists of long pillars and thin laminae. In longitudinal section, pillars are rod-like, thick, long, spacing 9 to 11 per 2 mm and 0.1 to 0.2 mm in width (Fig. 18.5). Laminae are thin, locally occur, spacing 10 per 2 mm, ranging from 0.02 to 0.1 mm in thickness. Galleries are small, circular. In tangential section, pillars are isolated between laminae (Fig. 18.6). Laminae are cut as closed works. Microstructure is unclear.

### Material

One specimen from Lower Devonian of Mitchel’s Quarry (NHMUK PI H3893), Cave Hill, Lilydale, in Victoria (Australia).

### Remarks

This species was revised to *Columnostroma clathratum* by Webby et al. (1993) based on long pillars. We follow this identification.

Genus *Habrostroma* Fagerstrom, 1982

*Type species*.—*Stromatopora proxilaminata* Fagerstrom, 1961.

*Habrostroma tyersense* Webby et al., 1993 Figure 19.1, 19.2

**Figure 19.**
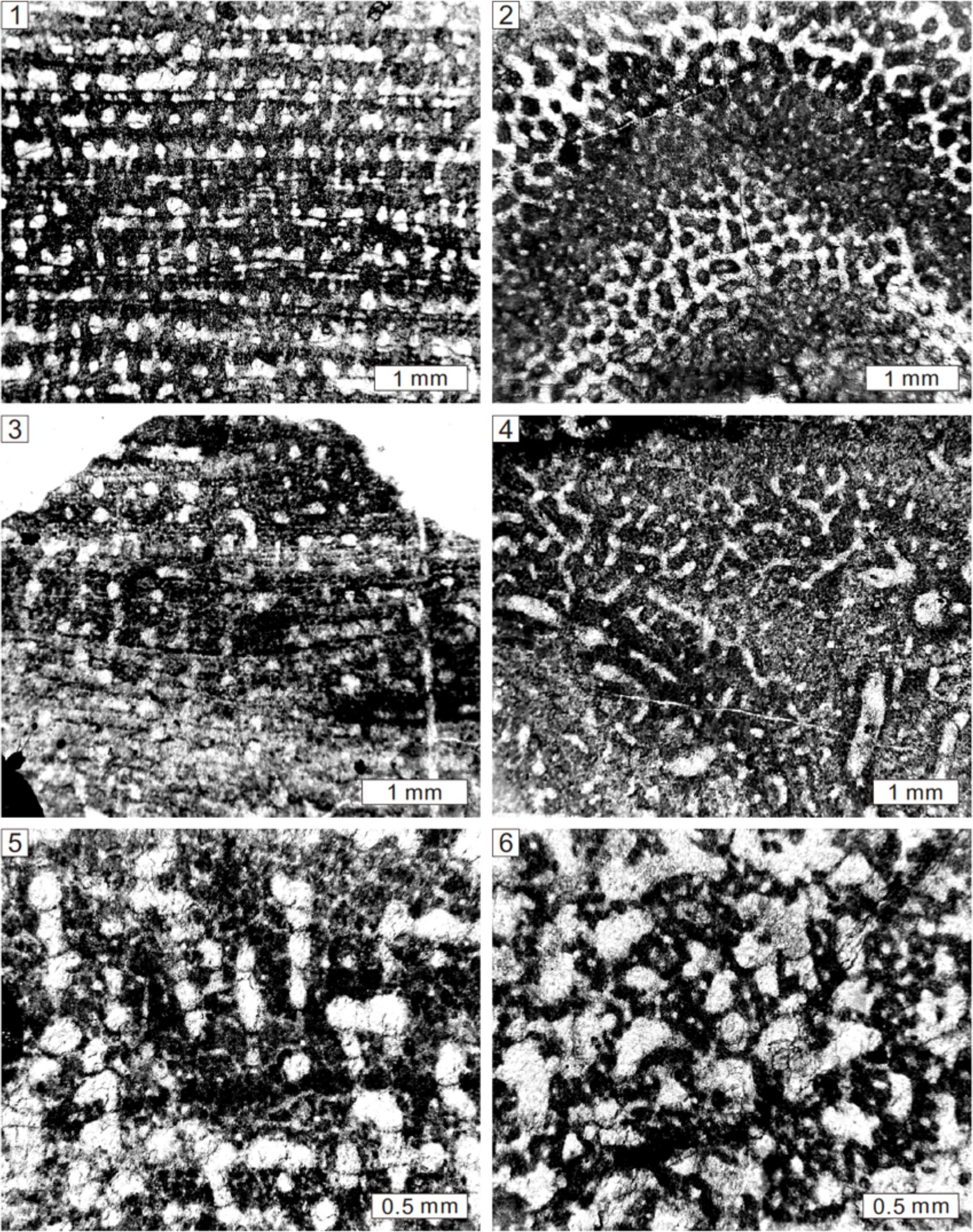
Longitudinal and tangential sections of *Habrostroma tyersense* Webby et al., 1993 from the Lower Devonian of Cave Hill, Lilydale, Victoria, Australia, NHMUK PI H3899 (**1, 2**), *Parallelostroma* sp. from the Middle Devonian of Sötcnich, Germany, NHMUK PI P5631 (**3, 4**), *Paralellopora goldfussi* Bargatzky, 1881 from the Middle Devonian of Teignmouth, South Devon, UK, NHMUK PI P5548 (**5, 6**).

1895 *Syringostroma foveolatum*; Girty, p. 295, pl. 6, figs. 8, 9.

1937c *Stromatopora foveolata*; Ripper, p. 185, text.fig. 2.

1988 *Parallelostroma foveolatum*; Stock, p. 12, figs. 3.5–3.8, 4.1, 4.2.

1993 *Habrostroma tyersense*; Webby et al., p. 168, figs. 26E, F, 27A–E, 32D, E.

### Holotype

Specimen (NMV P136342-43) from the Lower Devonian Coopers Creek Limestone, Tyers Quarry near Tyers, Victoria, Australia (Webby et al., 1993, fig. 26E, F).

### Description

Growth forms are domical to bulbous according to Ripper (1937c). The skeleton is composed of short pillars and long laminae. In longitudinal section, pillars are spool-shaped, confined to interlaminar spaces, locally superposed, spacing 7 to 8 per 2 mm and 0.05 to 0.2 mm in width (Fig. 19.1). Laminae are thick, laterally extensive, common consisting of 2 to 4 microlaminae and micropillars, spacing 6 to 7 per 2 mm, ranging from 0.1 to 0.3 mm in thickness. Allotubes are circular or laterally irregular, with sporadic dissepiments. In tangential section, skeletons are isolated between interlaminar spaces and interconnected in laminae (Fig. 19.2). Microstructure is cellular.

### Material

Two specimens from the Lower Devonian of Cave Hill (NHMUK PI H3899), Lilydale and the Lower Devonian of Rocky Camp (Commonwealth) Quarry (NHMUK PI H3927), Buchan, in Victoria (Australia).

### Remarks

This species was identified as *Stromatopora* by Ripper (1937c), but cassiculate structure has not been observed in it. The occurrence of isolated pillars, multi-microlaminae, and cellular microstructure justify the assignment into *Habrostroma*.

Family Parallelostromatidae Bogoyavlenskaya, 1984

Genus *Parallelostroma* Nestor, 1966

*Type species*.—*Parallelostroma typicum* (Rosen, 1867).

*Parallelostroma* sp. Figure 19.3, 19.4

### Description

Growth forms are unclear due to limited preservation. The internal skeletons consist of regular networks formed by short pachysteles and long pachystromes. In longitudinal section, pachysteles are thick, short, confined to interlaminar spaces, locally superposed, spacing 6 to 7 per 2 mm and 0.1 to 0.5 mm in width (Fig. 19.3). Pachystromes are thick, laterally extensive, locally discontinuous, consisting of several microlaminae and micropillars, spacing 8 to 10 per 2 mm, ranging from 0.1 to 0.2 mm in thickness. Allotubes are circular or irregular, with sparse dissepiments. In tangential section, skeletons are vermicular, with common astrorhizal canals and rarely isolated pillars (Fig. 19.4). Microstructure is orthoreticular.

### Material

Five specimens from the Middle Devonian of Sötenich (NHMUK PI P5631, P5669), Gerolstein (NHMUK PI P5632, P5867, P5873), in Germany.

### Remarks

The specimens were previously identified as *Stromatopora concentrica* by Nicholson (1886a), but the present species differs from the latter in regular networks and orthoreticular microstructure. It shows less development of cassiculate structures, we keep it in open nomenclature.

Genus *Parallelopora* Bargatzky, 1881

*Type species*.—*Parallelopora ostiolata* Bargatzky, 1881.

*Paralellopora goldfussi* Bargatzky, 1881 Figure 19.5, 19.6

1881 *Parallelopora goldfussi* Bargatzky, p. 295.

1891 *Parallelopora goldfussi*; Nicholson, p. 191, pl. 25, figs. 4–9.

1940 *Parallelopora goldfussi*; Chi, p. 298, pl. 4, fig. 2.

1952 *Stromatopora goldfussi*; Lecompte, p. 280, pl. 57, figs. 3–4; pl. 58, fig. 1.

1979 *Parallelopora goldfussi*; Yang and Dong, p. 63, pl. 34, figs. 5, 6.

1992 *Parallelopora goldfussi*; Dong and Song, p. 32, pl. 4, fig. 3a, b.

1999 *Parallelopora goldfussi*; May, p. 129.

### Holotype

Specimen from the Middle Devonian of Hand near Paffrath, Germany (Lecompte, 1952, pl. 57, fig. 3, 3a, b).

### Description

External forms are bulbous or domical (Nicholson, 1891). The skeleton consists of pachysteles and pachystromes. In longitudinal section, pachysteles are vertically long or confined to inter-pachystrome spaces, spacing 9 to 10 per 2 mm and 0.1 to 0.3 mm in width (Fig. 19.5).

Pachystromes locally develop, thin or rarely thick, 0.02 to 0.3 mm in thickness. Autotubes are circular or vertically elongated, with common dissepiments. In tangential section, pachysteles are interconnected, forming closed networks (Fig. 19.6). Branching canals are observed. Microstructure is orthoreticular.

### Material

10 specimens from the Middle Devonian of Dartington (NHMUK PI P5557), Teignmouth (NHMUK PI P5547, P5548, P5551, P5938), in South Devon (UK); and Hebborn (NHMUK PI P5555, P5932), Hand near Paffrath (NHMUK PI P5930), Steinbreche, also near Paffrath (NHMUK PI P5552, P5931), in Germany.

### Remarks

This species was revised into *Stromatopora* (Lecompte, 1952), but cassiculate structure is not observed and the orthoreticular microstructure is different from the cellular microstructure of the latter genus.

*Parallelopora ostiolata* Bargatzky, 1881 Figure 20.1, 20.2

**Figure 20.**
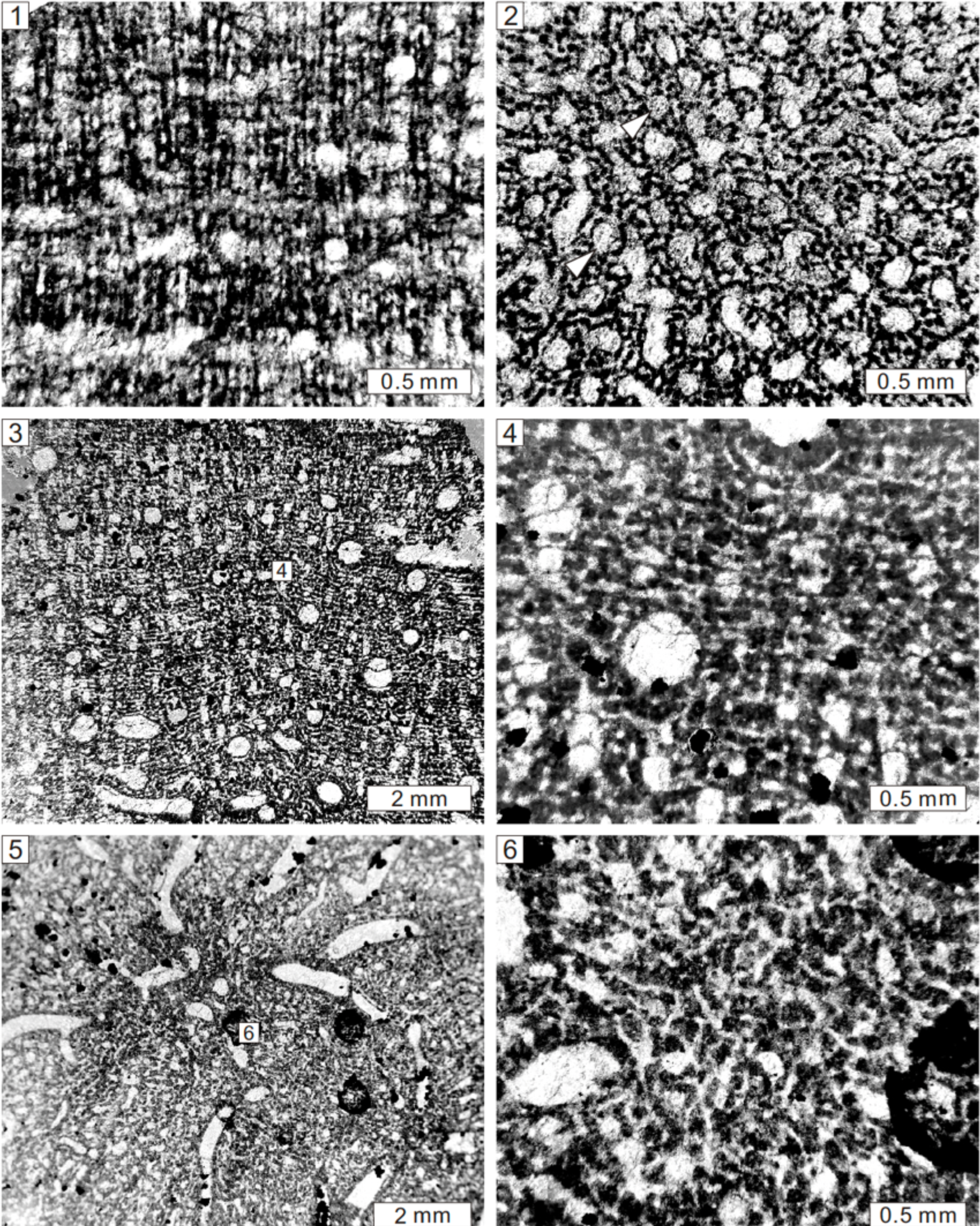
Longitudinal and tangential views of *Parallelopora ostiolata* Bargatzky, 1881 from the Middle Devonian of Büchel, Germany, NHMUK PI P5936 (1, 2), *Parallelopora planulata* (Hall and Whitfield, 1873) from the Devonian of Iowa (USA), NHMUK PI P5801 (3–6). Note the autotubes (white arrows).

1881 *Parallelopora ostiolata* Bargatzky, p. 292.

1886a *Parallelopora ostiolata*; Nicholson, p. 95, pl. 2, figs. 6, 7.

1952 *Parallelopora ostiolata*; Lecompte, p. 292, pl. 51, fig. 3.

1957 *Parallelopora ostiolata*; Galloway and St. Jean, p. 207, pl. 19, fig. 2.

1960 *Parallelopora ostiolata*; Galloway, p. 632, pl. 76, fig. 4.

1979 *Parallelopora ostiolata*; Yang and Dong, p. 62, pl. 33, figs. 7, 8.

### Holotype

Specimen from the Middle Devonian of Büchel, Germany (Lecompte, 1952, pl. 51, fig. 3, 3a–c).

### Description

External form is uncertain due to limited preservation. The skeleton consists mainly of long pachysteles. In longitudinal section, pachysteles are long, vertically extensive, locally joining together, densely packed (Fig. 20.1), spacing 9 to 11 per 2 mm and 0.2 to 0.3 mm in width. Pachystromes rarely develop, short, confined to inter-pachystele spaces. Autotubes are circular or elongate, with sparse dissepiments. In tangential section, pachysteles are interconnected, forming closed networks, enclosing autotubes or allotubes (Fig. 20.2). Microstructure is orthoreticular, with long micropillars and short microcolliculi.

### Material

One specimen from the Middle Devonian of Büchel (NHMUK PI P5936), Germany.

### Remarks

This specimen is the holotype of this species, it differs from other species in closely spaced pachysteles and orthoreticular microstructure.

*Parallelopora planulata* (Hall and Whitfield, 1873) Figure 20.3–20.6

1873 *Caunopora planulata* Hall and Whitfield, p. 228, pl. 9, fig. 2.

1886a *Stromatoporella*? *incrustans*; Nicholson, p. 95, pl. 3, fig. 6.

1936 *Trupetostroma planulatum*; Parks, p. 62, p1. 11, figs. 1–3.

1952 “*Caunopora planulata*”; Lecompte, p. 286, pl. 50, figs. 1, 2.

1957 *Parallelopora planulata*; Galloway and St. Jane, p. 242.

1966 *Parallelopora planulata*; Stearn, p. 106.

1984 *Stromatopora incrustans*; Stock, p. 782, figs. 3E–H, 4.

2008 “*Habrostroma*” *incrustans*; Stock, p. 49.

### Holotype

Specimen (NHMUK P5808) from the Devonian of Iowa, USA.

### Description

Growth forms appear to be laminar. The skeleton is composed mainly of long pachysteles and short pachysteles (Fig. 20.3, 20.4). In longitudinal section, pachysteles are vertically extensive, locally joining together, spacing 7 to 8 per 2 mm and 0.1 to 0.3 mm in width. Pachystromes are irregular distributed, laterally confined, or locally passing through several pachysteles. Autotubes are vertically elongated or circular, with sporadic dissepiments. Astrorhizal canals are cut as circles, 0.3 to 1 mm in width. In tangential section, pachysteles are interconnected, forming closed networks, enclosing autotubes (Fig. 20.5). Astrorhizal canals are common (Fig. 20.3, 20.5). Microstructure is orthoreticular (Fig. 20.4, 20.6).

### Material

Three specimens (NHMUK PI P5799, P5801, P5808) from the Devonian of Iowa (USA).

### Remarks

This species is problematic since it was erected (see Stock, 1984). Given the absence of ring pillars, tripartite pachystromes, cassiculate structures, and isolated pillars, the present specimens differ from *Stromatoporella*, *Trupetostroma*, *Stromatopora*, *Habrostroma*, respectively. It closely resembles *Parallelopora* due to dominant long pachysteles and orthoreticular microstructure.

Family Stachyoditidae Khromykh, 1967

Genus *Stachyodes* Bargatzky, 1881

*Type species*.—*Caunopora verticillata* McCoy, 1850.

*Stachyodes verticillata* (McCoy, 1850) Figure 21.1–21.4

**Figure 21.**
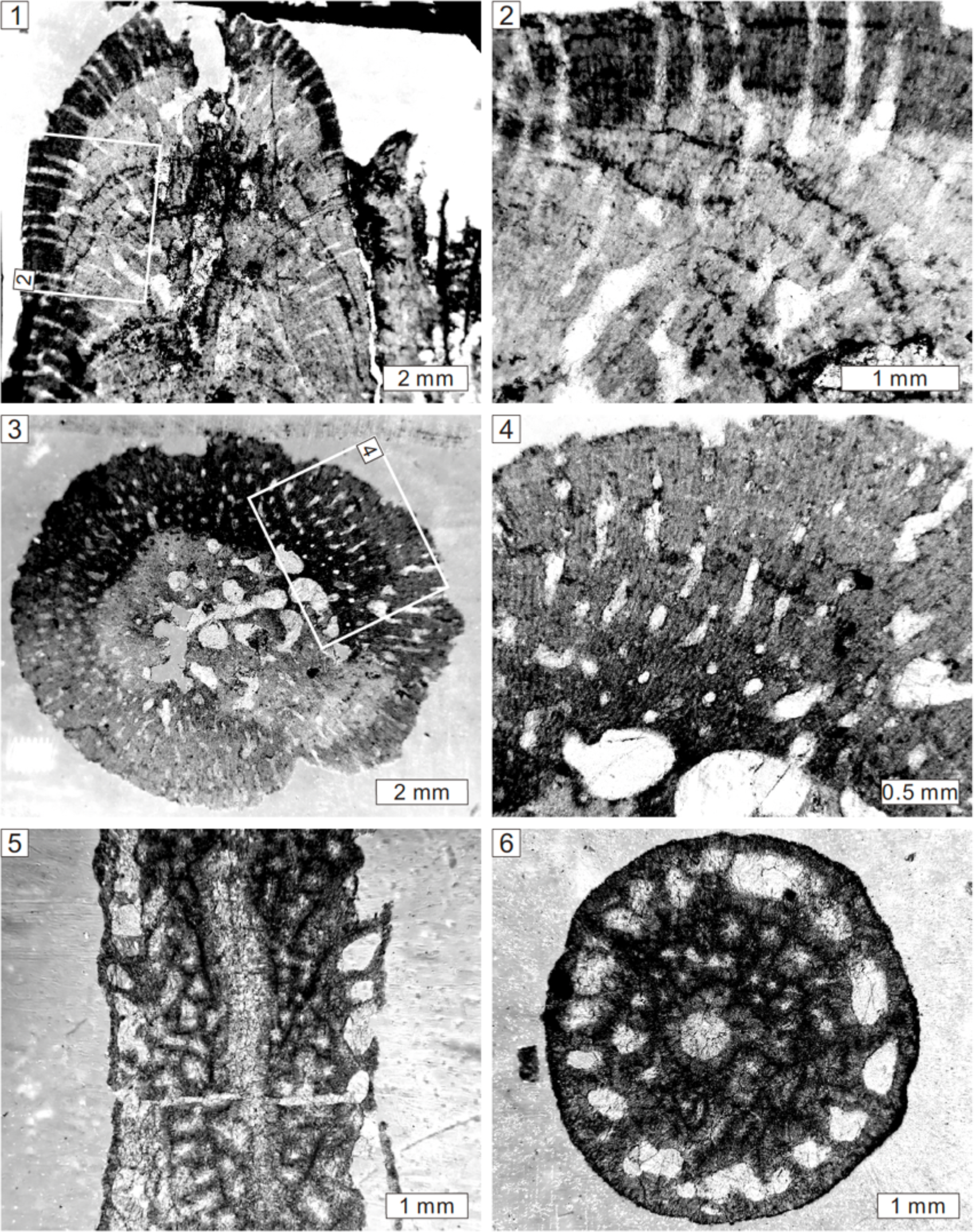
Longitudinal and tangential views of *Stachyodes verticillata* (McCoy, 1850) from the Middle Devonian of Hebborn, Germany, NHMUK PI P6069 (1–4), *Amphipora ramosa* (Phillips, 1841) from the Middle Devonian of Hebborn, Germany, NHMUK PI H6071 (5, 6).

1850 *Caunopora verticillata* McCoy, p. 67, text-figs. a, b.

1886a *Stachyodes verticillata*; Nicholson, p. 107, pl. 8, figs. 9–14.

1892 *Stachyodes verticillata*; Nicholson, p. 221, pl. 29, figs. 1, 2.

1952 *Stachyodes verticillata*; Lecompte, p. 303, pl. 57, figs. 1–3.

1957 *Stachyodes verticillata*; Galloway, p. 444, pl. 34, fig. 10.

1970 *Stachyodes verticillata*; Fischbuch, p. 1080, pl. 149, figs. 1–3.

### Holotype

Specimen (NHMUK P6069) from the Middle Devonian of Hebborn, Germany (Nicholson, 1886a, pl. 8, figs. 9–14).

### Description

External forms are dendroid, but commonly cut as fragmented branches, in diameter of 5 to 10 mm. In longitudinal section, the skeleton comprises thick pachysteles and dark thin pachystromes (Fig. 21.1). In the central zone, skeletons are densely compact, with axial and branching canals 0.2 to 1 mm in diameter. In the peripheral zone, skeletons are differentiated. Pachysteles radiate outward and upward from the central area and end perpendicularly to the surface (Fig. 21.2), 0.2 mm in width. Pachystromes are not well developed, 0.02 mm in thickness. Allotubes are elongate, perpendicular to the surface, 0.1 mm in diameter. In tangential section, closed networks are formed by interconnected pachysteles. The central zone is dominated by axial and branching canals (Fig. 21.3). In the marginal zone, skeleton is featured by long pachysteles (Fig. 21.4). The microstructure is striated.

### Material

Seven specimens from the Middle Devonian of Hebborn (NHMUK PI P6069, P6098, P6099), in Germany; and Middle Devonian of Teignmouth (NHMUK PI P6066, P6067, P6068, P6070) in South Devon (UK).

### Remarks

This species is characterized by densely packed skeletons, well-developed axial and branching canals, thin allotubes, and typical striated microstructure. It is different from *S*. *radiata* in undifferentiated skeletons.

Order Amphiporida Rukhin, 1938

Family Amphiporidae Rukhin, 1938

Genus *Amphipora* Schulz, 1883

*Type species*.—*Caunopora ramosa* Phillips, 1841.

*Amphipora ramosa* (Phillips, 1841) Figure 21.5, 21.6

1841 *Caunopora ramosa* Phillips, p. 19, pl. 8, fig. 22.

**Figure 22.**
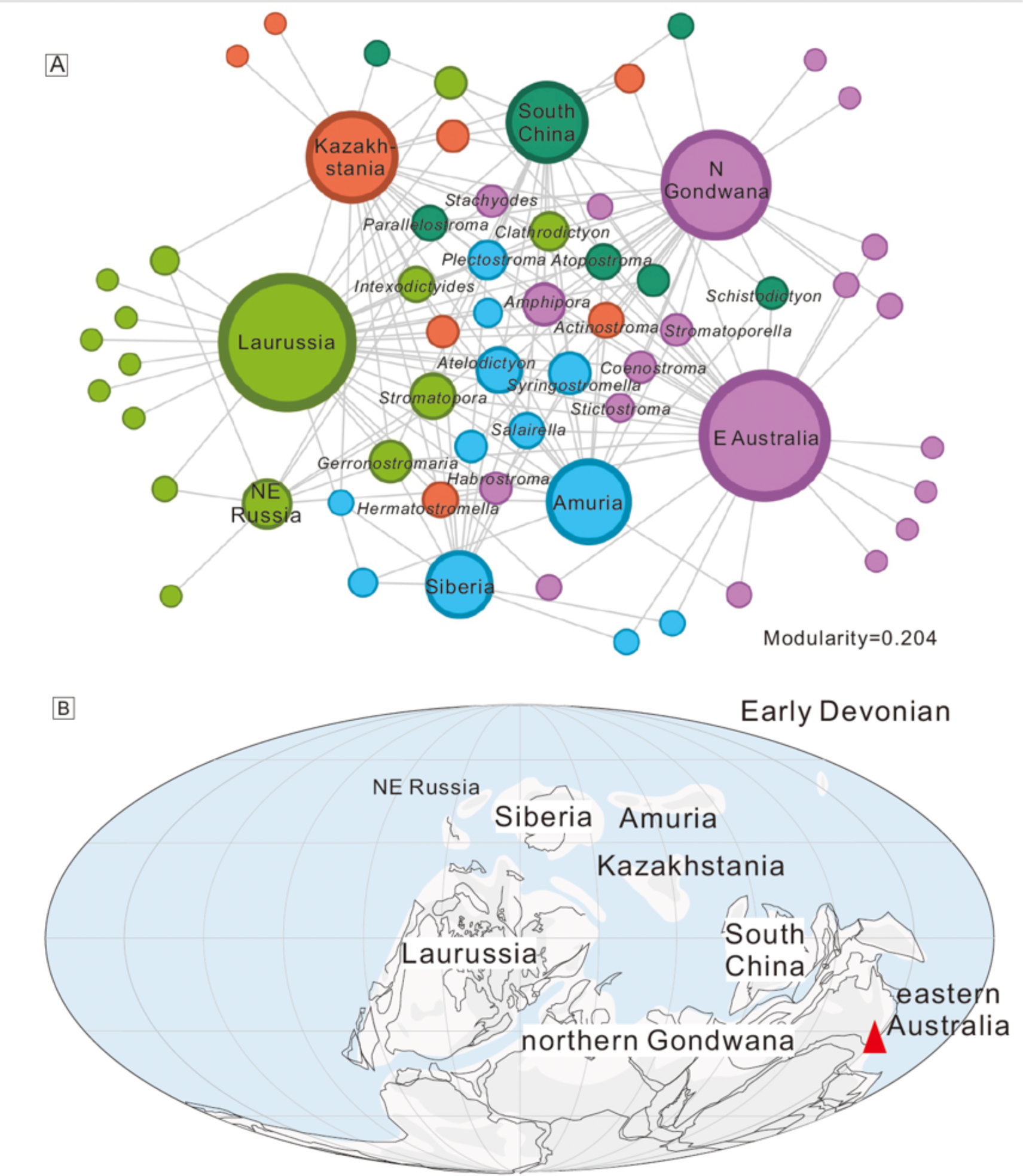
Network analysis of the Early Devonian stromatoporoids around the globe. Note the red triangle represents the location of Early Devonian stromatoporoids used in this study, which are all in Australia. Paleogeographic map follows Golonka (2002, 2007).

**Figure 23.**
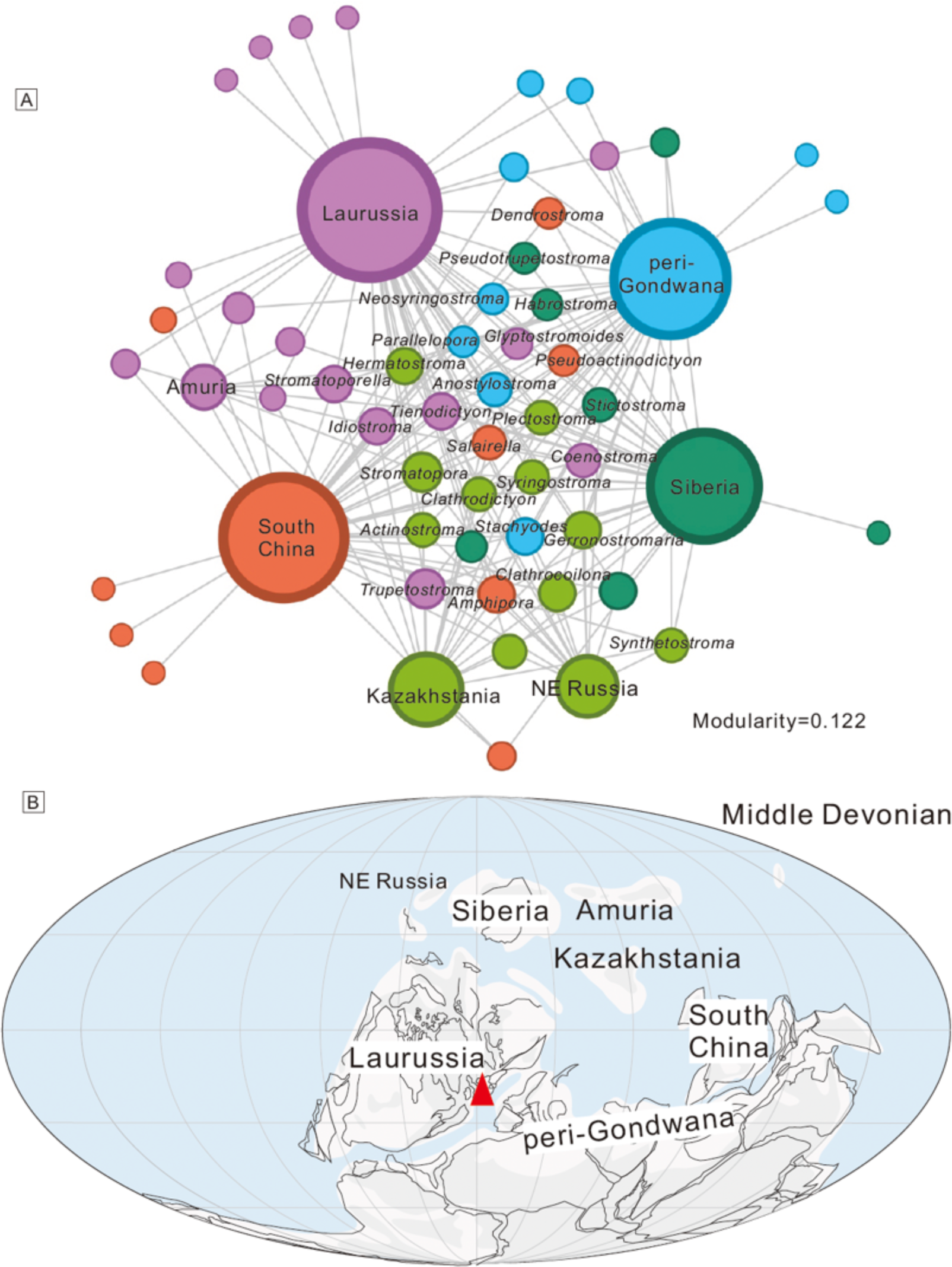
Network analysis of the Middle Devonian stromatoporoids around the globe. Note the red triangle represents the location of Middle Devonian stromatoporoids used in this study. Paleogeographic map follows Golonka (2002, 2007).

1883 *Amphipora ramosa*; Schuelz, p. 90, pl. 22, figs. 5–7.

1886a *Amphipora ramosa*; Nicholson, p. 109, pl. 9, figs. 1–4.

1940 *Amphipora ramosa*; Chi, p. 312, pl. 5, figs. 1, 2.

1952 *Amphipora ramosa*; Lecompte, p. 325, pl. 67, fig. 3; pl. 68, figs. 1–7.

1957 *Amphipora ramosa*; Galloway and St. Jean, p. 233, pl. 23, figs. 2–6.

1961 *Amphipora ramosa*; Stearn, p. 946, pl. 107, figs. 9, 10.

1967 *Amphipora ramosa*; Birkhead, p. 84, pl. 16, fig. 2.

1971 *Amphipora ramosa*; Zukalová, p. 117, pl. 37, fig. 1; pl. 38, fig. 22.

1979 *Amphipora ramosa*; Yang and Dong, p. 79, pl. 43, figs. 7, 8.

1989 *Amphipora ramosa*; Dong et al., p. 278, pl. 16, fig. 3a, b.

1999 *Amphipora ramosa*; Cook, p. 515, fig. 35A–H.

2008 *Amphipora ramosa*; Salerno, p. 104, pl. 2, fig. 4.

### Holotype

Neotype (NHMUK P0308) from the Middle Devonian Chercombe Bridge Limestone, near Newton Abbott, Devon, UK (Stearn, 2015b, figs. 475, 476).

### Description

Growth forms are dendroid, in diameter of 4 mm. The skeleton shows networks formed by oblique pillars and short elements. In longitudinal section, pillars are long, oblique, radiating outward and upward from the skeletal center (Fig. 21.5), 0.1 to 0.3 mm in width. Short elements are also oblique, connecting with pillars and forming irregular networks, 0.1 to 0.2 mm in width. Dark central lines are observed in skeletal elements. Axial canal is conspicuous, in diameter of 0.5 mm. Peripheral sheaths are well developed, 0.5 mm in width, with common occurrence of dissepiments. Skeletal walls are thin, connecting with outside extensions of pillars. Galleries are irregular. In tangential section, central networks are irregular, consisting of oblique pillars and short elements (Fig. 21.6). Axial canal is round. Peripheral sheaths are subdivided by pillars. Microstructure is compact.

### Material

Nine specimens from the Middle Devonian of Teignmouth (NHMUK PI P6072, P6083, P6084, H4776) and Torquay (NHMUK PI H4779) in South Devon, (UK); and Middle Devonian of Herrenstrase (NHMUK PI P5756), Sötenich (NHMUK PI P5758), Hebborn (NHMUK PI P6071, H4778), in Germany.

### Remarks

This species is very common and widely distributed, it is characterized by dendroid skeleton, apparent axial canal, and peripheral sheaths.

## 5. Paleobiogeographical implication of global stromatoporoid distribution and development during the Early to Middle Devonian

Early to Middle Devonian time saw the expansion of stromatoporoids compared with Ordovician and Silurian counterparts. A large number of stromatoporoid species have been erected by global workers, which is too complex to be clearly listed. Therefore, this study focuses mainly on generic level. A total of 55 genera (Early Devonian) and 51 genera (Middle Devonian) of global stromatoporoids are compiled in this study, using a combination of NHM samples and data from publications. The validity of all these genera cannot be fully addressed in this study, therefore we follow the list of genera in the revised Hypercalcified Sponges volume of Treatise on Invertebrate Paleontology (Webby, 2015; Nestor, 2015; Stock, 2015; Stearn, 2015b). Stromatoporoids in seven or eight continents (regions) are analyzed, with a separate focus on the Early Devonian stromatoporoids in eastern Australia.

Network analysis of the global stromatoporoids is carried out and shown in Figs. 22, 23. In the Early Devonian, 15 genera are endemic taxa and confined to one single region, whereas 30 genera are more cosmopolitan and present in three or more regions. The nodes of Kazakhstania, South China. and northern Gondwana are distributed in the upper part of the diagram. And Amuria, northeastern Russia, and Siberia appear to be related in the lower part of the plot. During the Middle Devonian, 10 genera are endemic taxa and occur in a single continent, whereas 37 genera are more cosmopolitan and emerge in three or more regions. Different continents are clearly distributed in the diagram. By using modularity analysis, a grouping pattern is illustrated by colored nodes (Figs. 22, 23). A gradual decrease of modularity value from 0.204 to 0.122 is shown during the Early to Middle Devonian. Compared with the relatively high modularity value from 0.38 to 0.27 of early Silurian brachiopods (Huang et al., 2018), this study therefore implies a relatively close relationship and more cosmopolitan of global stromatoporoid fauna. The plot further shows three groups during Early Devonian, including northern Gondwana-eastern Australia, northeastern Russia-Laurussia, and Amuria-Siberia, possibly indicating a close connectivity of these regions. Another two groups in the Middle Devonian include Laurussia-Amuria and Kazakhstania-northeastern Russia. Although the Devonian marine fauna is traditionally divided into three realms of Malvinokaffric Realm, Eastern Americas Realm, and Old World Realm (Dowding et al., 2022) mainly based on brachiopod and coral fauna, the Early to Middle Devonian stromatoporoid fauna is distributed in lower to middle latitudes and shows relatively weak provincialism in generic level.

Devonian stromatoporoids show a gradual diversification from the Early Devonian Lochkovian Age to Middle Devonian Eifelian Age (Stearn, 2015a). But the comparison between Early Devonian Epoch and Middle Devonian Epoch shows a similar and even higher generic diversity of the former interval. Among the stromatoporoids in the Early Devonian, genera known as *Clathrodictyon*, *Parallelostroma*, *Intexodictyides*, *Schistodictyon*, *Simplexodictyon*, *Syringostromella*, *Petridiostroma*, and *Actinostromella* are holdovers from Silurian (Webby et al., 1993), which may attribute to the relatively high diversity of Early Devonian. And many genera make their first appearance, such as *Atelodictyon*, *Stromatoporella*, *Tubuliporella*, *Stictostroma*, *Pseudotrupetostroma*, *Habrostroma*, *Salairella*, *Parallelopora*, *Coenostroma*, *Glyptostromoides*. The Middle Devonian is characterized by more common Devonian taxa among continents, with a relatively minor role of Silurian holdovers. Representative genera illustrated in this study are known as *Stromatopora*, *Salairella*, *Actinostroma*, *Clathrocoilona*, *Gerronostromaria*, *Hermatostroma*, *Trupetostroma*, *Stachyodes*, *Amphipora*.

The Early Devonian is regarded as a downturn period for stromatoporoid development (May, 2022; Kershaw and Jeon, 2024). Many areas, such as South China, England to Poland, or Spain to Carnic Alps, show a limitation of the abundance of stromatoporoid growth due to sea-level fall, terrigenous input, or possible extreme temperature (May, 2022; Kershaw and Jeon, 2024). Although a contraction has been noticed in this interval, the normal development has not been interrupted by critical events, and the communication between different continents is still moving on, which accounts for the relatively common occurrence of cosmopolitan taxa. The Middle Devonian shows a much closer relationship of global stromatoporoid fauna possibly due to transgression or proper climate, with abundant fossils having been reported especially in regions that previously showed a weak stromatoporoid growth.

## 6. Conclusions

A total of 50 species (belonging to 29 genera) of Devonian stromatoporoids, which were described mainly by Nicholson and now deposited in NHMUK, were systematically redescribed and revised based on recent progress regarding taxonomy. Nicholson’s historical collections show a diverse stromatoporoid fauna in the southeast Devon stromatoporoid assemblage, together with some additional material from North America, Germany and Australia. The material is consistent with Devonian stromatoporoid faunas in the areas of study and other areas of occurrence of stromatoporoids (e.g. S. China, Canada). This Project brings the legacy material of the NHMUK into the modern context of stromatoporoid research, and adds to the global database of Devonian stromatoporoids. Network analysis including data from this study, previous publications, and the Paleobiology Database shows a high cosmopolitanism of global stromatoporoids for both Early and Middle Devonian, with the latter even higher than the former during the proliferation stage of Paleozoic stromatoporoid development.

## Acknowledgements

The authors are indebted to reviewers XX, XX and the journal editor XX for their valuable comments that constructively enhance this manuscript. This study was supported by National Key R&D Program of China (2022YFF0800200). Jiayuan Huang appreciates the financial support of China Scholarship Council (file No.202104910277) during his visit to NHM, London (UK) from 2021 to 2022 in the context of the Covid-19 pandemic.

## Declaration of competing interests

The authors declare none.

## Notes

### Competing Interest Statement

The authors have declared no competing interest.

